# Mu suppression reveals auditory-motor predictions after short musical training

**DOI:** 10.1101/2025.10.06.680771

**Authors:** Oscar Bedford, Alberto Ara, Jérémie Ginzburg, Philippe Albouy, Robert J. Zatorre, Virginia B. Penhune

**Author notes:** Correspondence: Oscar Bedford, Montreal Neurological Institute, McGill University, 3801 University Street, Montréal, Québec, H3A 2B4, Canada.

## Abstract

Auditory-motor coupling is a bidirectional neural mechanism that supports speech and music, with evidence of motor system activation during passive listening to both spoken language and learned melodies. Such activation is anticipatory, occurs in non-musicians, and can be elicited at the single-note level. These findings support the idea that motor activity guides auditory perception by relaying predictive timing information. However, the neural processes underlying this activity are not fully understood. EEG studies in musicians have linked it to mu-band suppression, but the temporal scale and the generalizability to the broader population remain unclear. We recruited 25 non-musicians who learned to play a simple melody on a piano-like keyboard. Before and after training, participants passively listened to the trained melody and control melodies. Offline, EEG data from the motor training were used to create a time-frequency mask with which to identify mu suppression occurring during passive listening. Significant mu suppression emerged before each note only during post-training exposure to the practiced melody. Results suggest that mu suppression occurs at the single-note level following short motor training and is not dependent on prior musical experience. Our findings support the notion that motor activity aids perception by anticipating the unfolding of learned auditory-motor sequences.

---

The perception and production of music and speech depend on the integrated representation of sounds and the actions required to produce them (Lahav et al. 2007; Wilson et al. 2004). Moreover, the coupling between sound and action is thought to rely on bidirectional information flow between auditory and motor networks (Hickok and Poeppel 2007; Arnal and Giraud 2014). Sound-action coupling may be part of innate mechanisms important for learning, or it may be acquired through practice. On the one hand, there is evidence of intrinsic bidirectional interactions between the auditory and motor systems at rest (Bedford et al. 2025) and even in the absence of overt movement (Morillon and Baillet 2017). On the other hand, there is evidence that these interactions are strengthened by both short-and long-term training (Du and Zatorre 2017; Herholz and Zatorre 2012). Regardless, the notion of integrated auditory-motor representations is central to theoretical accounts of sound-action coupling. For instance, the *common coding theory* (Prinz 1990) and the broader *embodied cognition framework* (Hommel 2015) maintain that sound-action integration is rooted in overlapping neural codes for both the planning of an action and the representation of its sensory consequences.

The *embodied music cognition* framework as formulated in Maes et al. (2014) proposes that the brain operates on an internal model of the relationship between the body and the environment, which comprises inverse and forward components (Wolpert et al. 1994). Briefly, inverse models enable the prediction of the physical aspects of motion and space implied in the music (Rizzolatti et al. 2001), whereas forward models allow the prediction of the likely auditory outcome of a planned or executed action (Davidson and Wolpert 2005). Crucially, earlier *embodied music cognition* formulations assumed a passive role for the motor system, relying solely on inverse models to explain the relationship between sounds and actions (Halász and Cunnington 2012). This initial view was supported by findings that listening to sounds can automatically trigger motor responses based on previously established auditory-motor associations—a phenomenon commonly referred to as *motor resonance*—with neurophysiological and behavioural studies demonstrating faster or more consistent motor activation when sounds and actions had been repeatedly paired (Haueisen and Knösche 2001; Rusconi et al. 2006).

However, more recent *embodied music cognition* accounts recognise that sensory predictions tied to planned actions are routinely projected onto upcoming auditory percepts (Maes et al. 2014), and that predictive models are important for motor control (Hommel 2015). Specifically, auditory predictions of one’s own actions are thought to be responsible for helping the system attenuate, facilitate, and disambiguate the processing of the subsequent self-produced sounds (Maes et al. 2014). Thus, in line with *predictive coding* accounts (Friston et al. 2010), one of the roles of the motor system must be to generate and maintain forward models—whenever auditory-motor associations are available—to streamline auditory processing. For instance, the motor system is now believed to provide top-down predictions about the timing of upcoming auditory events (Patel and Iversen 2014), a notion that fits into the *predictive timing* framework proposed by Arnal and Giraud (2012).

Evidence for joint auditory-motor representations can be found in neuroimaging and neurostimulation studies showing enhanced activation of motor regions when people listen to melodies they know how to play (Bangert et al. 2006; D’Ausilio et al. 2006; Baumann et al. 2007; Lahav et al. 2007; Lappe et al. 2008; Chen et al. 2012; Herholz et al. 2016). However, none of these studies provide direct evidence that predictable melodic sequences can automatically cue specific upcoming actions. In other words, it is unclear whether this motor activity reflects predictive processes indicative of forward modelling or reactive inverse modelling processes, such as motor resonance. This is because effector/note level anticipatory motor activity can only be reliably assessed at the single-note timescale, as indicated by Novembre and Keller (2014). Therefore, in a previous study from our lab, non-musicians were trained to play a simple melody and measured motor-evoked potentials (MEPs) to show that hearing the tones of the learned melody automatically activated corresponding motor representations of the associated finger muscles *in advance* of the specific tones (Stephan et al. 2018). This finding is highly suggestive of forward modelling processes, but the MEP is only an indirect measure of motor preparation. Therefore, in the current experiment we used EEG to inspect preparatory motor activity by measuring *mu suppression*—a candidate neural index of forward modelling.

The mu rhythm is a subcomponent of the alpha band typically found between 9-13 Hz (Pineda 2005) that is thought to reflect the activity of large groups of pyramidal neurons in M1 (Niedermeyer 1997). This rhythm is characterised by synchronized neural firing at rest, but it desynchronizes quickly in anticipation of voluntary movement, producing a pattern of reduced amplitude known as mu suppression (Pineda 2005). Another context in which mu suppression has been observed is during the observation of action by others, which inspired mirror neuron (Rizzolatti and Craighero 2004) and echo neuron (Keysers and Gazzola 2010) theories, both of which have been implicated in music perception (Overy and Molnar-Szakacs 2009). However, mu suppression is not limited to anticipating active self-movement or when observing movement by others; it also emerges during passive listening to sounds produced by familiar actions, lateralizing to the contralateral hemisphere at effector-specific locations within sensorimotor cortex (Pineda et al. 2013). This observation suggests that mu suppression may index predictive action-perception coupling more broadly (Pineda 2005).

For instance, Wu et al. (2016) reported mu suppression in expert pianists during passive listening to a practiced melody, but not to unfamiliar melodies. This effect preceded the onset of each practiced melody presentation, suggesting that mu suppression may indeed reflect covert motor activity that supports forward modelling. However, because mu suppression was only measured per melody and not in relation to each note/finger movement in the melody, this study still could not demonstrate effector-specific predictions driven by the learned sequence rather than general expectancy (Novembre and Keller 2014). Moreover, because only musicians were tested, the study could not determine whether the observed mu suppression was the result of global predictions contingent on prior musical training or whether it was directly linked to learned auditory-motor associations.

In light of these gaps, in the current study we used the same melody learning task as in the previous MEP study, combined with EEG, to measure possible changes in mu suppression following short-term training in non-musicians. In this task, participants passively listened to learned and unlearned melodies before and after 15 blocks of training on the learned melody. We then compared mu activity preceding each note, with the hypothesis that mu suppression for the learned melody would be greater than for unlearned melodies before each note, demonstrating that a predictive auditory-motor representation or forward model had been established through learning (Stephan et al. 2018; Maes et al. 2014). To objectively identify the likely time-frequency distribution of mu suppression during the passive listening task, we used the EEG data collected during active motor training to derive a time-frequency mask of all frequency bands engaged by the task. We then applied this mask to the passive listening data, which allowed us to assess the pattern of change across all active frequency bands, as well as to test whether any observed suppression effects would be specific to the mu frequency band.

## Materials and methods

### Participants

We recruited 27, healthy, right-handed non-musicians. Two participants were excluded due to poor EEG data quality, resulting in a final sample of 25 participants (14 females) aged between 18 and 35 years (mean = 22.6 ± 4.5 years). Consistent with previous studies, all participants had less than 3 years of lifetime musical training and/or experience and reported no active music-making within the 3 years prior to the study (Slater and Kraus, 2016). Exclusion criteria included left-handedness and a history of psychiatric or neurological disorders. All participants provided written informed consent at the start of the experimental session. The study protocol adhered to the principles of the Declaration of Helsinki (World Medical Association 2001) and was approved by the by the Institutional Review Board of the Montreal Neurological Institute (IRB Study Number A02-B15-22B / 22-01-058).

### Task and stimuli

The goal of the experiment was to assess the degree of mu suppression during passive listening to a trained melody. EEG was recorded throughout the experiment, which featured three conditions summarized in Figure 1A: pre-training passive listening (PRE); melody playback training (TRAIN); and post-training passive listening (POST). In the PRE and POST conditions participants heard two types of melodies: 1) the trained melody (Trained); and 2) untrained melodies (Untrained). In the TRAIN condition participants either listened to and played back to blocks containing repetitions of the Trained melody or catch blocks containing repetitions of the reversed Trained melody (Reversed).

**Figure 1.**
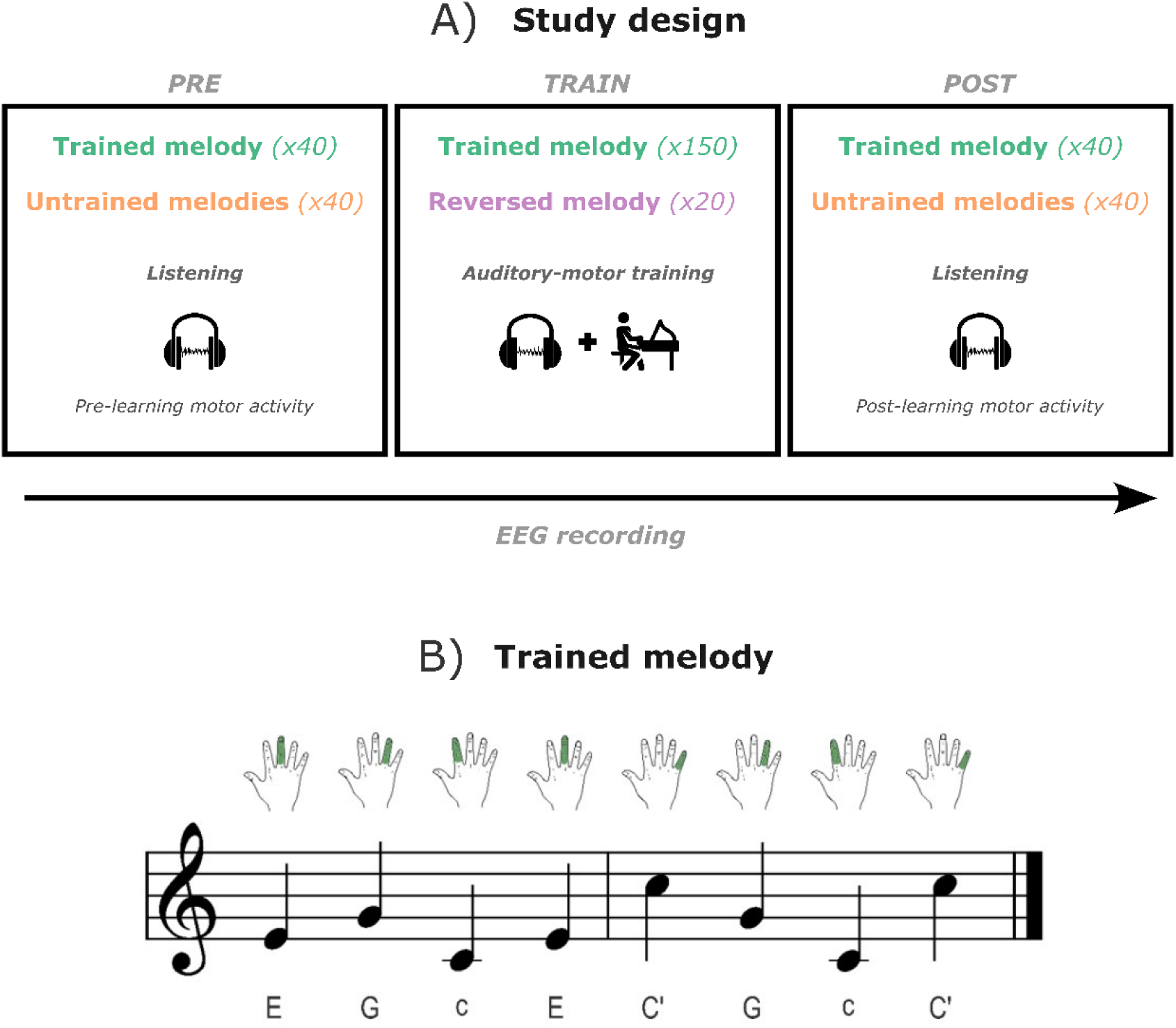
Experimental paradigm. **A)** Study design. **PRE (panel 1)**: subjects passively listened to one block of 40 repetitions of the Trained melody and one block of 40 Untrained melodies. **TRAIN (panel 2)**: subjects listened to and played back to 15 blocks of 10 repetitions of the Trained melody and two catch blocks of 10 repetitions of the Reversed melody. **POST (panel 3)**: identical to PRE. **B)** Trained melody. The arrangement is based on a C-major arpeggio. Each note is associated with one digit of the right hand. The sequence has isochronous timing.

41 unique 8-note melodies (1 trained melody and 40 untrained melodies) were created using the four notes in a C-major arpeggio that corresponded to the four digits of the right hand when played back on a purpose-built piano-like keyboard (C4 (259 Hz) = index; E4 (329 Hz) = middle; G4 (389 Hz) = ring; C5 (531 Hz) = pinkie). Melodies were 12 seconds long, with each note lasting 600 milliseconds, followed by an inter-stimulus interval of 900 milliseconds, resulting in an onset-to-onset asynchrony of 1500 milliseconds. Melodies were constructed such that each note appeared twice and no notes were repeated sequentially (Stephan et al. 2018). The Trained melody is depicted in Figure 1B. All sounds were delivered via E-A-RTONE 3A foam-tipped insert-type earbuds (E-A-R® Auditory Systems). Sounds were synthesized with Adobe® Audition v.3.0 (Adobe Systems Incorporated 2007). All musical notes were rendered using a piano timber, and every melody presentation in the TRAIN condition was cued by a woodblock sound.

### PRE and POST passive listening conditions

During the PRE and POST passive listening conditions, participants listened to one block of 40 repetitions of the Trained melody and one block of 40 unique Untrained melodies (12.7 minutes per block). As an attentional control, seven catch melodies were randomly presented across the two blocks of each passive listening condition (PRE and POST), in which the last note was mistuned (C#5 [554 Hz] instead of the natural C5 [531 Hz]). Participants were instructed to silently count these mistuned notes and report the total at the end of each block. The presentation order of the Trained and Untrained blocks was held across passive listening conditions PRE and POST but counter-balanced between participants. Inter-trial intervals (ITIs) for melodies presented in the passive listening blocks varied randomly between a 3.3 and 3.7 second range (in 0.1 second steps) to prevent anticipation of melody onsets.

### Melody playback training (TRAIN)

The TRAIN condition consisted of 15 blocks (10 trials each) of practice on the Trained melody and 2 catch blocks (10 trials each) of practice on the Reversed melody, for a total of 17 blocks (∼50 minutes of training). Reversed melody blocks appeared in the 3^rd^ and 15^th^ block positions. The ITI within all blocks was fixed at 3.5 seconds. On each trial, participants heard a given melody and were asked to reproduce the notes by pressing four keys on a purpose-built piano-like keyboard. Notes produced by participants’ key presses were played back at an octave below the original melody to provide distinguishable auditory feedback. Participants were instructed to synchronize their responses as closely as possible to each melody presentation, and no additional instructions or performance feedback were provided.

Each keypress activated a switch on the custom keyboard that sent transistor-transistor logic pulses via a parallel port connected to a Beaglebone® Black microcomputer equipped with a Bela real-time module (Thangaraj 2016), based on the system introduced by Zappi and McPherson (2014). These pulses were received by the EEG recording computer with minimal delay and low jitter, and were used as precise event markers for keypress timing.

### Behavioural data analysis

Individual performance in the motor training task (TRAIN) was evaluated for each of the 17 blocks using two metrics: the percentage of correct key presses (accuracy) and the mean absolute timing difference between key presses and target note onsets (reaction time). A key press was considered accurate if it matched the pitch of the intended note within a ±300 millisecond window and was executed without overlapping with other key presses. Reaction times (RTs) were computed only for correct key presses. In cases where individual RTs were missing in a trial within a block, values were imputed using the mean RT for that block’s trial, calculated across all other participants.

### EEG data collection

EEG recordings were collected using a 64-channel active electrode cap (actiCAP), a unipolar EEG amplifier (BrainAmp), and dedicated recording software (BrainVision Recorder, Version 1.20.0801), all from the same manufacturer (Brain Products GmbH 2019). Data were sampled at 1 kHz. The ground electrode was left at the default FPz location, and the reference electrode was placed on the right canthus. This placement allowed us to monitor eye movements and thus encourage participants to blink as little as possible and in a manner uncorrelated with task structure. Any potential visual confounds introduced by this choice of reference placement were eliminated using standard re-referencing offline (Yao, 2019; Hu et al., 2019). For the ground channel, the average kΩ value per participant was 2.5 (SD: 2.4; Median: 2; Range: 0.7 to 10.5 kΩ). For the reference channel, the average kΩ value per participant was 1.1 (SD: 1.3; Median: 0.5; Range: 0 to 5 kΩ). Cap electrodes were arranged at 60 standard positions according to the international 10-10 system (Chatrian et al. 1988).

Electrodes corresponding to channels FT9 and FT10 were placed on the left outer canthus and right cheek, respectively, to capture horizontal and vertical electrooculography (EOG) signals used to remove blinks and saccadic eye movements at preprocessing. Electrodes corresponding to channels TP9 and TP10 were placed on the left and right mastoids, respectively, and served as re-reference sites for offline data preprocessing. For TP9, the average kΩ value per participant was 2.8 (SD: 2.6; Median: 2.3; Range 0.8 to 12.1 kΩ). For TP10, the average kΩ value per participant was 3.1 (SD: 2.7; Median: 2.4; Range: 0 to 11.7 kΩ). Throughout the EEG recording, participants were instructed to maintain their gaze on an onscreen central fixation cross. Impedance levels were checked at the end of each passive listening block in the PRE and POST conditions, as well as midway through the TRAIN condition, and were continuously maintained below 10 kΩ. On average, each participant underwent 5.7 impedance checks during the course of the experiment (SD = 0.9; Median = 6; Range: 2–6 checks). During passive listening, participants were instructed to remain as still as possible, and their right hand was monitored to ensure that no movements contaminated the EEG recording.

### EEG pre-processing

EEG data were pre-processed using a combination of Brainstorm (Tadel et al. 2011) custom scripts in MATLAB R2021a (The MathWorks Inc. 2022), following standard preprocessing and artifact removal steps: EEG signals were re-referenced to bilateral mastoids using a simple linear transform (Tadel et al., 2019), notch-filtered at 60 Hz, and bandpass-filtered between 1-100 Hz. Noisy segments and channels were identified and removed through visual inspection (Gross et al. 2013). To assist in removing artifacts and bad channels, power spectrum density (PSD) plots were generated at successive stages of pre-processing. Across all participants, a total of 21 faulty channels were interpolated across 10 participants, with no more than 4 interpolations for any given participant (mean = 2.1 ± 1.2 channels). In all cases, interpolation was applied upon observing that successive stages of pre-processing had not resolved abnormally overpowered activity in the frequencies contained within our 2-49.5 Hz spectrum. Interpolation was chosen instead of channel removal because preserving continuous scalp coverage was essential for conducting our group-level, cluster-based permutation analysis in sensor space (Courellis et al. 2016).

Repetitive eye blink artifacts were removed for all subjects, and saccade artifacts were removed in three participants, using Brainstorm’s built-in signal space projection (SSP) method (Tesche et al. 1995). SSP was chosen instead of removing trials with eye artifacts because this procedure allows for the attenuation of systematic ocular artifacts without having to exclude data (Uusitalo and Ilmoniemi 1997). The first SSP projector was removed from all participants, and it explained an average 43.3% of the variance (SD: 13%; Median: 45%; Range: 12-66%). Only 7/25 participants required removal of a second projector, which explained an average 16.6% of the variance (SD: 4.7%; Median: 17%; Range: 10-22%). Projectors were assessed from multiple angles before removal: (i) projector timeseries data were compared with EOG timeseries data to ensure that blink shapes closely matched. Also, (ii) projector topographies were inspected to ensure that they featured anterior scalp distributions indicative of ocular activity. Lastly, (iii) projector candidates with topographies that included central channel locations were ruled out to avoid distorting motor signals.

### EEG data analysis

#### Time-frequency analysis

After preprocessing, the time-domain EEG data from the four passive listening conditions were downsampled to 200 Hz and epoched from −2000 to 2000 milliseconds relative to each individual note’s onset. Within each ±2000 millisecond time-domain epoch, the segment ranging from −1245 to 1245 milliseconds was used to baseline-correct the ±2000 millisecond time window in the time domain. This step was applied prior to time-frequency decomposition as a standard de-meaning procedure (Makeig et al., 2004). In addition, the average event-related potential for each participant and listening block was calculated and removed from each time-domain using standard pointwise subtraction (sample-by-sample). This step was implemented to ensure that effects preceding note onset reflected induced (non-phase-locked) rather than evoked (phase-locked) responses, under the notion that any activity occurring prior to note onset can only be induced (Cohen 2014). All in all, this first stage preceding time-frequency decomposition yielded 320 time-domain epochs of ±2000 milliseconds each per participant and listening block (40 melodies × 4 unique notes per melody × 2 unique note repetitions per melody).

Each time-domain epoch was then separately convolved over a linearly scaled frequency range of 2-49.5 Hz (step size: 0.5 Hz) using a family of Gaussian-tapered scaling Morlet wavelets with the mother wavelet centred at 1 Hz and an initial full-width at half maximum of 3 seconds. This approach yielded time-frequency power estimates, which were subsequently normalized using Brainstorm’s event-related spectral perturbation (ERSP) baseline correction method (Tadel et al. 2011). The time-frequency baseline window was taken from within the previously de-meaned ±1245 millisecond segment of interest, itself contained within the full ±2000 millisecond time-frequency epoch. Namely, the 200 millisecond-long time-frequency baseline period was established from−850 to −650 milliseconds relative to note onset. This corresponds to the midpoint (750 milliseconds) ±100 milliseconds between successive notes (IOI/2), where motor-related brain activity was expected to be minimal. The final time-frequency analysis window on which all further statistics were performed was restricted to the segment ranging from−650 to −100 milliseconds in time-frequency space, which also falls within the previously de-meaned ±1245 milliseconds segment of interest. This analysis window was chosen as the most likely to contain relevant preparatory auditory-motor activity (Shibasaki and Hallet 2006). The window was truncated −100 milliseconds prior to tone onset to avoid contamination from auditory (Ross et al. 2022a) and movement activity (Frølich et al. 2018) typically present close to note onset.

The 320 time-frequency plots we obtained at this stage were averaged within each passive listening block for each participant, yielding four averaged time-frequency plots per participant comprising the mean activity across all four fingers. These final plots contained baseline-corrected power estimate values across 64 channels, 97 frequency bins spanning 2 to 49.5 Hz, and 800 timepoints sampled at 200 Hz. Only the 110 baseline-corrected timepoints corresponding to the −650 to −100ms time window were retained for further analysis.

#### Time-frequency mask

To examine changes in EEG activity during listening in the PRE and POST conditions we wanted to independently identify the predicted distribution of possible effects across channels, time and frequency. To this end, we computed a group-level, time-frequency mask based on correctly performed trials of individual key presses in the TRAIN condition, across all frequency bands within the 2-49.5 Hz range, in the time window expected to be associated with response preparation (−650 to −100 milliseconds before note onset). The purpose of this mask was to identify channels, frequencies, and timepoints most likely to exhibit mu suppression during post-training passive listening to the Trained melody. In other words, our mask was defined from independent data to avoid circularity (Kriegeskorte et al. 2009), in line with standard practice and recommendations (Maris and Oostenvelt 2007).

For each participant, we selected the four blocks from blocks 9-17 (excluding catch block 15) of the TRAIN condition with the highest average accuracy scores. From these, a total of 320 time-domain epochs were extracted per participant (10 melodies × 8 notes per melody × 4 Trained melody blocks), using the same epoching, de-meaning and ERP subtraction procedures described above. Next, time-frequency decomposition was performed on each epoch using the same time-frequency parameters and ERSP baseline-correction method described above. Thus, each resulting plot contained baseline-corrected power estimate values across 64 channels, 97 frequency bins spanning 2 to 49.5 Hz, and 800 timepoints sampled at 200 Hz. Consistent with the passive listening conditions, only the 110 timepoints corresponding to the −650 to −100 millisecond time window of interest were retained. All 320 time-frequency plots were averaged at the participant level, yielding one average time-frequency plot per participant comprising the mean activity across all four fingers, for a total of 25 participant-level plots. These 25 active motor training time-frequency plots were compared against zero using a one-sample, left-tailed *t*-test.

The resulting *t*-map was submitted to a cluster-based permutation test implemented via FieldTrip’s native `ft_freqstatistics` function within Brainstorm (Oostenveld et al. 2011; Tadel et al. 2011). To control for multiple comparisons, the test was configured to run 1000 Monte Carlo permutations (Candia-Rivera and Valenza 2022; Pernet et al. 2015). The significance threshold for both the *t*-test and the permutation test was set at the standard alpha level of.05 (*p* =.05). No averaging was performed across dimensions. Regions of interest (ROIs) were allowed to vary freely across time and frequency, provided that activity maintained contiguity within a cluster, defined as there being a minimum of one time-frequency pixel connected across a minimum of two channels.

The resulting cluster-corrected *t*-map was thresholded to retain only pixels with an associated *p*-value less than.05. This thresholded t-map served as a binary three-dimensional mask (channels, frequencies, timepoints), whereby all surviving pixels were weighted equally, regardless of their associated *t*-statistic value, before being transferred to the time-frequency plots derived from the passive data.

The significant mask cluster was divided into three frequency bands: mu (9-16.5 Hz), beta (16.5-33.5 Hz), and gamma (33.5-49.5 Hz). The beta-gamma division was determined based on prior literature, as significant activity between 17 and 49.5 Hz was spectrally contiguous across all 60 channels. In contrast, the mu-band boundaries were determined empirically: among the 33 channels showing significant activity between 9 and 16.5 Hz, 29 displayed mu-band activity that was clearly separated from beta-band activity by several frequency bins containing no significant pixels (see Fig. 3B, channel C3, for an example). In the remaining four channels (C5, CP5, T7, TP7), a small amount of intervening activity bridged the mu and beta ranges. This bridging likely caused pixels in the mu range to be included in the same statistical cluster as those in the beta-gamma range. However, the intervening activity was temporally restricted, and the overall time-frequency profile still matched the isolated mu-band patches observed in the other 29 channels (see Supplementary Fig. 2). Thus, we believe our mu-beta partition to be sufficiently grounded.

**Figure 2.**
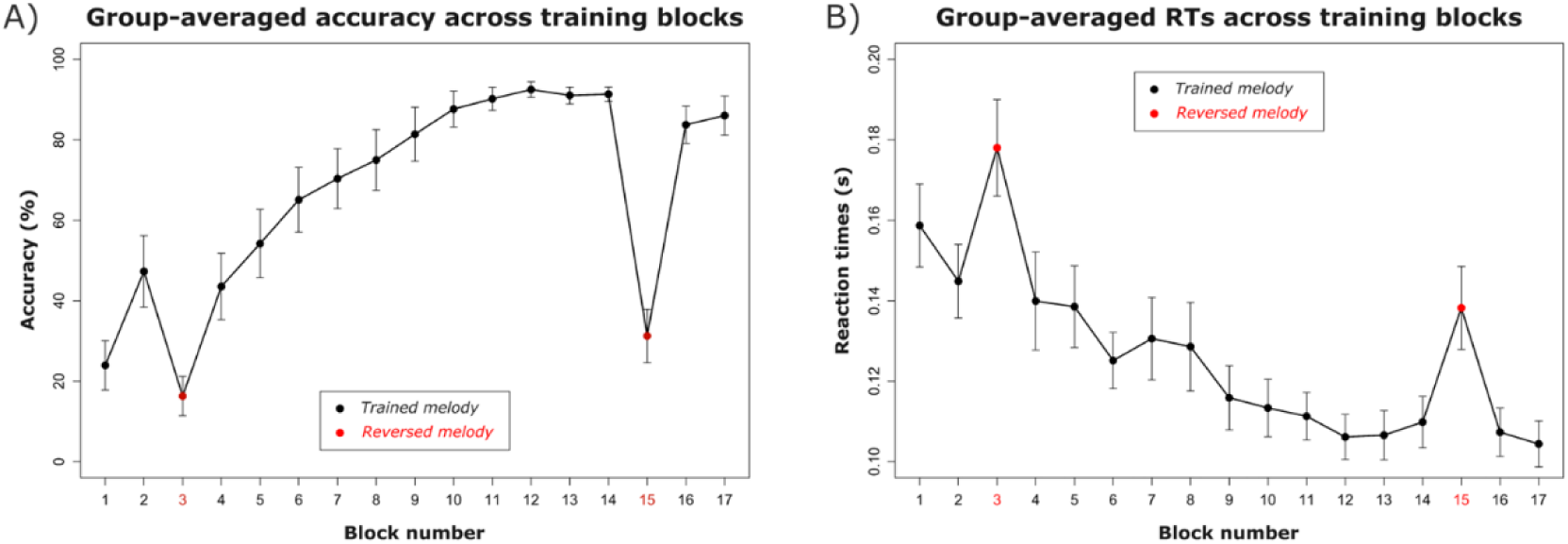
Behavioural performance scores of the TRAIN condition. **A)** Group-averaged accuracy across training blocks. Overall accuracy scores improved throughout the TRAIN condition. Reversed melody blocks 3 and 15 exhibit a significant drop in performance compared to Trained melody blocks 2 and 14, indicating sequence-specific learning of the Trained melody. **B)** Group-averaged RTs across training blocks. Overall reaction times diminished throughout the TRAIN condition. Reversed melody blocks 3 and 15 exhibit a significant increase in latency compared to Trained melody blocks 2 and 14, indicating sequence-specific learning of the Trained melody.

**Figure 3.**
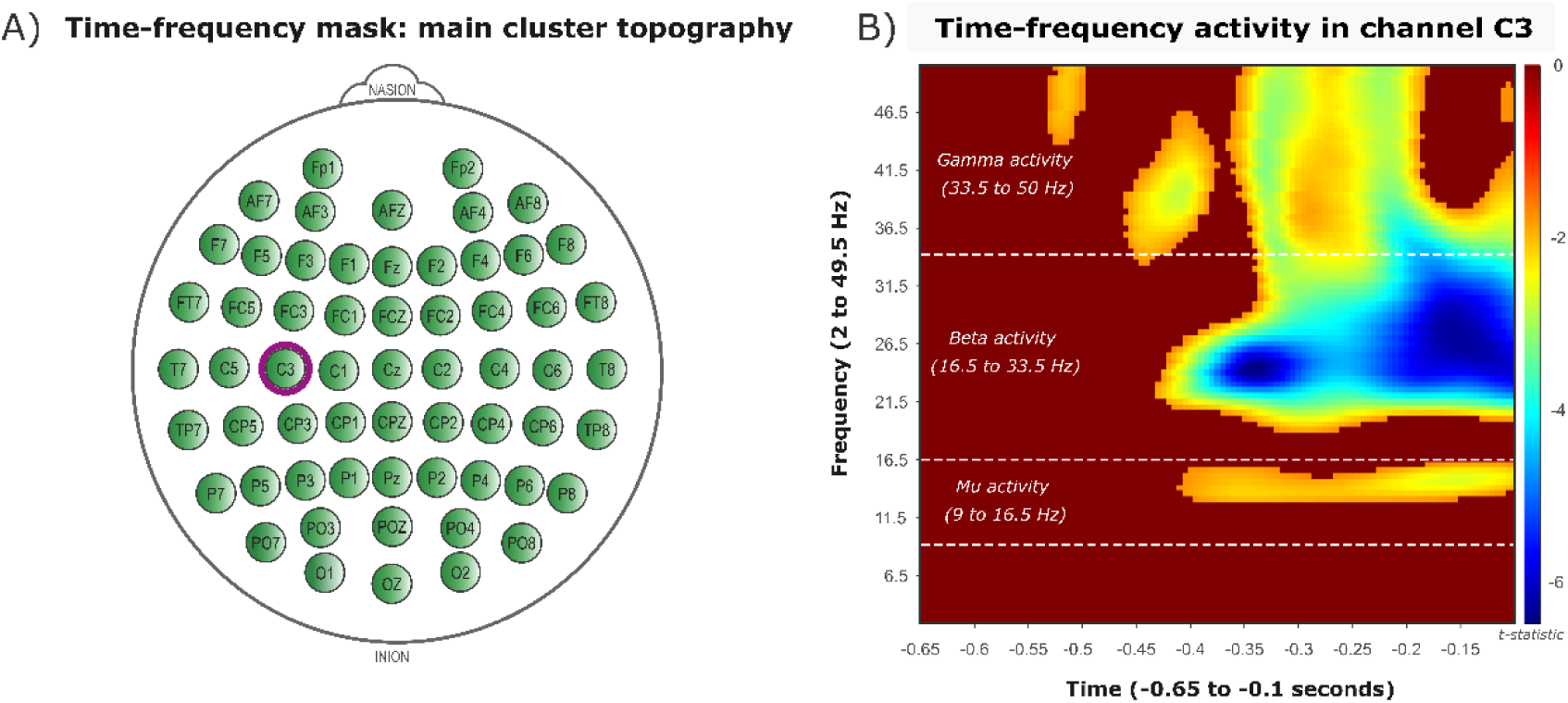
Results of the time-frequency mask derived from the TRAIN condition. **A)** Topography of the time-frequency mask. **Green** indicates channels that survived the cluster-permutation test (n = 60). Channel C3 is highlighted in purple. **B)** Spectrotemporal distribution of the time-frequency mask in channel C3. Coloured pixels are those that survived standard thresholding (*p* <.05). The colour coding indexes the t-statistic at each pixel. Surviving pixels were masked equally onto the passive data in binary fashion, regardless of *t*-statistic value. The cluster activity was further divided into **mu** (9-16.5 Hz), **beta** (16.5-33.5 Hz), and **gamma** (33.5-49.5 Hz) frequency bands. The spectral and temporal distribution of the surviving pixels in these frequency bands varied slightly across channels.

After defining the mu-beta-gamma partitions, we used the thresholded coordinates from the active data to mask the passive data. For each participant and passive listening block, all nonzero baseline-corrected power values (representing the difference between signal power and the time-frequency baseline at each pixel) contained in the mask boundaries were averaged within each frequency band separately, over all channels that displayed activity in that band (33 for the mu band and 60 for the beta and gamma bands) and across the entire time window of interest (−650 to −100 milliseconds relative to note onset). This procedure yielded one scalar value per participant, per frequency band, and per passive listening block to be used in subsequent statistical analyses.

### Statistical analyses

#### Behavioural performance (TRAIN)

A Generalized Linear Mixed Model (GLMM) approach was used for all statistical analyses involving behavioural measures because of the non-normal distribution of the data and this method’s capacity to account for random effects associated with individual differences (Stroup 2012). All GLMM analyses were conducted under the `glmmTMB` package (Brooks et al. 2017) in RStudio version 2022.02.0 (Posit Software 2022).

Two sets of statistical analyses were performed to verify that participants had successfully learned the Trained melody sequence. The first analysis tested for learning across the 15 Trained melody blocks using a one-factor, within-subjects design in two separate GLMMs: one with accuracy and the other with reaction time as the dependent variable. In both, block was included as a fixed within-subjects factor. The same block factor was incorporated into the random effects structure—allowing for random slopes and intercepts—with subject ID set as the grouping variable (1 + block || subject).

The second analysis tested for sequence-specific learning using a two-factor, within-subjects design across two additional GLMMs: one for accuracy and one for reaction time. These analyses compared average performance for Trained melody blocks 2 and 14 to the two Reversed melody blocks (3 and 15). Both GLMMs included condition (Trained vs. Reversed) and block (start vs. end of training) as fixed within-subjects factors, along with their interaction term (condition × block). These two factors were also included in the random effects structure to allow for random slopes and intercepts, and subject ID was set as the grouping variable (1 + condition + block || subject).

Accuracy data exhibited a bimodal, non-normal distribution. As such, percentage scores were converted to proportions and modelled using a beta distribution with a logit link function. Reaction time data closely followed a gamma distribution, as expected for response latency measures (McGill and Gibbon 1965), and were therefore modelled using a gamma distribution with an inverse link function. Across GLMMs, omnibus tests were evaluated using *F* statistics, and FDR-corrected *t*-statistics were computed from two-tailed, paired-sample *t*-tests on the prediction scale. Resulting metrics were back-transformed to the response scale. Degrees of freedom were specified as the residual degrees of freedom at the observation level.

#### EEG activity during passive listening (PRE and POST)

To test whether EEG activity would exhibit the predicted mu suppression exclusively during post-training passive listening to the Trained melody, we performed left-tailed, one-sample, *t*-tests against zero on the 25 channel-averaged time-frequency plots, separately for each of the four passive listening blocks, within each of the three frequency bands. Because a total of 12 tests were conducted (4 passive listening blocks × 3 frequency bands), significance was corrected for multiple comparisons with non-parametric permutation testing (10,000 iterations).

We further tested for factorial interactions by implementing three GLMMs, one for each frequency band. Specifically, we used a 2×2 within-subjects design with condition (Trained vs. Untrained) and time (Pre-vs. Post-training) as fixed effect factors, along with their interaction term (condition × time). These two factors were also included in the random effects structure, which allowed for random slopes and intercepts, with subject used as the grouping variable (1 + condition + time || subject). The dependent variable in each model was the 25 channel-averaged, baseline-corrected signal power estimate values in each of the four passive listening blocks. The dependent variable was modelled using a Gaussian distribution, as differential values resulting from baseline correction operations are expected to approximate normality (Field 2013).

Omnibus tests were computed from the model using *F* statistics, and uncorrected *t*-statistics were derived from two-tailed, paired-sample *t*-tests conducted directly on the response scale. Degrees of freedom at the observation level were defined by the model’s residual degrees of freedom. Because this GLMM structure was applied across three frequency bands, significant *t*-statistics were corrected for multiple comparisons using non-parametric permutation testing (10,000 iterations).

Finally, we assessed relative pre-post changes between Trained and Untrained melodies. This approach explicitly accounted for differences in post-training values relative to baseline pre-training values, which may have been obscured when examining absolute changes under a factorial design. For each frequency band, relative change scores were calculated by subtracting pre-training baseline-corrected power estimate values from post-training values and scaling the result by the absolute value of the former. Given the expected presence of outliers resulting from this type of ratio-based calculation, statistical comparisons were conducted using a non-parametric Wilcoxon signed-rank exact test. As these tests were conducted across three frequency bands, significance levels were corrected for multiple comparisons with non-parametric permutation testing (10,000 iterations).

#### Brain-behaviour correlations

To assess whether individual differences in motor learning in the TRAIN condition were associated with changes in brain activity in the PRE and POST passive listening conditions, we conducted four Spearman rank-order correlation tests. The degree of motor learning was quantified by computing the slope of change in average accuracy and RTs across Trained melody blocks 4-14 of the motor training task. Individual accuracy and RT slopes were then correlated with individual scores in two EEG-derived measures within the mu frequency band: (1) baseline-corrected power estimate values during passive listening to the Trained melody in the post-training block, representing *absolute* mu suppression activity; and (2) the normalized pre-post difference between Trained and Untrained melodies, representing *relative* mu suppression activity after training with respect to before training. Spearman’s rank-order method was used due to the presence of non-normality and potential outliers in the data.

## Results

### Attentional control task

As an attentional control, seven mistuned notes were presented during each of the PRE and POST passive listening conditions. Across both conditions, participants reported an average 10.44 and 7.52 mistuned notes, indicating that they were actively engaged in listening during the passive task.

### Behavioural performance

We tested for a main effect of motor learning across blocks of training. For accuracy, we observed a significant effect of block (*F*(14, 344) = 47.53, *p* <.001, *η*² = 0.66), indicating a global increase in accuracy as training progressed (Figure 2A). For RT, we similarly observed a significant main effect of block (*F*(14, 344) = 19.96, *p* <.001, *η*² = 0.45), indicating a concomitant global decrease in reaction times over the course of training.

Next, we tested for sequence-specific learning by comparing Trained melody blocks 2 and 14 to Reversed melody blocks 3 and 15. For accuracy, we observed a significant main effect of both condition (*F*(1, 92) = 48.4, *p* <.001, *η*² = 0.34)—such that accuracy was greater for Trained than Reversed melodies throughout training—and block (*F*(1, 92) = 28.05, *p* <.001, *η*² = 0.23), such that accuracy increased from Start to End of training across both melody types. We also obtained a significant interaction term (*F*(1, 92) = 12.8, *p* <.001, *η*² = 0.12), such that the start-to-end increases in performance were comparably greater for Trained melodies than Reversed melodies, as revealed by a second-order interaction test (*t*(92) = 4.72, *p* <.001, Cohen’s *d* = 0.49).

For RT, we similarly observed significant main effects for both block (*F*(1, 92) = 22.77, *p* <.001, *η*² = 0.2)—indicating shorter RTs throughout training for the Trained melody compared to the Reversed melody (*t*(92) = −4.66, FDR-corrected *p* <.001, Cohen’s *d* =-0.48)—and time (*F*(1, 92) = 39.1, *p* <.001, *η*² = 0.3), indicating shorter RTs at the End compared to the Start of training for both melody types (*t*(92) = −5.79, FDR-corrected *p* <.001, Cohen’s *d* =-0.6). The interaction term was not statistically significant (*F*(1, 92) = 0.9, *p* =.34, *η*² = 0.01). Together, these results indicate that participants learned the specific Trained melody sequence well.

### Brain activity

#### Time-frequency mask

The cluster permutation test yielded a time-frequency *t*-map (Figure 3A) comprising 60 channels, 97 frequency bins, and 110 timepoints (sampled at 200 Hz). Channel FPz had been used as ground, and channels FT9, FT10, TP9, and TP10 had been used as external electrodes, so all five were excluded from the test.

The time-frequency *t*-map or mask comprised one significant *channel × frequency × time* cluster (1,000 permutations, *p* =.001). The cluster (cluster-level statistic [maxsum] = −725,558; size = 238,299 pixels) encompassed all 60 channels, spanned frequencies specifically from 9 to 49.5 Hz, and featured activity across the entire time window ranging from −650 to −100 milliseconds (see Supplementary Fig. 2). Within this cluster, we examined three frequency bands: mu (9-16.5 Hz; **33** channels), beta (16.5-33.5 Hz; **60** channels), and gamma (33.5-49.5 Hz; **60** channels; Figure 3B).

The activity in the mu frequency band resulting from the cluster permutation test was interpreted as reflecting true, active mu suppression prior to note onset, based on its distinct, spectrally and spatially confined location within the cluster (Figure 4A). Namely, this activity was restricted to a reduced subset of 33 channels—29 of which were spectrally isolated from the beta band (see Figure 3B for an example in C3)—and was left-lateralized, as expected and required for a right-hand response. In other words, while the beta and gamma frequency bands recruited all available channels, the mu frequency band displayed a left-lateralized distribution with no right-central electrode recruitment, consistent with previous work (Pineda 2013).

**Figure 4.**
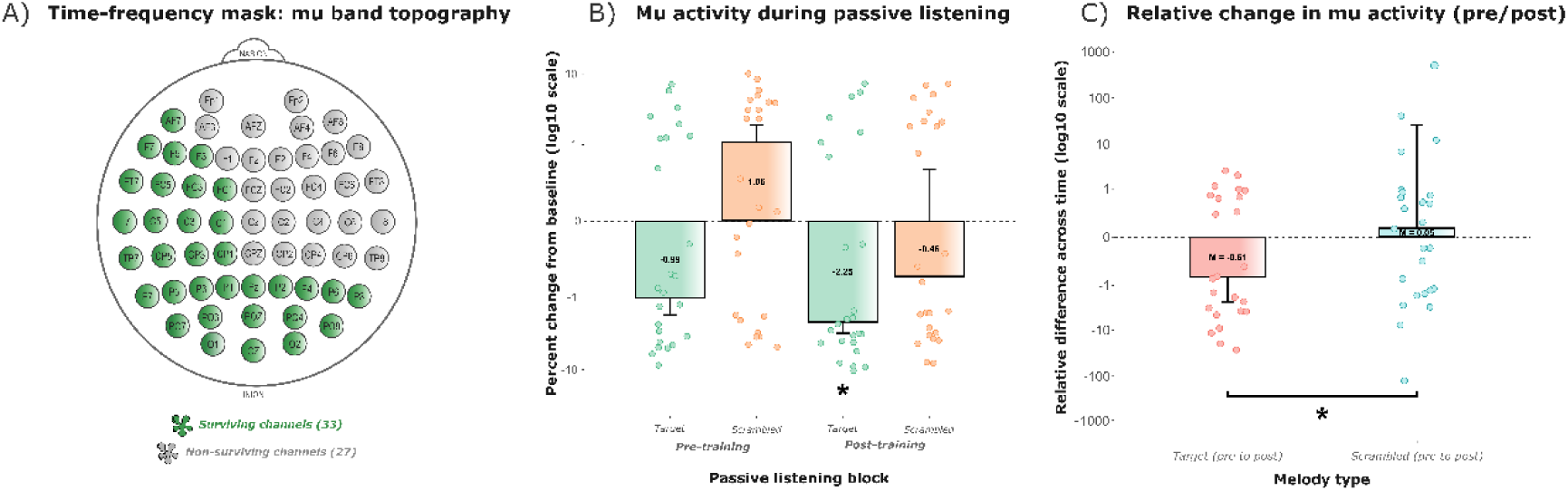
Mu band activity across PRE and POST conditions. **A)** Topography of the mu frequency band. **Green** indicates channels in the cluster that display mu activity (n = 33). **Grey** indicates the channels in the cluster that do not display any mu activity (n = 27). **B)** Subject-averaged mu activity within each passive listening block. Baseline-corrected power estimate values representing the difference between signal power and baseline in each pixel of the mu frequency band were averaged across channels within each passive listening condition, for each subject. The post-training Trained melody block was the only block that displayed significant group-level suppression with respect to zero in the mu frequency band. Data are presented in a logarithmic scale for display purposes. **C)** Relative pre-post change in mu activity for Trained vs Untrained melodies. Relative change scores were computed by subtracting pre-training baseline-corrected power estimate values from post-training values and dividing by the absolute value of the former. The Trained melody exhibited significantly greater median suppression across time compared to the Untrained melodies. Data are presented in a logarithmic scale for display purposes.

#### EEG activity during passive listening blocks

To test the presence of suppression across the four passive listening blocks, we conducted a series of 12 left-tailed, one-sample *t*-tests against zero (Fig. 4B; Table 1; Supplementary Fig. 2) across the mu, beta and gamma frequency bands. Results revealed significant suppression relative to baseline only within the pixels representing activity in the mu frequency band, and only during post-training passive listening to the Trained melody (*t*(24) = −2.4, permutation-corrected *p* =.01, Cohen’s *d* =-0.48).

**Table 1.** Left-tailed, one-sample *t*-tests against zero.

| Frequency band | Condition | *df* | *t*-value | *p*-value | Effect |
| --- | --- | --- | --- | --- | --- |
| **Mu** | Target melody PRE training | 24 | -1.3 | .10 | -0.26 |
|  | Scrambled melodies PRE training | 24 | 1.32 | .90 | 0.26 |
|  | **Target melody POST training** | **24** | **-2.4** | **.01\*** | **-0.48** |
|  | Scrambled melodies POST training | 24 | -0.57 | .29 | -0.11 |
| **Beta** | Target melody PRE training | 24 | -0.46 | .32 | -0.09 |
|  | Scrambled melodies PRE training | 24 | 1.07 | .85 | 0.21 |
|  | Target melody POST training | 24 | -0.5 | .30 | -0.1 |
|  | Scrambled melodies POST training | 24 | 0.45 | .67 | 0.09 |
| **Gamma** | Target melody PRE training | 24 | -1.34 | .10 | -0.27 |
|  | Scrambled melodies PRE training | 24 | 1.41 | .91 | 0.28 |
|  | Target melody POST training | 24 | -1.24 | .11 | -0.25 |
|  | Scrambled melodies POST training | 24 | -0.19 | .42 | -0.04 |

**Note.** Asterisk (\*) indicates significance at $p < .05$ (uncorrected).

Within the mu frequency band, GLMM results additionally revealed a significant main effect of condition (*F*(1, 92) = 5.2, *p* =.02, *η*² =.05). A subsequent post hoc test indicated that the change in power with respect to time-frequency baseline (baseline-corrected power estimate values) for the Trained melody was negative and greater than for the Untrained melodies (*t*(92) = −1.92, permutation-corrected *p* =.047, Cohen’s *d* =-0.2). Neither the main effect of time (*F*(1, 92) = 3.18, *p* =.08, *η*² =.03) nor the interaction term (*F*(1, 92) = 0.03, *p* =.87, *η*² <.001) reached significance. By contrast, the same GLMM structure revealed no significant main effects or interactions within the beta frequency band (condition: *F*(1, 92) = 1, *p* =.32, *η*² =.01; time: *F*(1, 92) = 0.03, *p* =.87 *η*² <.001; condition × time: *F*(1, 92) = 0, *p* =.95; *η*² <.001), nor within the gamma frequency band (condition: *F*(1, 92) = 2.38, *p* =.13, *η*² =.03; time: *F*(1, 92) = 0.67, *p* =.41, *η*² <.01; condition × time: *F*(1, 92) = 0.6, *p* =.44, *η*² <.01).

To further test whether passive mu suppression was greater in response to the Trained melody following motor learning, we assessed the relative pre-post change in activity.

The Wilcoxon signed-rank exact test revealed significantly greater normalized pre-post mu suppression for the Trained melody compared to the Untrained melodies (Figure 4C; *V* = 97, permutation-corrected *p* =.04, *r* =.41). Conversely, no significant differences were observed in the beta frequency band (*V* = 140, *p* =.28, *r =*.22) or in the gamma frequency band (*V* = 217, *p* =.93, *r* =.02). These findings confirm that, when explicitly accounting for pre-post differences, a significant relative suppression emerges for the Trained melody only within the mu frequency band, thereby complementing and extending the results of the one-sample *t*-tests against zero and the factorial design.

#### Brain-behaviour correlations

To test for a relationship between individual differences in learning and mu suppression, we correlated one absolute measure and one relative measure of mu suppression with learning slopes representing change in accuracy and RT during motor training. Neither correlation between the accuracy slope and the two measures of brain activity were statistically significant (post-training Trained melody values: *ρ* =.25, *p* =.23; relative pre-post difference between Trained and Untrained melody types: *ρ* = −.04, *p* =.85). Equally, neither correlation between the RT slope and the two measures of brain activity were statistically significant (post-training Trained melody values: *ρ* =.03, *p* =.89; relative pre-post difference between Trained and Untrained melody types: *ρ* = −.27, *p* =.19).

## Discussion

The goal of this study was to assess the learning of new auditory-motor representations by quantifying the degree of anticipatory mu suppression occurring at the single-note level during passive listening to trained versus untrained melodies. To test these ideas, we generated a time-frequency mask to objectively identify the pattern of predicted oscillatory responses based on the active training data. This mask was used to parse the EEG data obtained during pre and post passive listening conditions. Results indicate that mu suppression was significantly pronounced only in response to the trained melody sequence after motor learning, and that no significant suppression effects were present in the other frequency bands (beta and gamma). Thus, our findings support the idea that mu suppression can be observed and measured at the single-note level in non-musicians after training, and that this effect is frequency-band specific. Overall, these findings align well with the current literature on the proactive role of covert, predictive motor processing of time-sensitive regularities in support of auditory perception.

Our behavioural results demonstrate that subjects successfully learned the target melody and that this learning was sequence-specific (Figure 2). Supporting our principal hypothesis, we obtained significant mu suppression only during passive listening to the target melody post-compared to pre-training (Figure 4B). Moreover, this significant suppression was not found in the other two frequency bands, beta and gamma (Table 1; Supplementary Fig. 1).

The greater post-training mu suppression we obtained while listening to the trained melody aligns with neuroimaging studies showing that passive listening to learned melodies elicits activity in motor areas (Baumann et al. 2007; Lahav et al. 2007; Lappe et al. 2008; Herholz et al. 2016). For example, Haueisen and Knösche (2001), demonstrated that passive listening to well-trained melodies increased activity over the contralateral M1 hand area, the putative generator of mu suppression (Pineda 2013).

Our evidence for mu suppression in non-musicians after a single session of training additionally suggests that similar predictive models are developed over both short and long-term practice, and that these mechanisms are common to both musicians and non-musicians. This conclusion is supported by early spectral analyses showing that short-lasting phasic mu suppression during voluntary movement is observable in essentially all investigated healthy adult participants (Pfurtscheller and Aranibar 1979), as well as functional connectivity studies indicating that the auditory and motor systems exhibit a strong resting functional connection in the brain that is independent of musicianship (Bedford et al. 2025).

Importantly, for a sequence of motor predictions to be paired to a sequence of auditory predictions, specific auditory-motor associations must necessarily have been formed. Note that this is not a prerequisite condition during beat production and perception tasks (Iversen and Patel 2008), where beta-mediated motor predictions arise during natural production and listening (Fujioka et al. 2015), as well as when meter representations are manipulated internally even in the absence of movement (Morillon and Baillet 2017). For this reason, we argue that the mu suppression observed in our study specifically reflects the prior establishment of auditory-motor associations, and that this is why it is seldom observed alongside beta activity in beat-perception tasks. By the same token, our findings support the notion that predicting *which* event will occur next within a sequence places different demands on cortical motor areas than simply predicting *when* a given event will occur based solely on periodicity (Novembre and Keller 2014; Stupacher et al. 2013).

Convergence for this idea can be found in Ross et al. (2022b), where musicians and non-musicians were exposed to untrained musical fragments and displayed mu enhancement—rather than suppression—in response. The authors interpreted this enhancement as suggestive of motor inhibition, noting that if mu suppression had been observed instead, it would likely have reflected motor imagery. We agree with this interpretation and propose that mu suppression would have emerged if participants had been trained on the melodies beforehand. In other words, because no auditory–motor associations had been established, there was no basis for eliciting the anticipatory, effector-specific motor imagery that we argue mu suppression indexes.

Taken together, our interpretation of mu suppression as reflecting anticipatory, effector-specific motor imagery aligns with prior findings in musicians (Wu et al. 2016). However, our results stand in contrast to the null effect reported in non-musicians by the same authors (Wu et al. 2017), a discrepancy we attribute to key methodological differences. First, Wu et al. (2017) did not use active mu suppression data to localize suppression during passive listening. Our results suggest that leveraging the active stronger mu suppression may be necessary for isolating the potentially fainter passive mu suppression in non-musicians. Second, given a sample size of only 13 subjects, the authors may have lacked statistical power to detect the comparably subtler passive mu suppression. Finally, training and passive exposure occurred in separate sessions in Wu et al. (2017), suggesting that suppression in non-musicians may be short-lived or more easily detected immediately after training.

In the current study we did not observe correlations between mu suppression and changes in accuracy or RT. If mu suppression is an index of motor prediction or preparation, we would expect greater gains in performance to be related to greater post-training suppression. Thus, it is possible that performance might be more closely linked to individual differences in peaks or latencies in the mu band that were not captured by our group-level time-frequency mask.

One key strength of our study is that we created a time-frequency mask to identify active mu suppression in the motor training task, allowing us to constrain the time-frequency-channel space to reveal mu suppression during passive listening. The pattern of time-frequency modulations observed in the active condition is consistent with the frequency range typically ascribed to the mu rhythm (9-16.5 Hz; Pfurtscheller and Lopes da Silva 1999). Although the mu rhythm is often defined within the 9-13 Hz range, several studies extend this range up to ∼16 Hz (Wilson et al. 2010; Woodruff et al. 2011), and reviews likewise report broader definitions such as 10-14 Hz (Hobson and Bishop 2016). Furthermore, the mu activity observed in our data is clearly distinct from lower beta activity, as the two bands are separated by several frequency bins across most channels (Fig. 3B; Supplementary Fig. 2). Moreover, while mu rhythms sometimes include a second component or harmonic in the beta range (Hari and Salmelin 1997), our results do not support the idea that this second mu-beta component was engaged.

The spatial extent of our mu partition is also consistent with the theoretical topography of mu suppression, given that we obtained a clustering of left-central channels around M1—the putative generator of mu rhythms (Pfurtscheller and Neuper 1997), as supported by evidence from intracranial EEG recordings (Jasper and Penfield 1949), MEG source-space analysis (Salmelin and Hari 1994), and cortex-muscle coherence (Salenius et al. 1997). This being said, it is important to note that sensor-level EEG has a limited capacity to pinpoint exact neural sources and generators, especially when compared to source-resolved MEG (Baillet 2017).

One key consideration in measuring mu suppression is the potential confound of occipital alpha, which can be caused by either visual or attentional processes (Hobson & Bishop 2016). In the current study, the inclusion of posterior channels in the mu partition of the mask is likely caused by volume conduction (Cohen 2014) and not a sign of occipital alpha. On the one hand, visual alpha is unlikely to be present, as the inclusion of a fixation cross within motor trials and passive data greatly diminishes the possibility of meaningful within-trial visual processing differences between the time-frequency baseline and the subsequent analysis window (-650 to-100 ms). On the other hand, while consistent within-trial attentional changes could have theoretically affected the training data, these would have had to be consistent across participants to feasibly distort the group localizer. Moreover, this contamination is unlikely to have crossed over to the subject-level passive listening data after masking, as within-trial attention was tightly controlled within and across passive listening trials via the distractor task of counting mistuned notes. Thus, the risk of occipital alpha contamination due to consistent attentional differences between the time-frequency baseline and the final analysis window is also unlikely and, at any rate, would not explain the differences in mu power we observed between passive listening blocks.

At first glance, our chosen timeframe of −650 to −100 milliseconds with respect to tone onsets may seem misaligned with the longer onsets of −2000 milliseconds typically reported in classical mu suppression studies (Pineda 2005), where this component is typically derived at the whole-trial level. However, given the short inter-onset interval of 1500 milliseconds in our task, a longer baseline would result in overlap with the preceding note, making it impractical. More importantly, to our knowledge no established benchmark currently exists for capturing mu suppression at the single-event level (Fox et al. 2016), which is the timescale used in the current study. Nevertheless, our time window appears appropriate as it is highly consistent with other measures of motor preparation. For instance, the onset of mu activity in our motor task (Figure 3B) closely overlaps with the onset of around −400 milliseconds of the late component of the lateralized readiness potential, an ERP marker of motor preparation that also originates in M1 and lateralizes contralaterally to the effector (Shibasaki and Hallett 2006; Trevena and Miller 2002).

The temporal mid-point of the observed mu-band suppression also coincides with the timing of the motor-evoked potentials (MEPs) elicited during post-training passive listening in our previous study using the same paradigm (Stephan et al. 2018). MEPs index motor excitability in M1 and have been used to measure the degree of motor preparation in the system in similar musical contexts (D’Ausilio et al. 2006). Moreover, the MEPs elicited in a previous study from our group (Stephan et al. 2018) were effector-specific, just as the mu suppression observed in our current study is linked to the anticipation of specific notes, and therefore to specific effectors. Moreover, our findings align with prior MEG findings in passive listening to trained melodies, where distinct spatial responses were found for notes associated with the thumb versus pinkie fingers that matched the homuncular organization in M1 (Haueisen and Knösche 2001). Thus, mu suppression may be linked to MEPs, as they both share a common generator in M1 and similar timing (cf. Lepage et al. 2008).

Besides M1, there are reasons to believe that the mu suppression we observed may be contingent on the co-activation of vPMC. On the one hand, because our task required the learning of a fixed, one-to-one sound-movement mapping; the kind of mapping that has been functionally tied to vPMC (Zatorre et al. 2007). Moreover, hand-related representations dominate vPMC’s body map (cf. Graziano et al. 2002) and are thought to support fine motor skills (Rizzolatti et al. 2002), not directly through corticospinal projections but indirectly by modulating excitability in M1 (Davare et al. 2009, Buch et al. 2010). On the other hand, prior fMRI literature has established that vPMC is tied to goal-related action programming (Nieuwenhuys et al. 2008) specifically when instantiated via action sequencing (Rosenbaum 1991), as required in our task. However, BOLD activity in dPMC and nearby motor areas has been concurrently tied to mu suppression (Arnstein et al. 2011). Thus, while vPMC involvement appears most plausible, we cannot rule out a broader dorsal auditory-motor network contribution that may accompany or modulate the effect observed here.

Our finding that active mu suppression preceding movement shares time-frequency coordinates with mu suppression during passive listening to the same auditory stimuli is consistent with prior work demonstrating substantial neural overlap between action and perception (Keller and Koch 2006), particularly within M1 (Fadiga et al. 2005; Arnstein et al. 2011; Pineda et al. 2013). Moreover, this overlap in our results reinforces two important lines of research. First, it strengthens the body of work linking mu suppression to motor imagery and mental simulation, a domain that has been comparatively less explored than mu suppression associated with overt self-movement and action by others (Pineda 2005). Second, our finding lends mechanistic support to the *common coding* theory (Prinz 1997), which proposes that perception and action rely on shared representational substrates. This is noteworthy because, while the notion of a unifying mechanism for perception and action has a rich history (James 1890) and profound implications for human cognition (Hommel et al. 2001), the neural functioning and implementation of such a mechanism are not yet fully understood (Novembre and Keller 2018).

Finally, while regularity-based *auditory* predictions during music listening have been thoroughly investigated (Rohrmeier and Koelsch 2012; Tillmann 2012), the neural basis of *motor* predictions in musical actions has not been explored in depth (Maidhof et al. 2010; Ruiz et al. 2009). Thus, our findings in non-musicians shed light on the neural dynamics of motor predictions during passive listening of learned melodies, indicating they are likely part of innate mechanisms that are globally important for perception (Patel and Iversen 2014), and therefore dissociable from long-term musical training.

## Acknowledgments

We thank Joseph Thibodeau for designing and building the piano-like keyboard used in this study, and for providing invaluable technical support throughout its implementation. We are also grateful to Jennifer Cohen, Ian Marci, Yingrui He, and Sebastian Kolde for their dedicated assistance during piloting and data collection. *Correspondence*: Oscar Bedford, 3801 University Street, Montréal, Québec, H3A 2B4, Canada.

