## Supplementary figures and images for "Mu suppression reveals auditory-motor predictions after short musical training"

### Figure 1

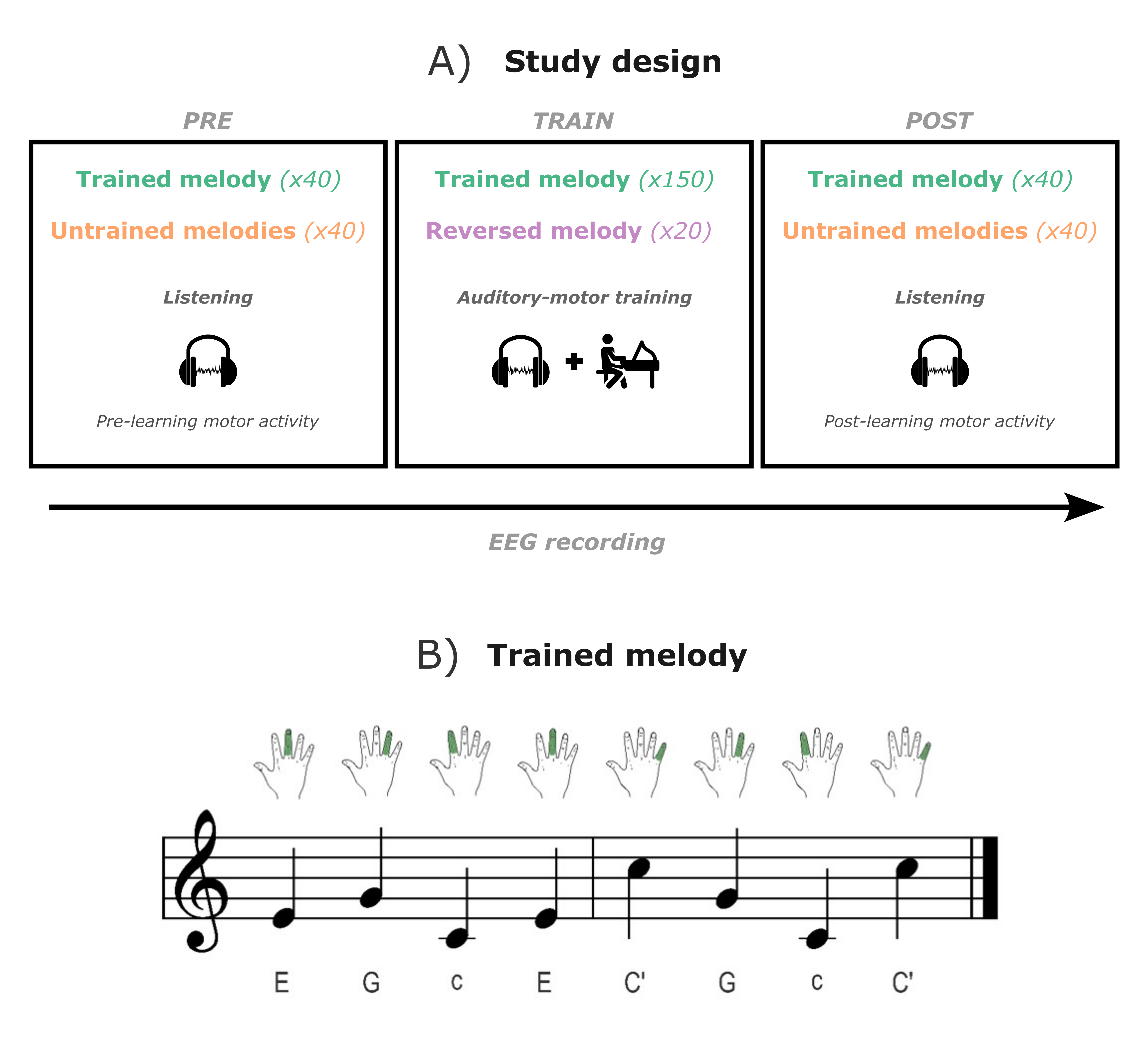

### Figure 2A

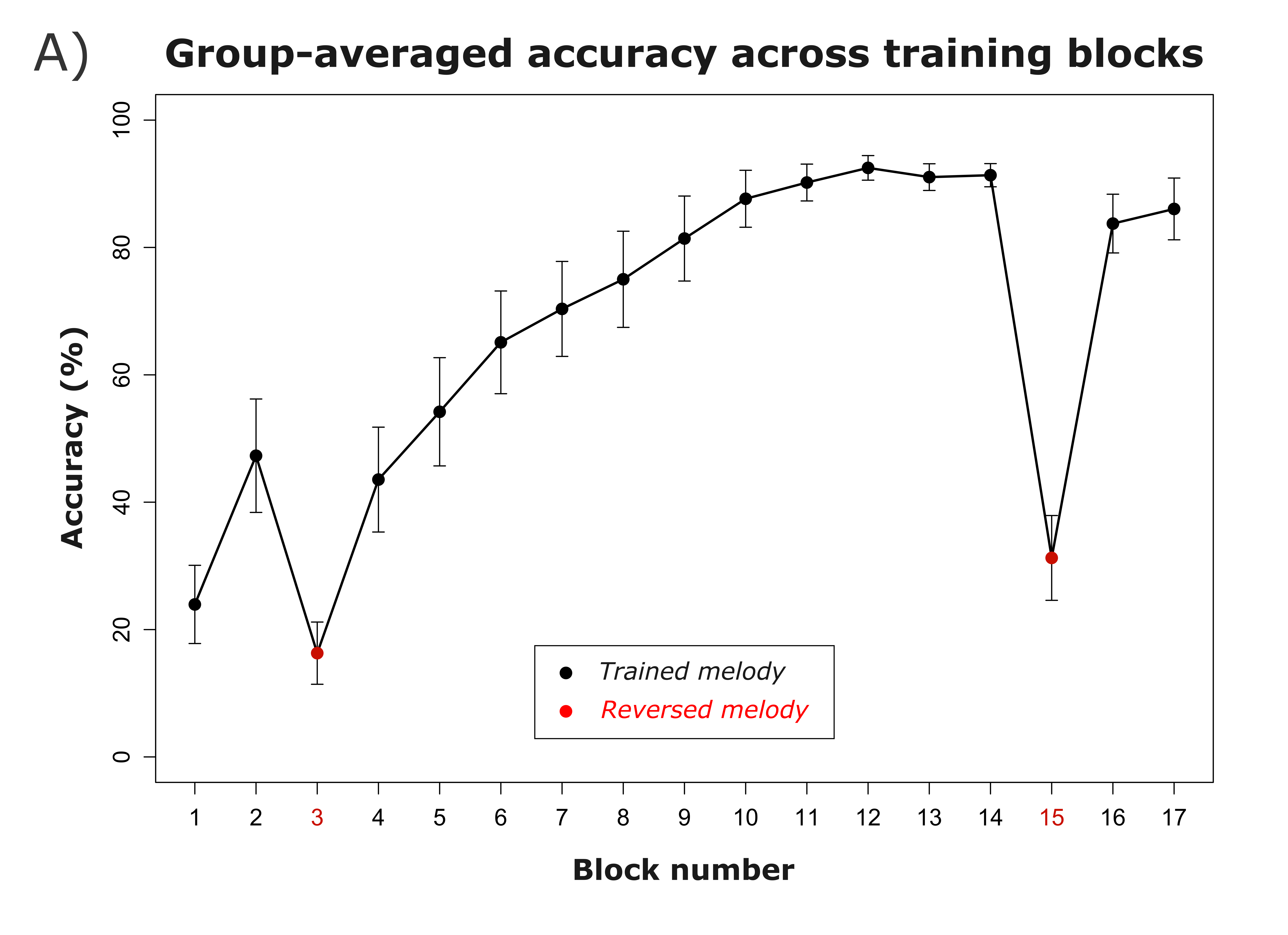

### Figure 2B

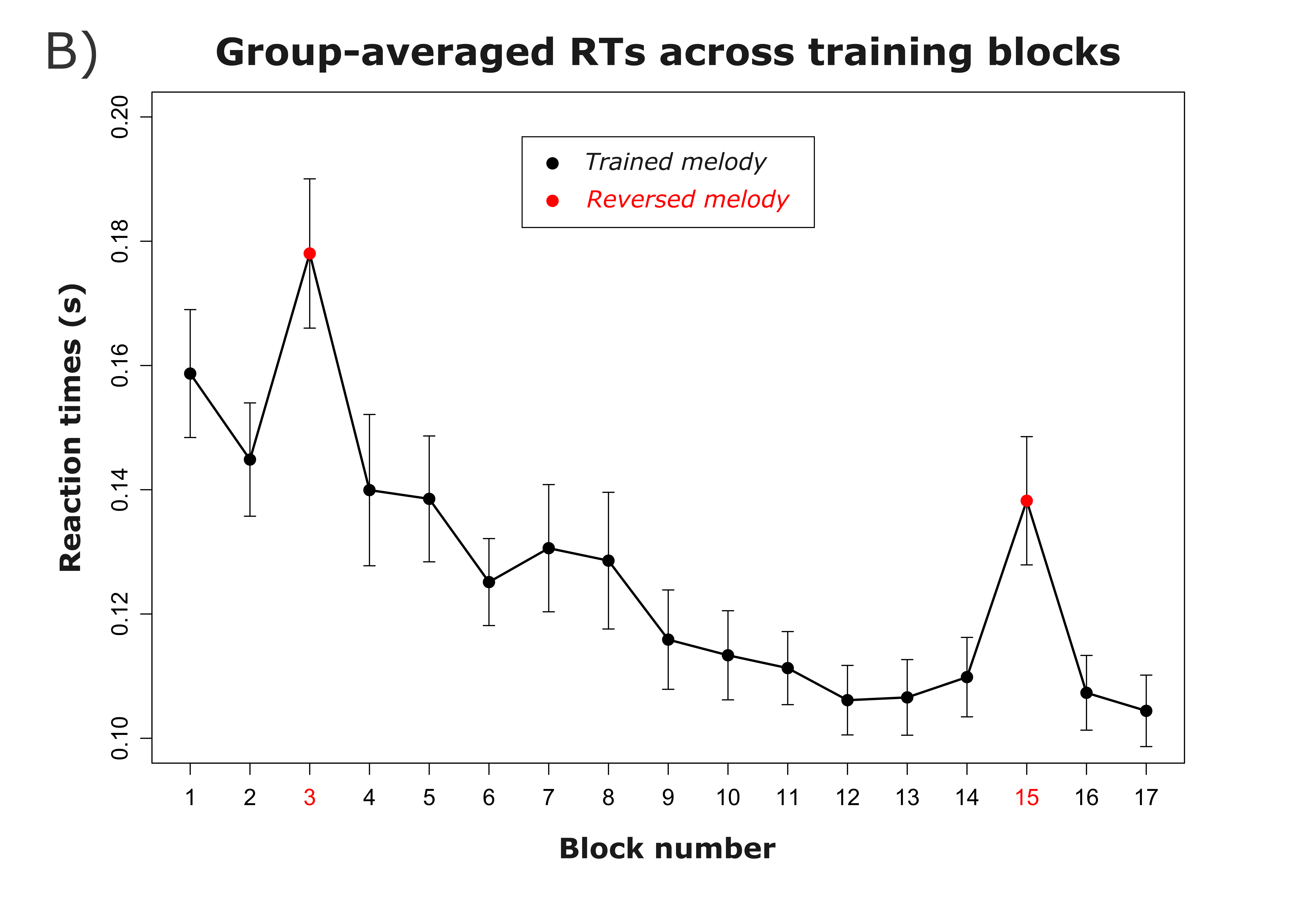

### Figure 3

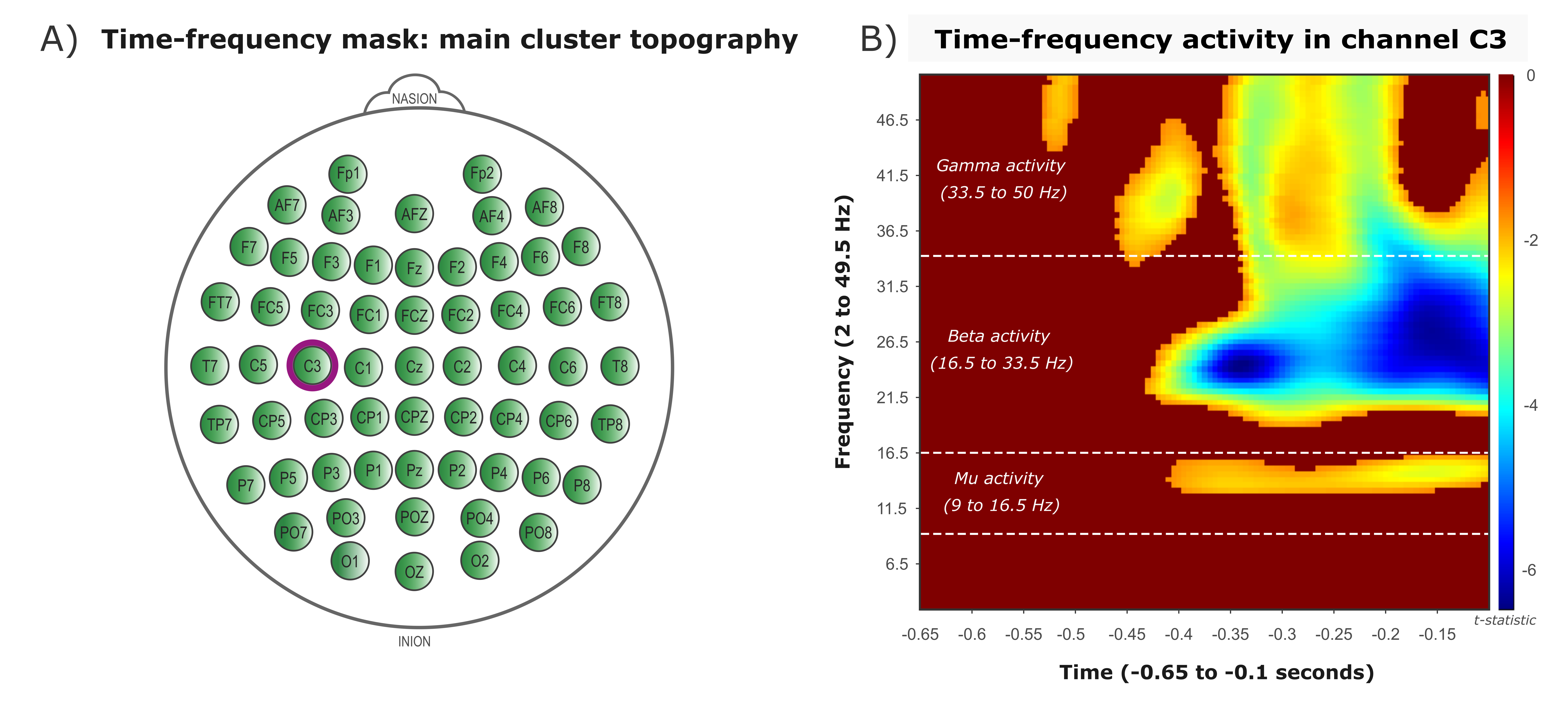

### Figure 4

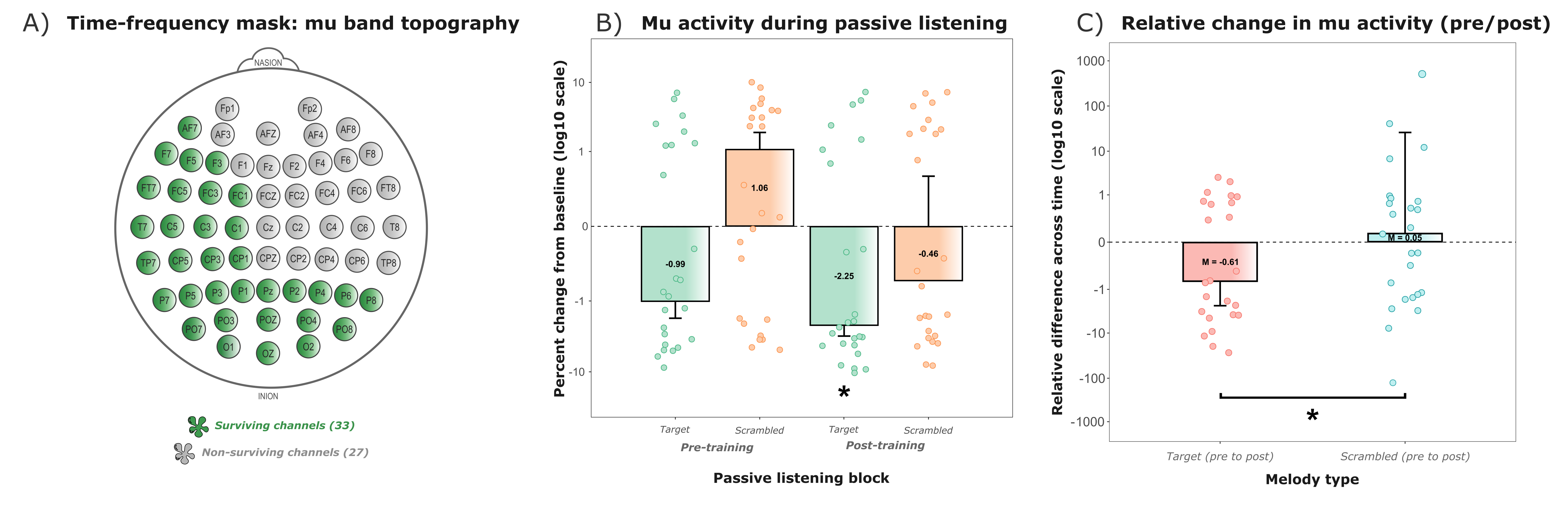

### Supplementary Figure 1

**Supplementary Figure 1.** Activity during passive listening in non-significant bands

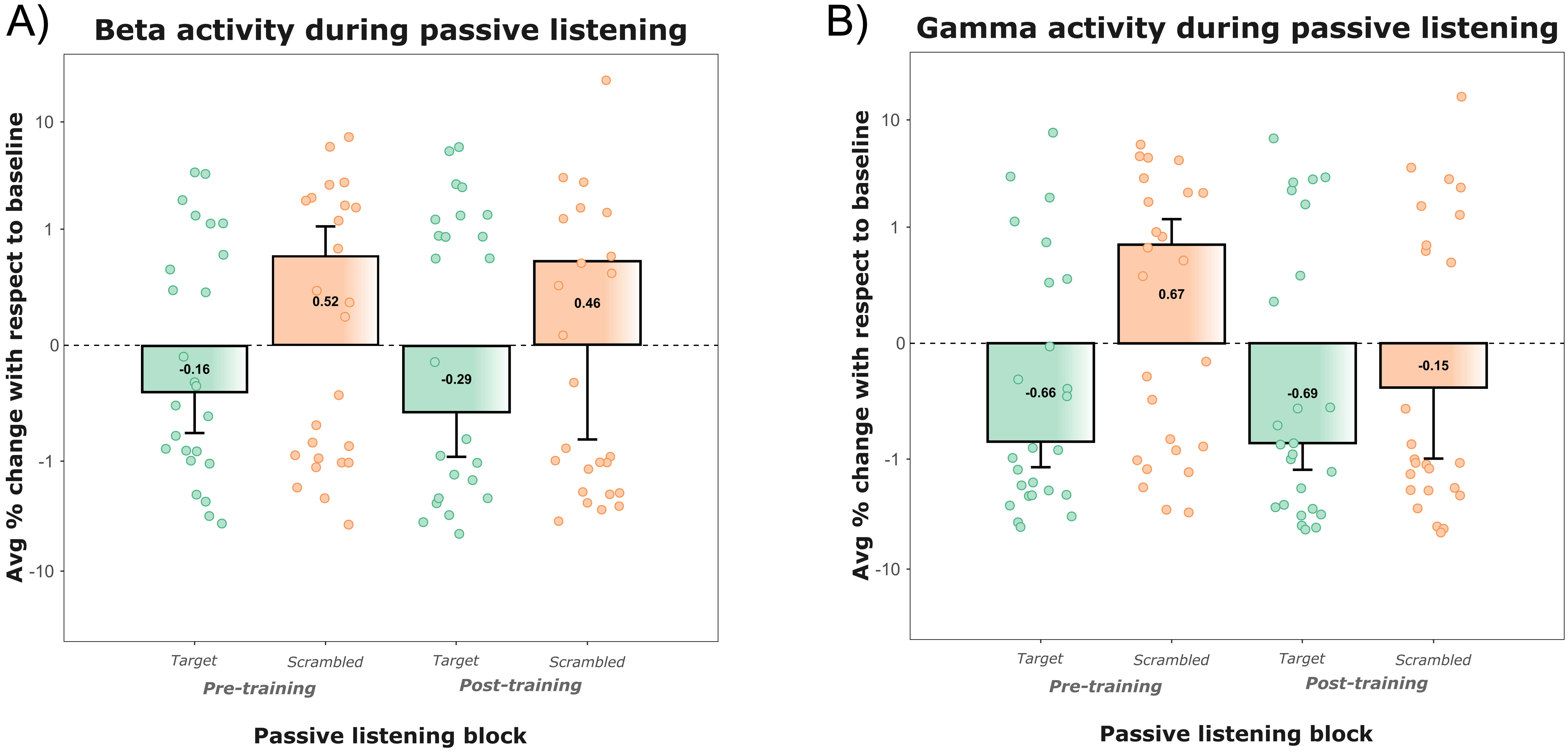
