## Supplementary Figure 2 for "Mu suppression reveals auditory-motor predictions after short musical training"

Time-frequency p-maps of all functional localizer channels (64 channels).

*Legend*.

x-axis: time (s); y-axis: frequency spectrum (Hz);

colorbar (red: $p>.05$; blue: $p = .000999$)

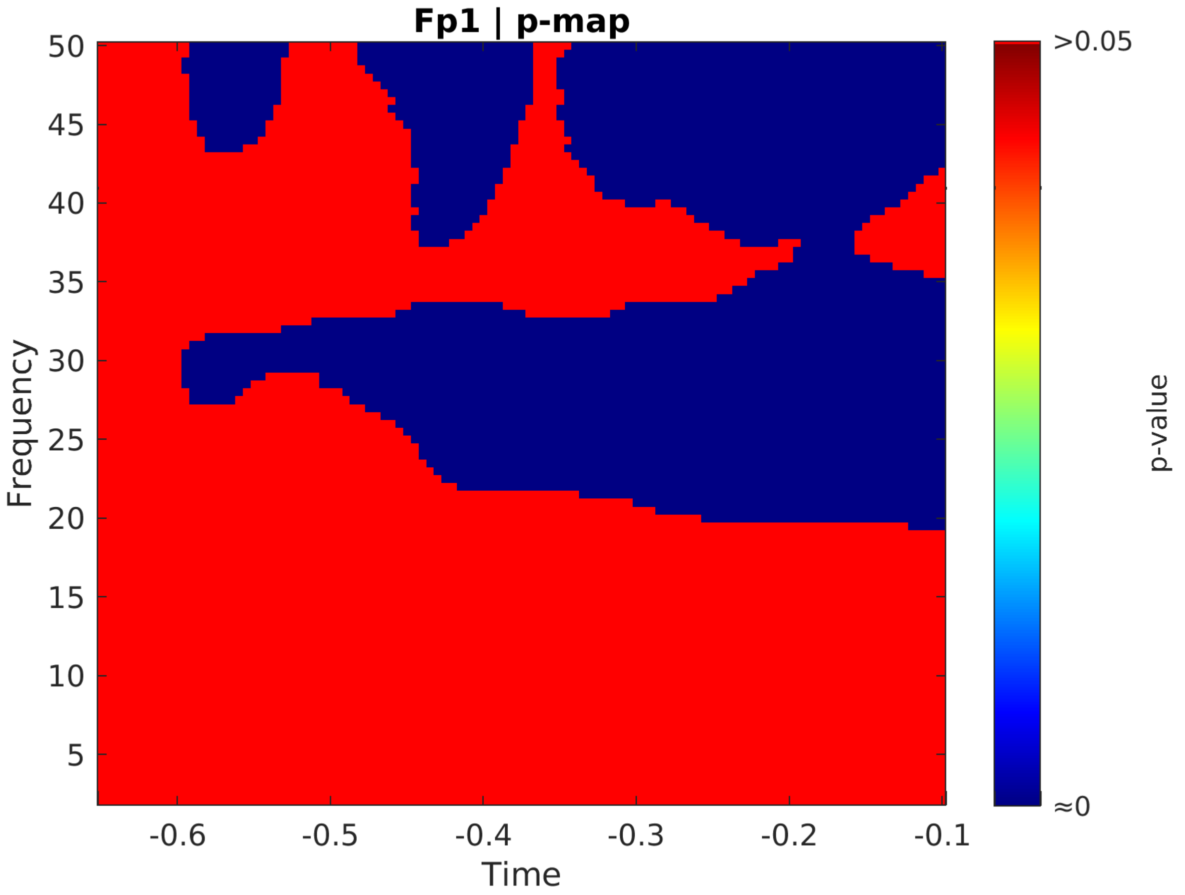

Fp1 | p-map
50
45
40
35
30
25
20
15
10
5
Frequency
-0.6
-0.5
-0.4
-0.3
-0.2
-0.1
Time
>0.05
≈0
p-value

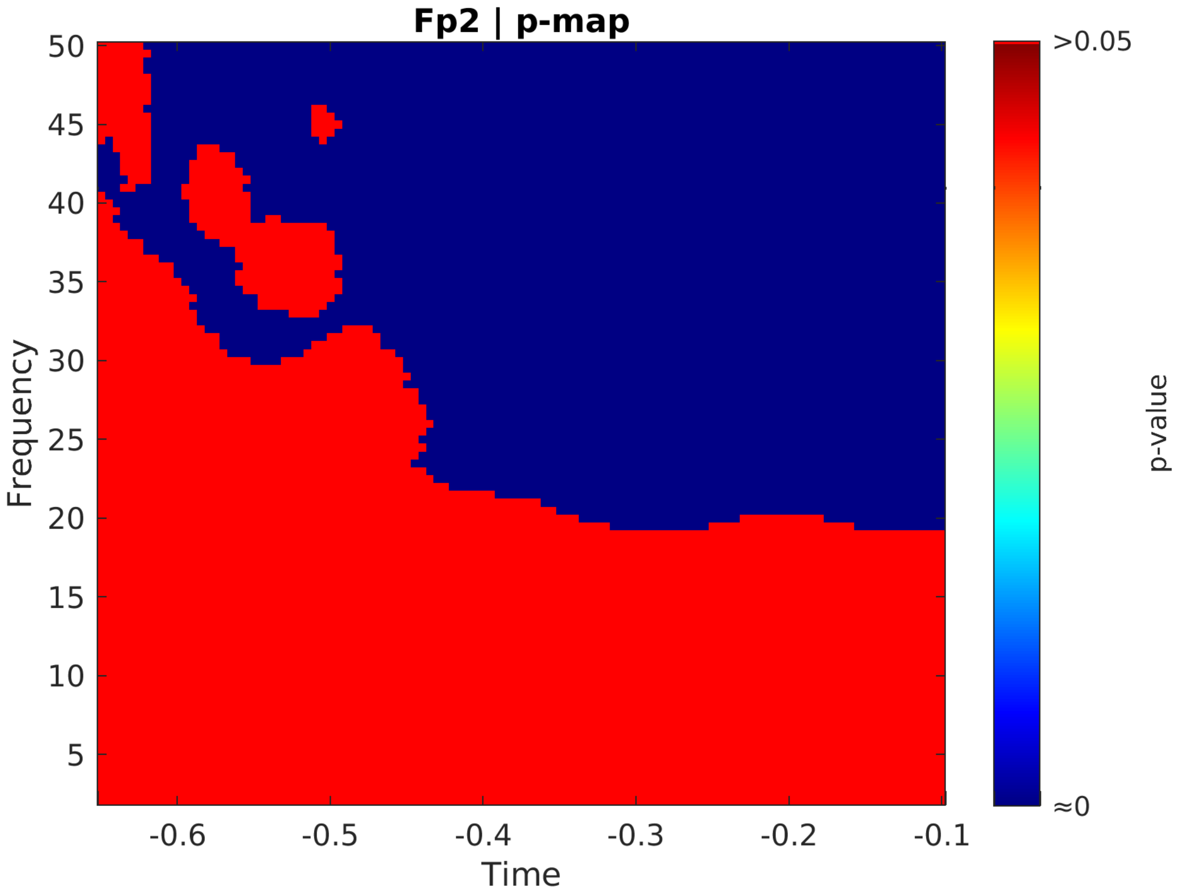
Fp2 | p-map
Frequency
Time
50
45
40
35
30
25
20
15
10
5
-0.6
-0.5
-0.4
-0.3
-0.2
-0.1
>0.05
≈0
p-value

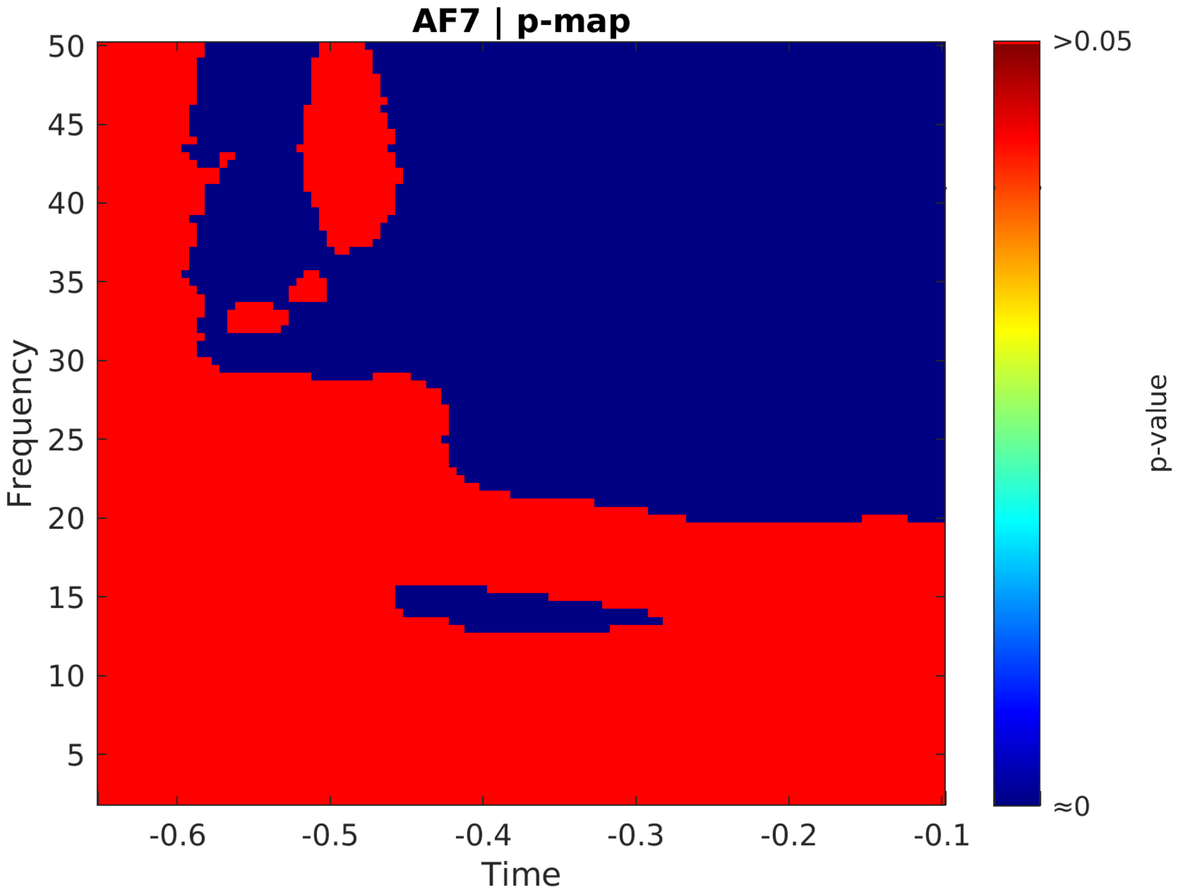
AF7 | p-map
Frequency
50
45
40
35
30
25
20
15
10
5
-0.6
-0.5
-0.4
-0.3
-0.2
-0.1
Time
>0.05
≈0
p-value

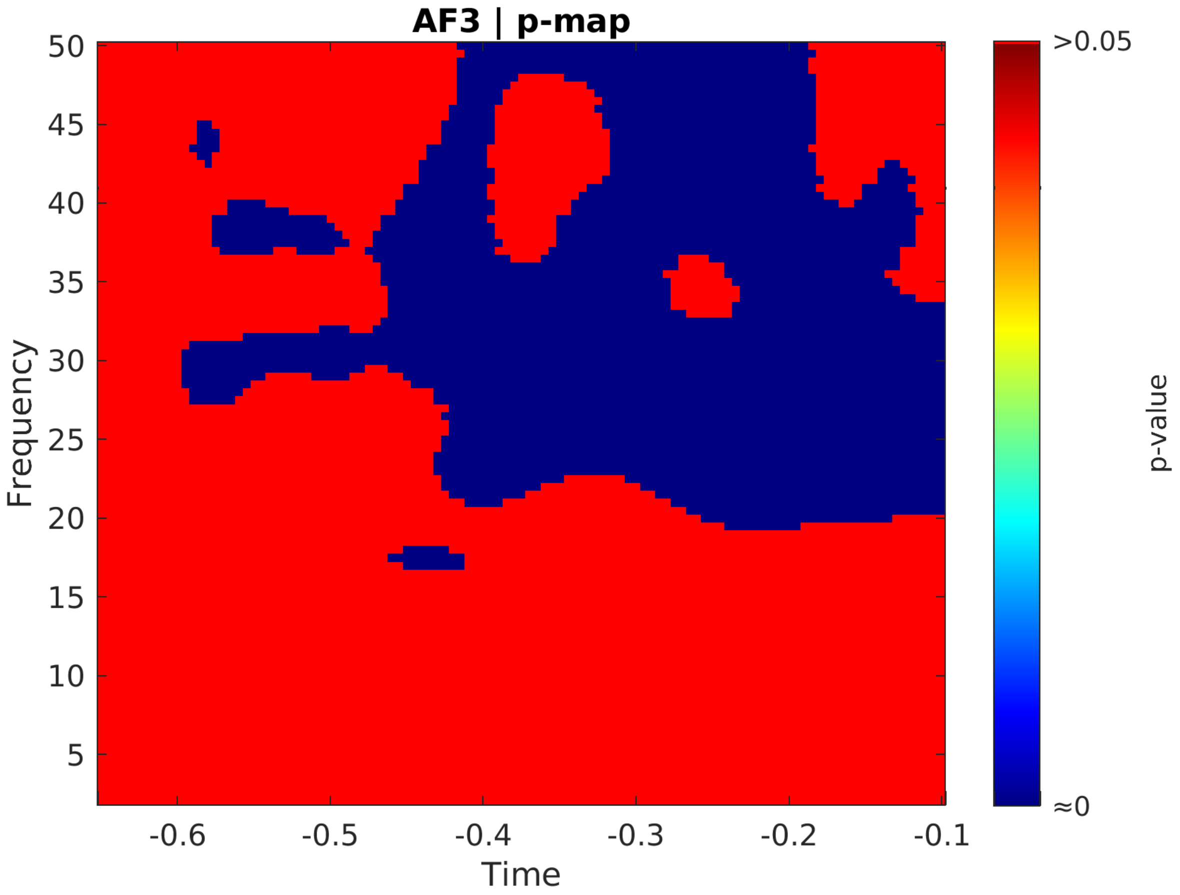

AF3 | p-map
Frequency
50
45
40
35
30
25
20
15
10
5
-0.6
-0.5
-0.4
-0.3
-0.2
-0.1
Time
>0.05
≈0
p-value

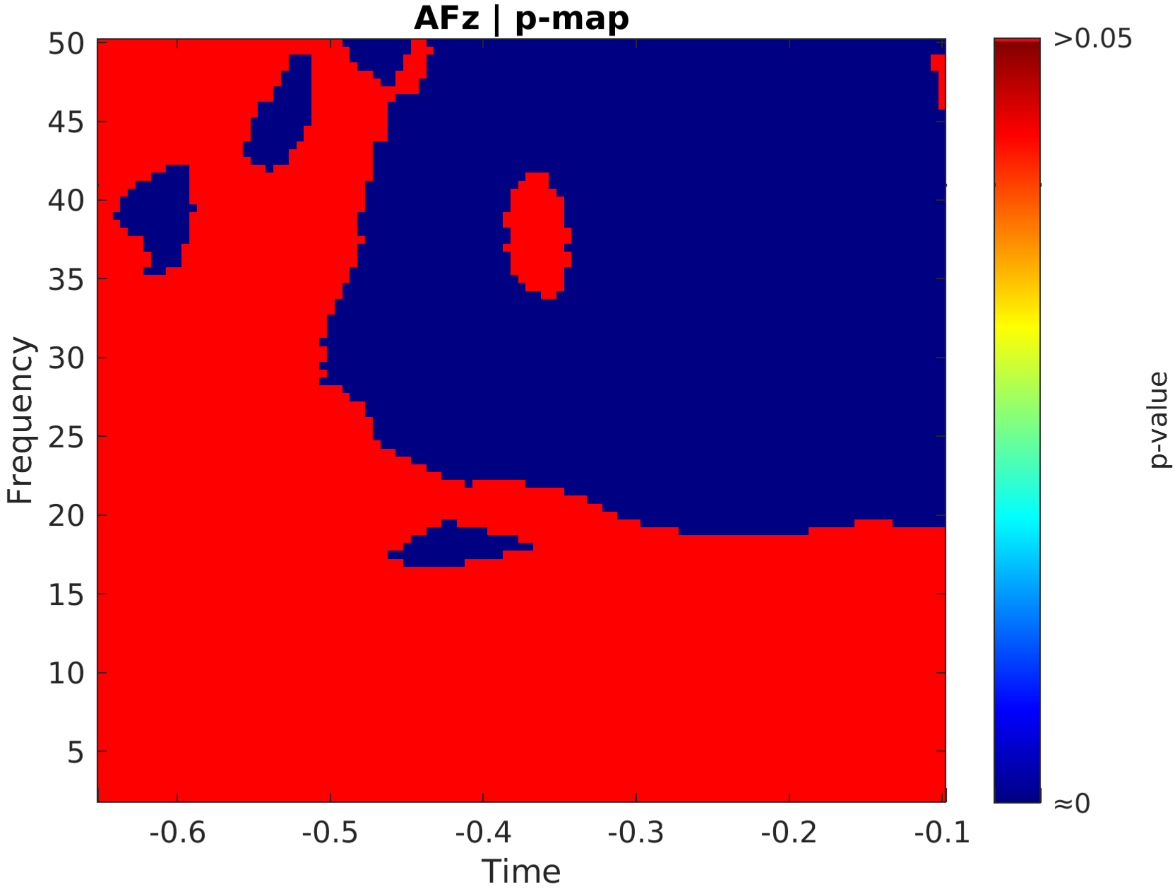
AFz | p-map
Frequency
50
45
40
35
30
25
20
15
10
5
-0.6
-0.5
-0.4
-0.3
-0.2
-0.1
Time
>0.05
≈0
p-value

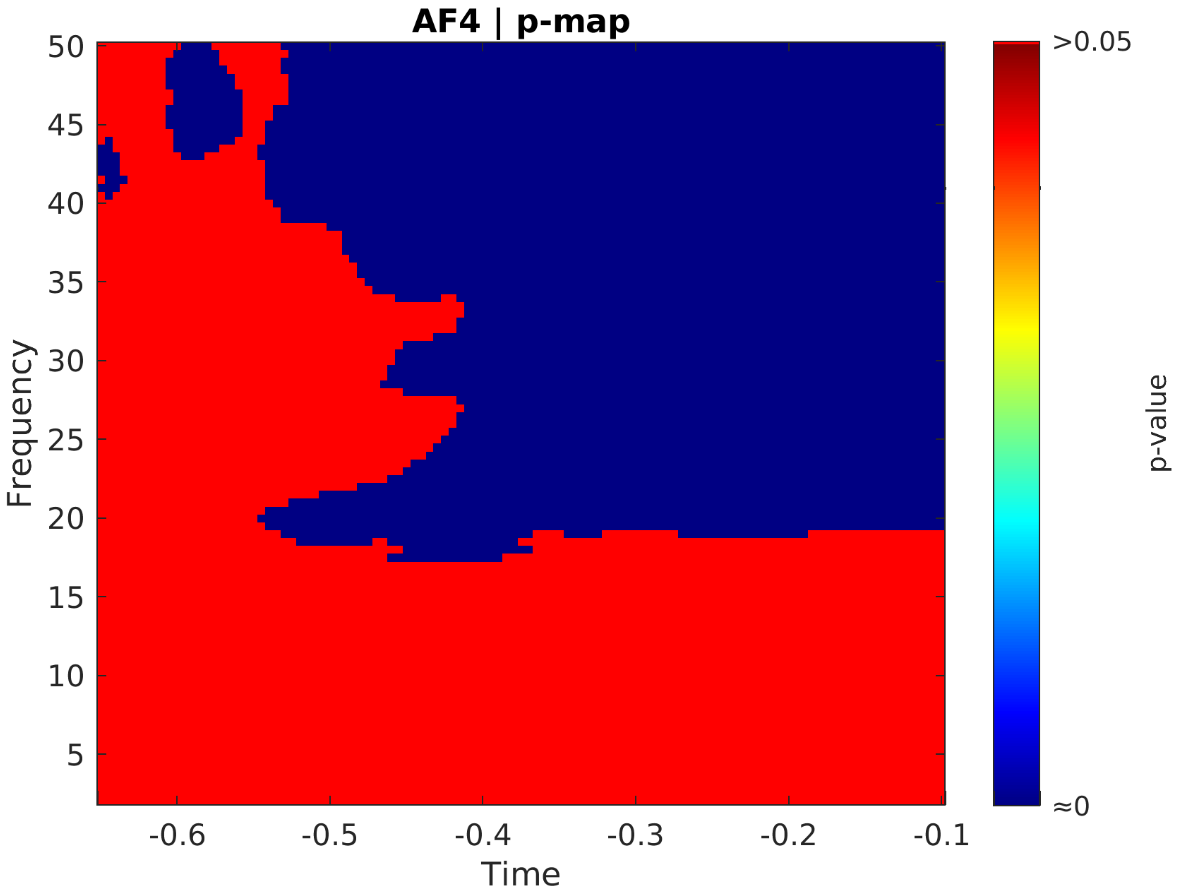

AF4 | p-map
Frequency
Time
p-value
>0.05
≈0
50
45
40
35
30
25
20
15
10
5
-0.6
-0.5
-0.4
-0.3
-0.2
-0.1

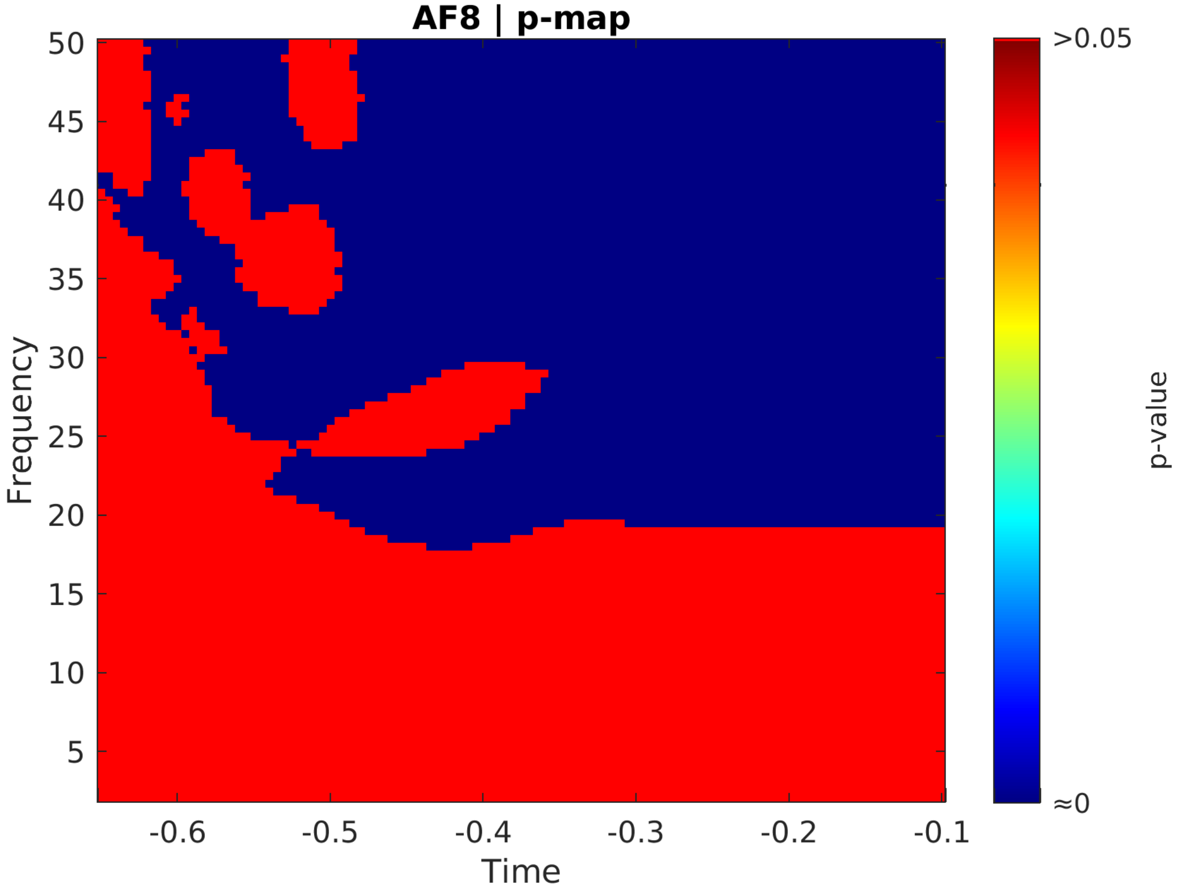
AF8 | p-map
Frequency
50
45
40
35
30
25
20
15
10
5
-0.6
-0.5
-0.4
-0.3
-0.2
-0.1
Time
>0.05
≈0
p-value

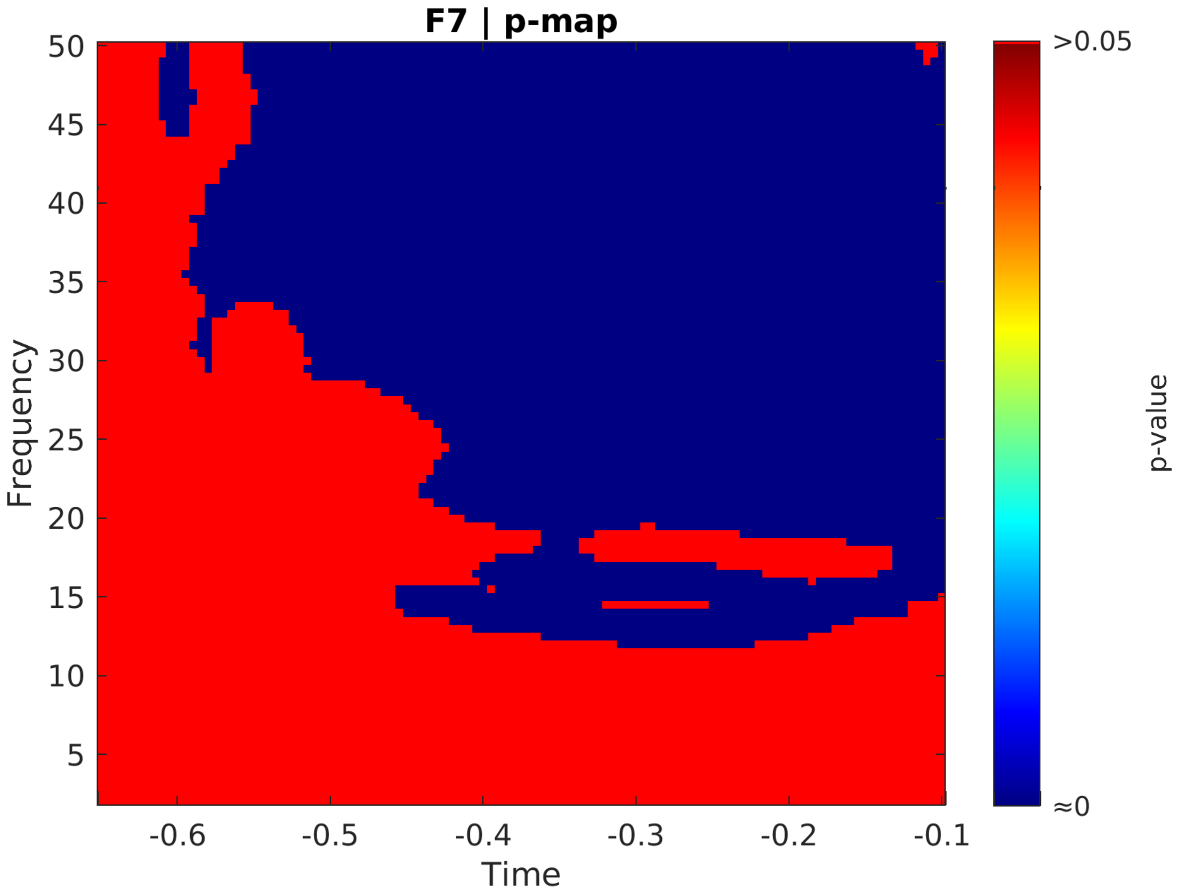
F7 | p-map
Frequency
50
45
40
35
30
25
20
15
10
5
-0.6
-0.5
-0.4
-0.3
-0.2
-0.1
Time
>0.05
≈0
p-value

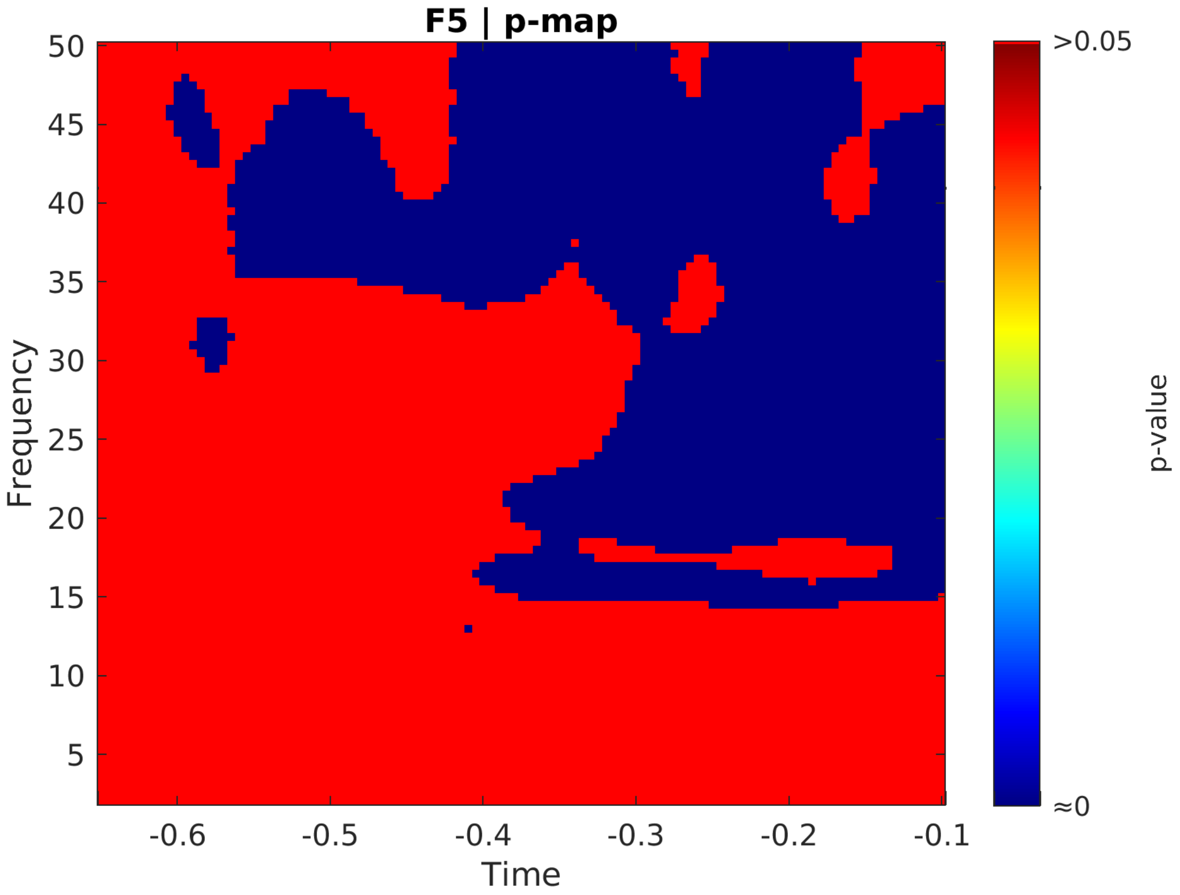
F5 | p-map
Frequency
50
45
40
35
30
25
20
15
10
5
-0.6
-0.5
-0.4
-0.3
-0.2
-0.1
Time
>0.05
≈0
p-value

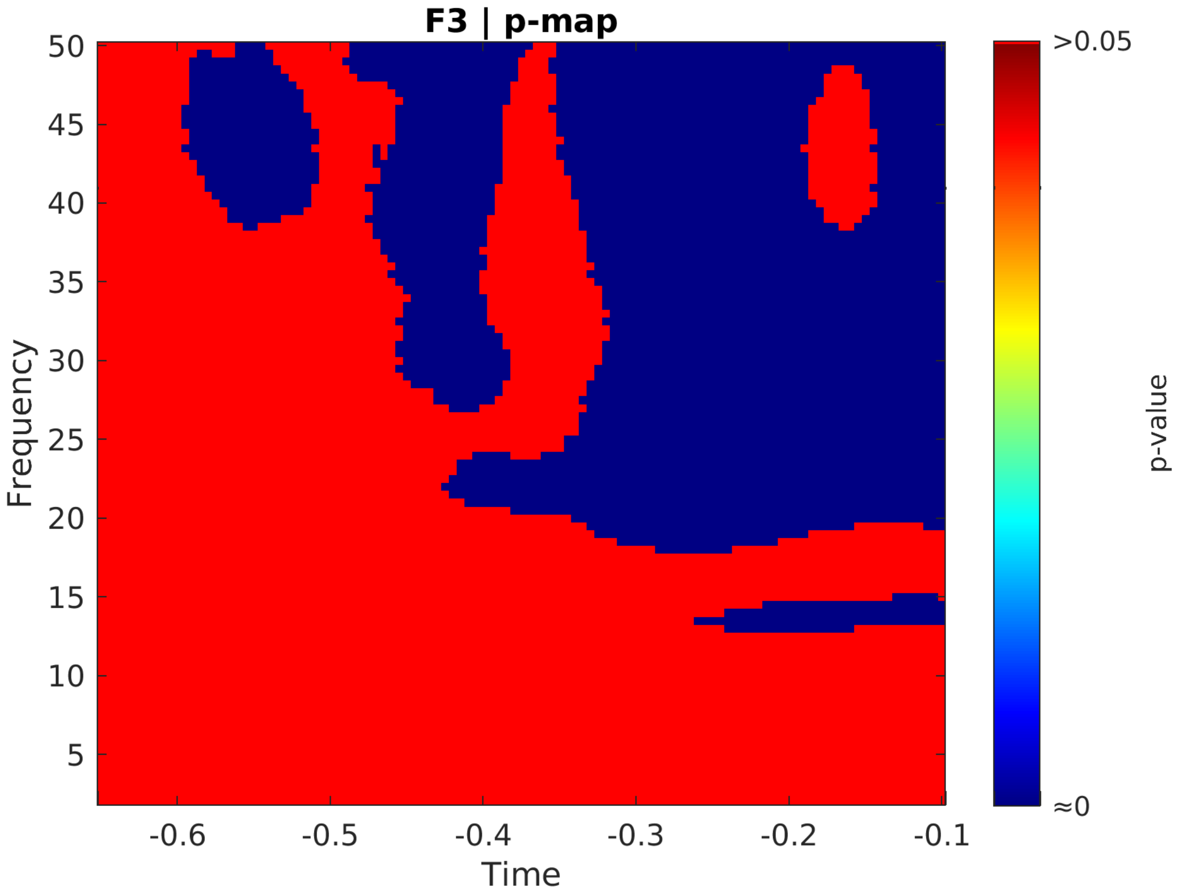

F3 | p-map
Frequency
50
45
40
35
30
25
20
15
10
5
-0.6
-0.5
-0.4
-0.3
-0.2
-0.1
Time
>0.05
≈0
p-value

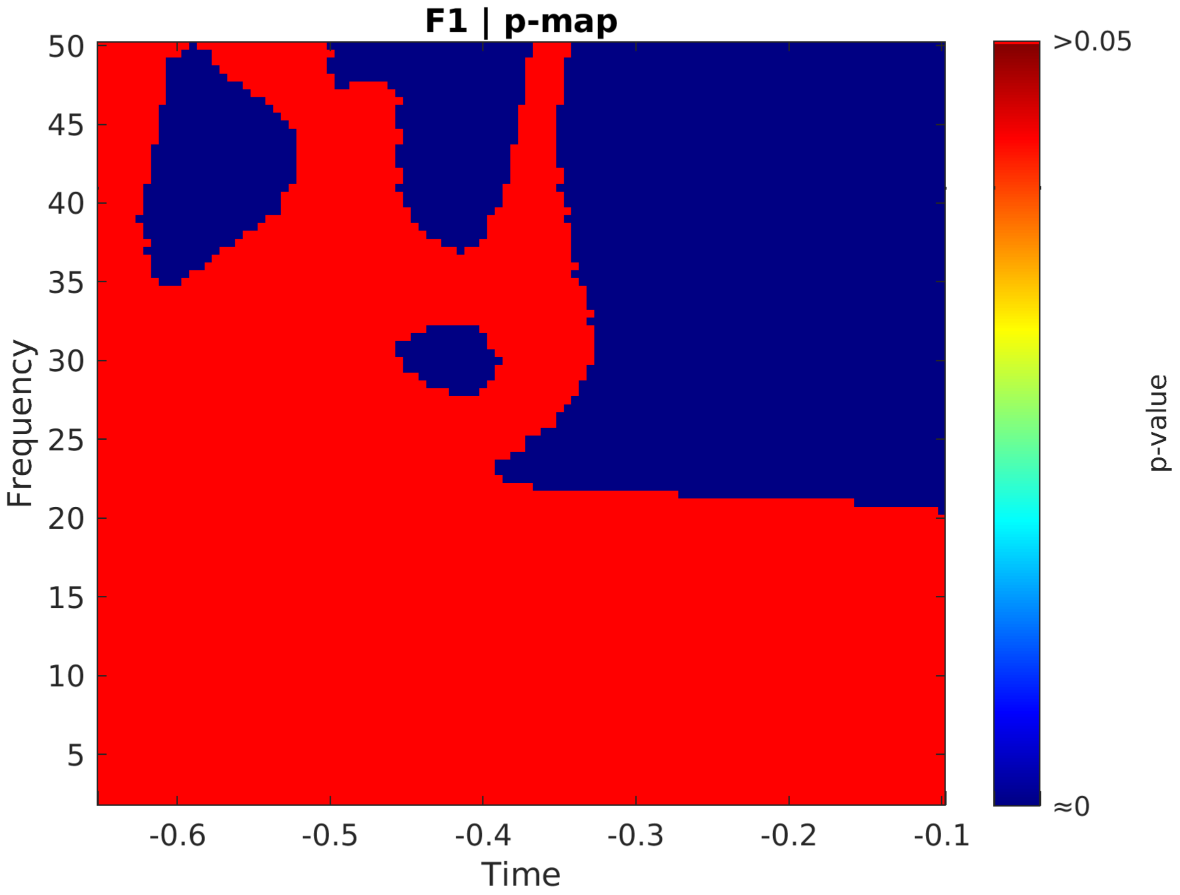
F1 | p-map
Frequency
50
45
40
35
30
25
20
15
10
5
Time
-0.6
-0.5
-0.4
-0.3
-0.2
-0.1
>0.05
≈0
p-value

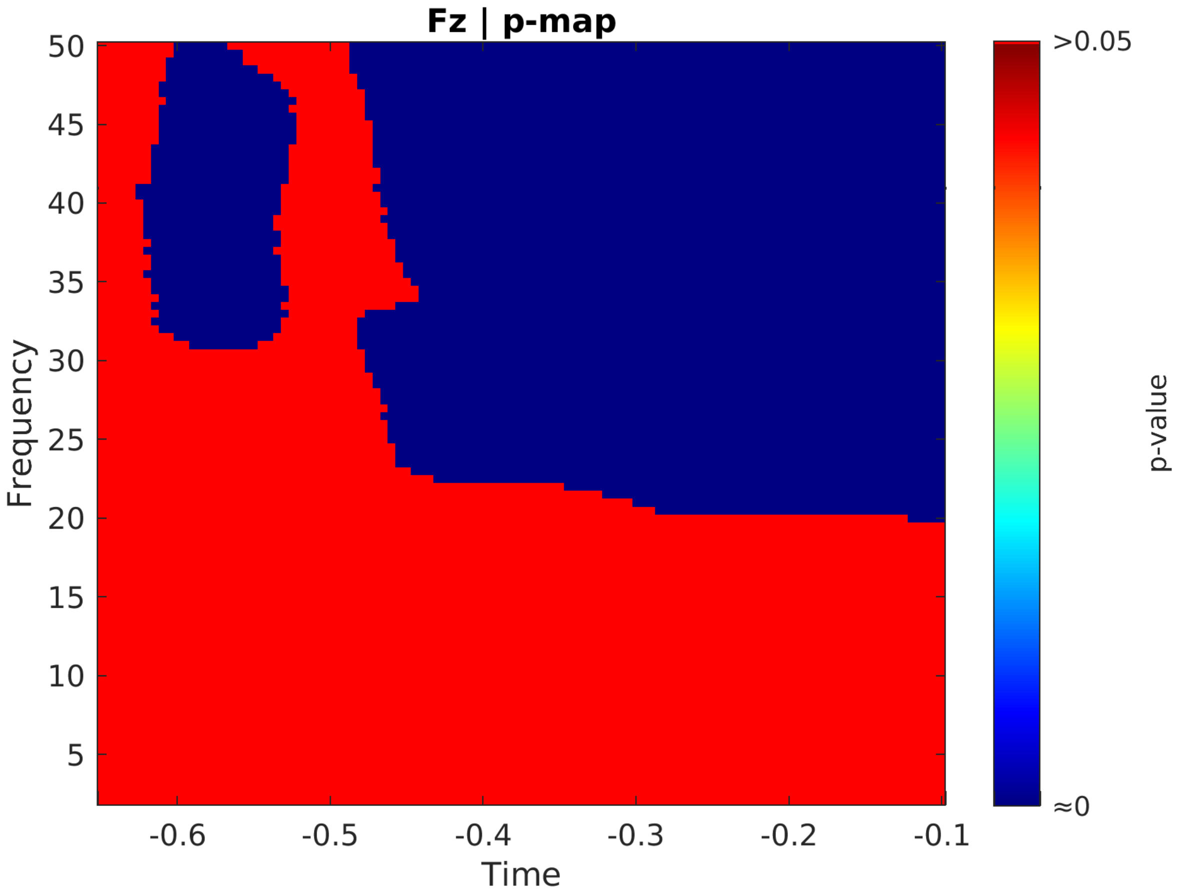

Fz | p-map
50
45
40
35
30
25
20
15
10
5
Frequency
-0.6
-0.5
-0.4
-0.3
-0.2
-0.1
Time
>0.05
≈0
p-value

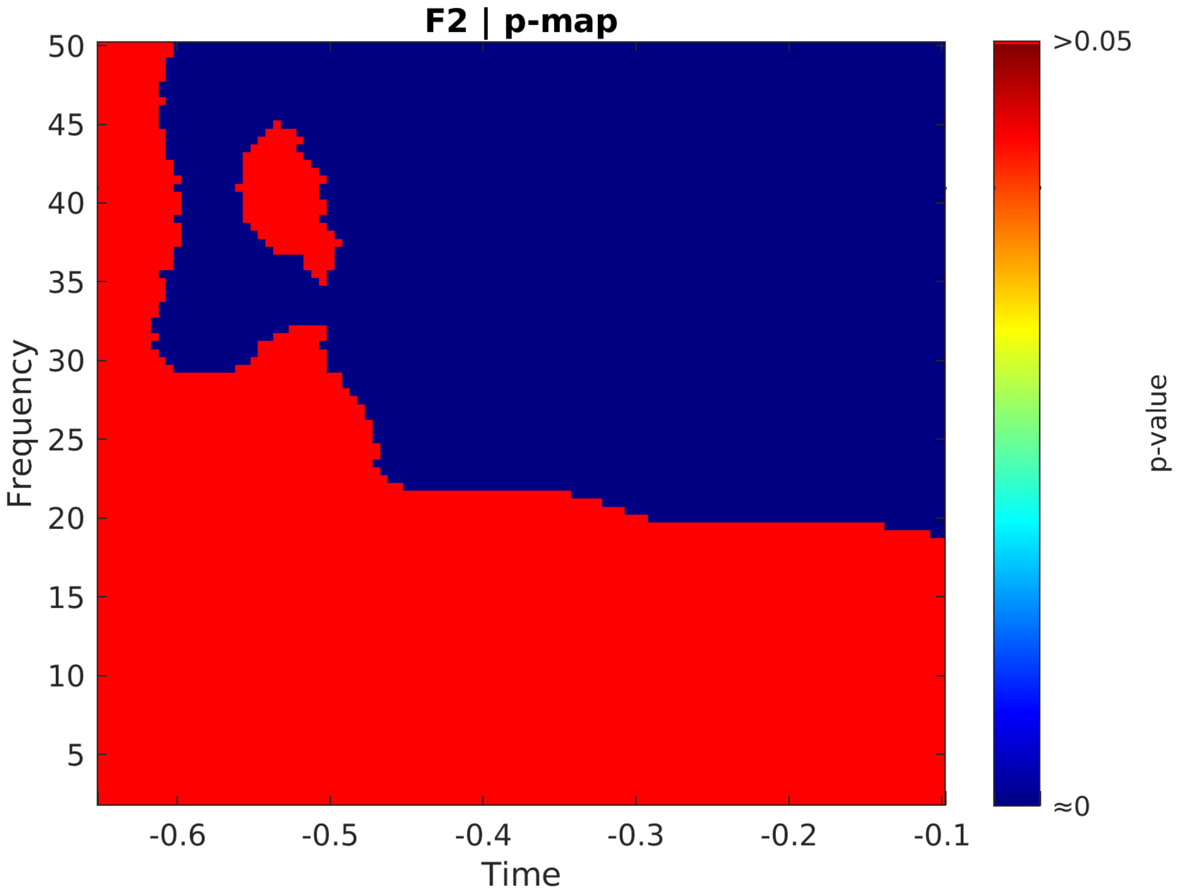
F2 | p-map
Frequency
50
45
40
35
30
25
20
15
10
5
-0.6
-0.5
-0.4
-0.3
-0.2
-0.1
Time
>0.05
≈0
p-value

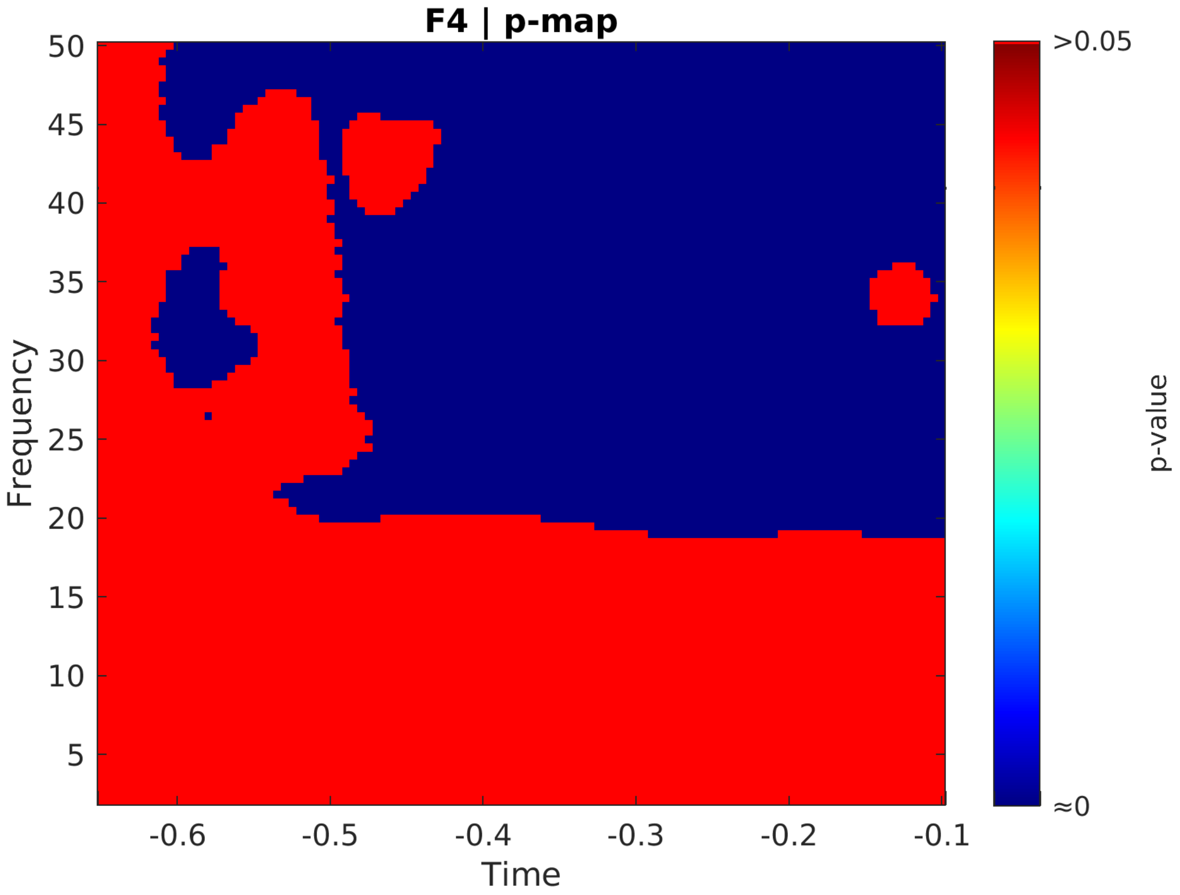
F4 | p-map
Frequency
Time
p-value
>0.05
≈0
50
45
40
35
30
25
20
15
10
5
-0.6
-0.5
-0.4
-0.3
-0.2
-0.1

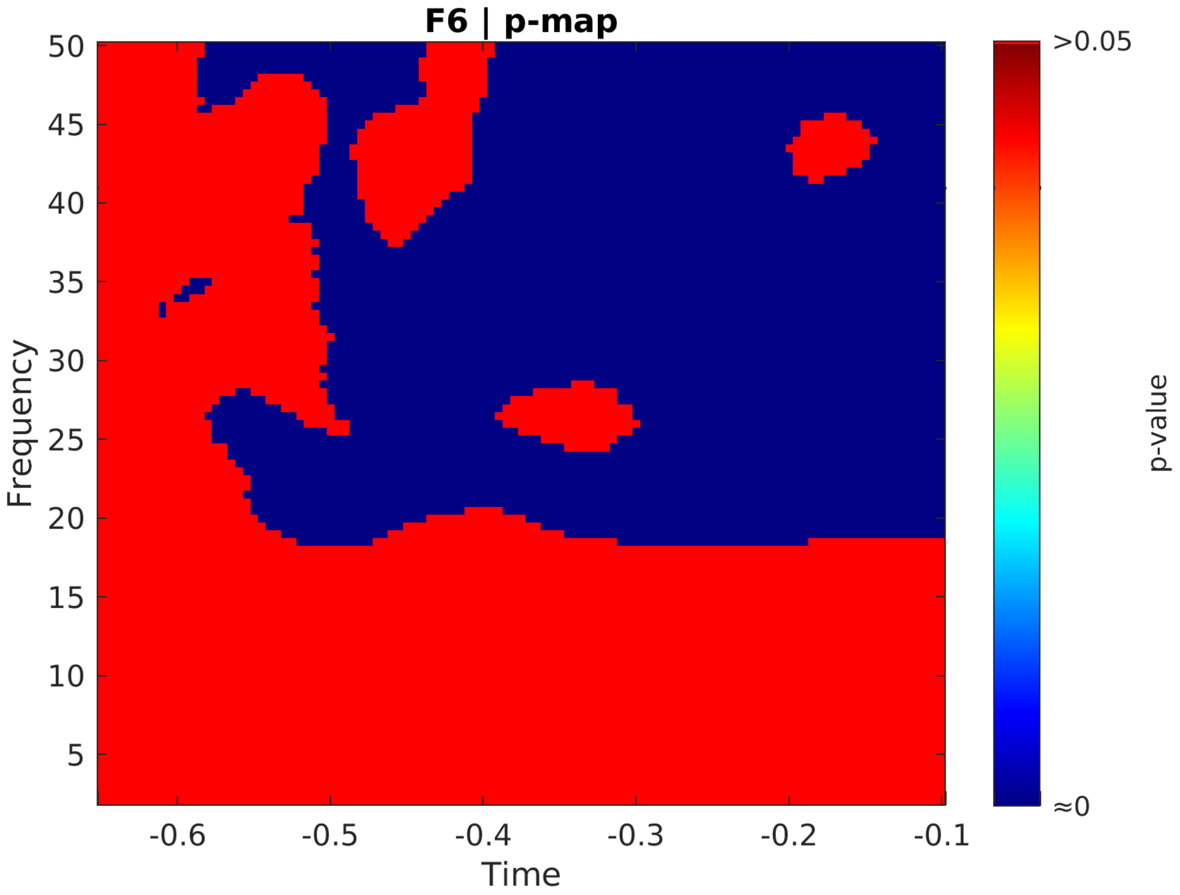
F6 | p-map
Frequency
Time
p-value
>0.05
≈0
50
45
40
35
30
25
20
15
10
5
-0.6
-0.5
-0.4
-0.3
-0.2
-0.1

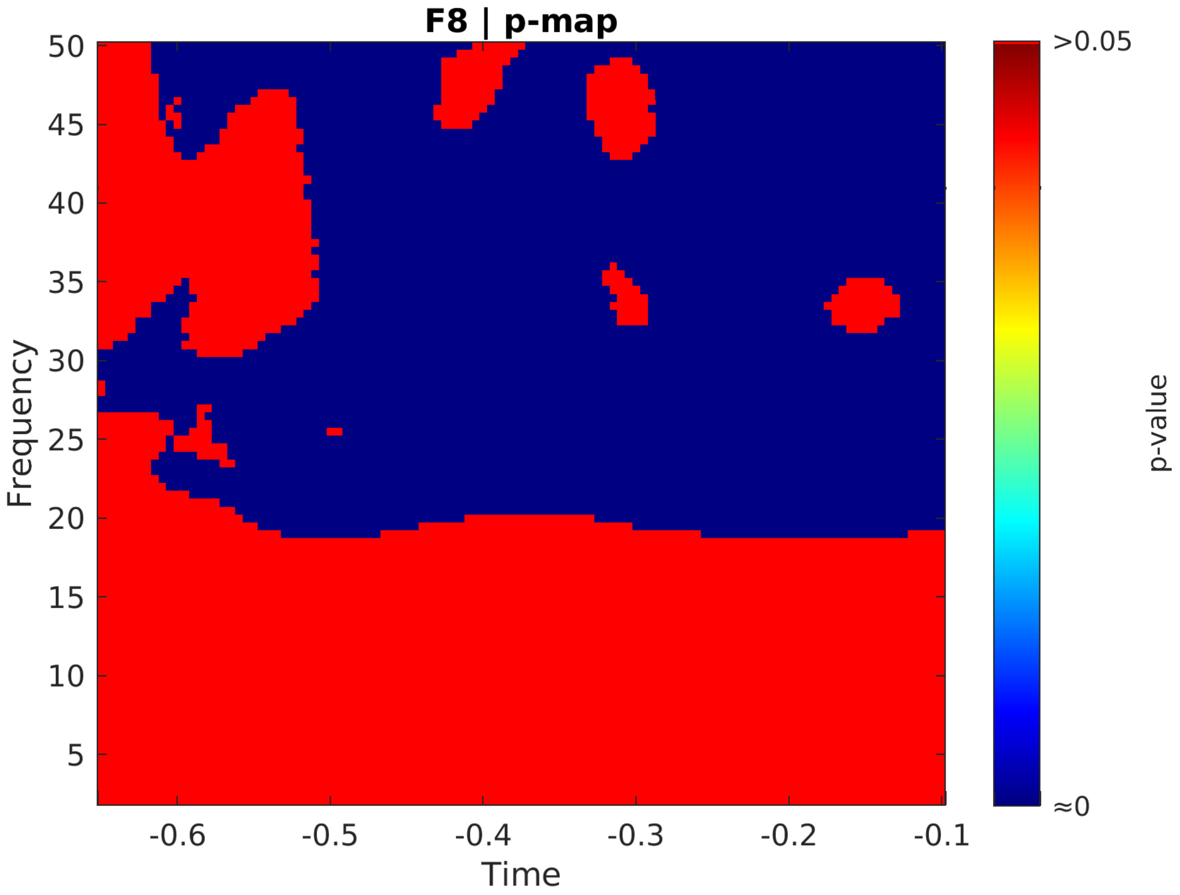

F8 | p-map
50
45
40
35
30
25
20
15
10
5
Frequency
-0.6
-0.5
-0.4
-0.3
-0.2
-0.1
Time
>0.05
≈0
p-value

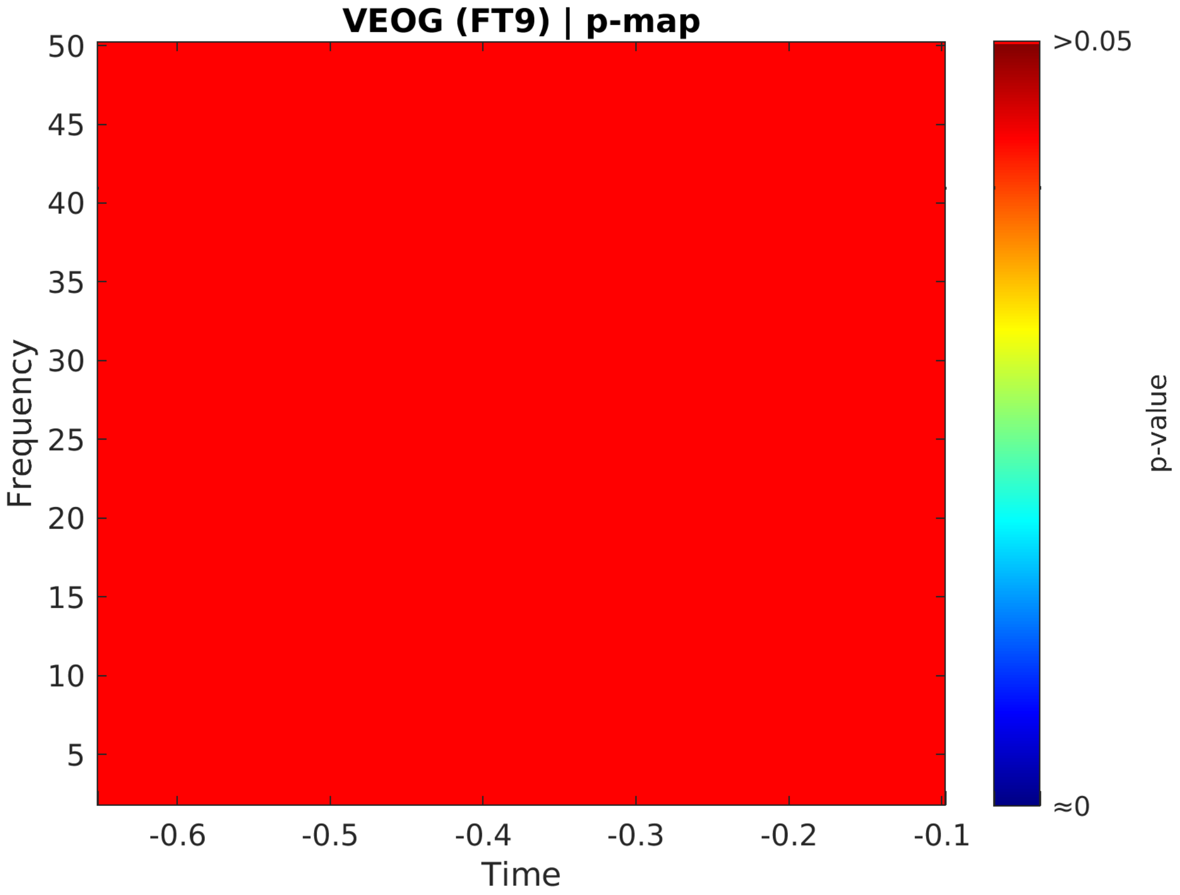

VEOG (FT9) | p-map
50
45
40
35
30
25
20
15
10
5
Frequency
-0.6
-0.5
-0.4
-0.3
-0.2
-0.1
Time
>0.05
≈0
p-value

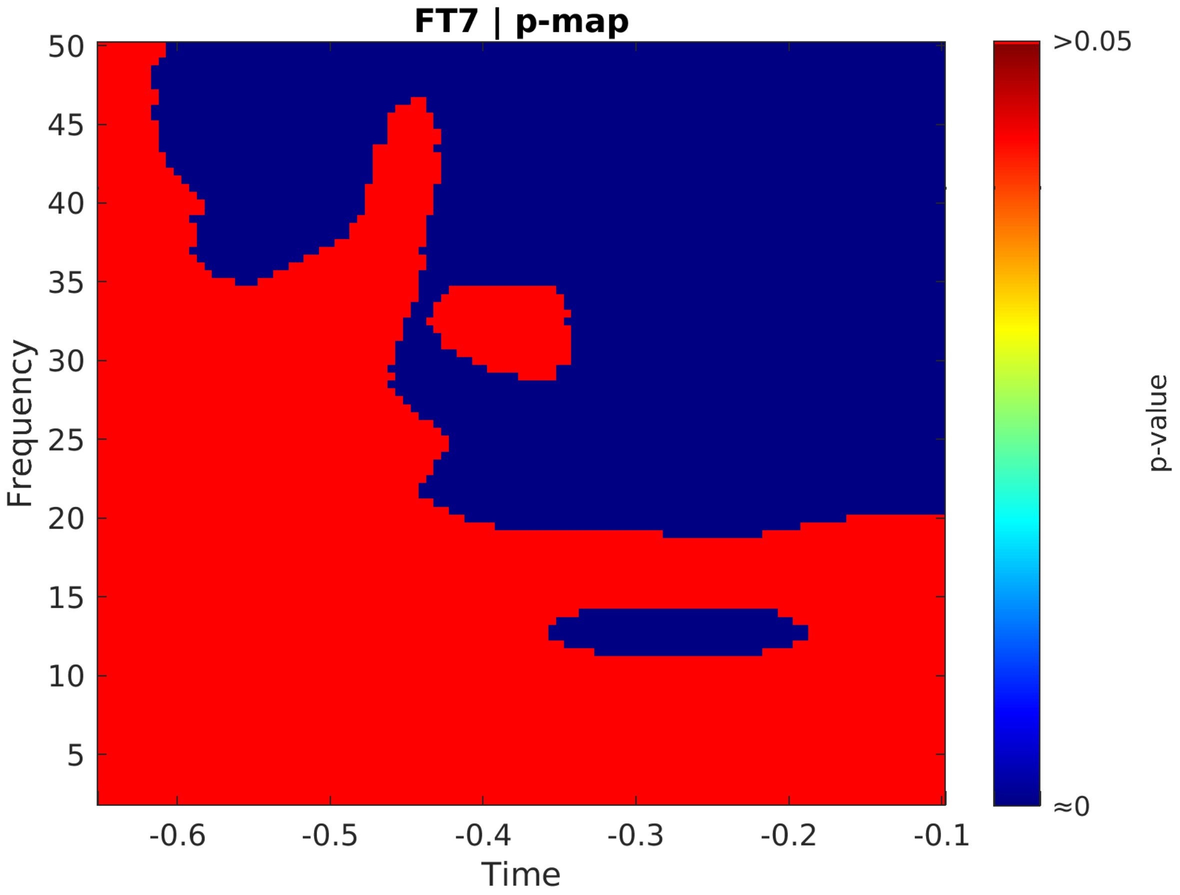

FT7 | p-map
Frequency
50
45
40
35
30
25
20
15
10
5
-0.6
-0.5
-0.4
-0.3
-0.2
-0.1
Time
>0.05
≈0
p-value

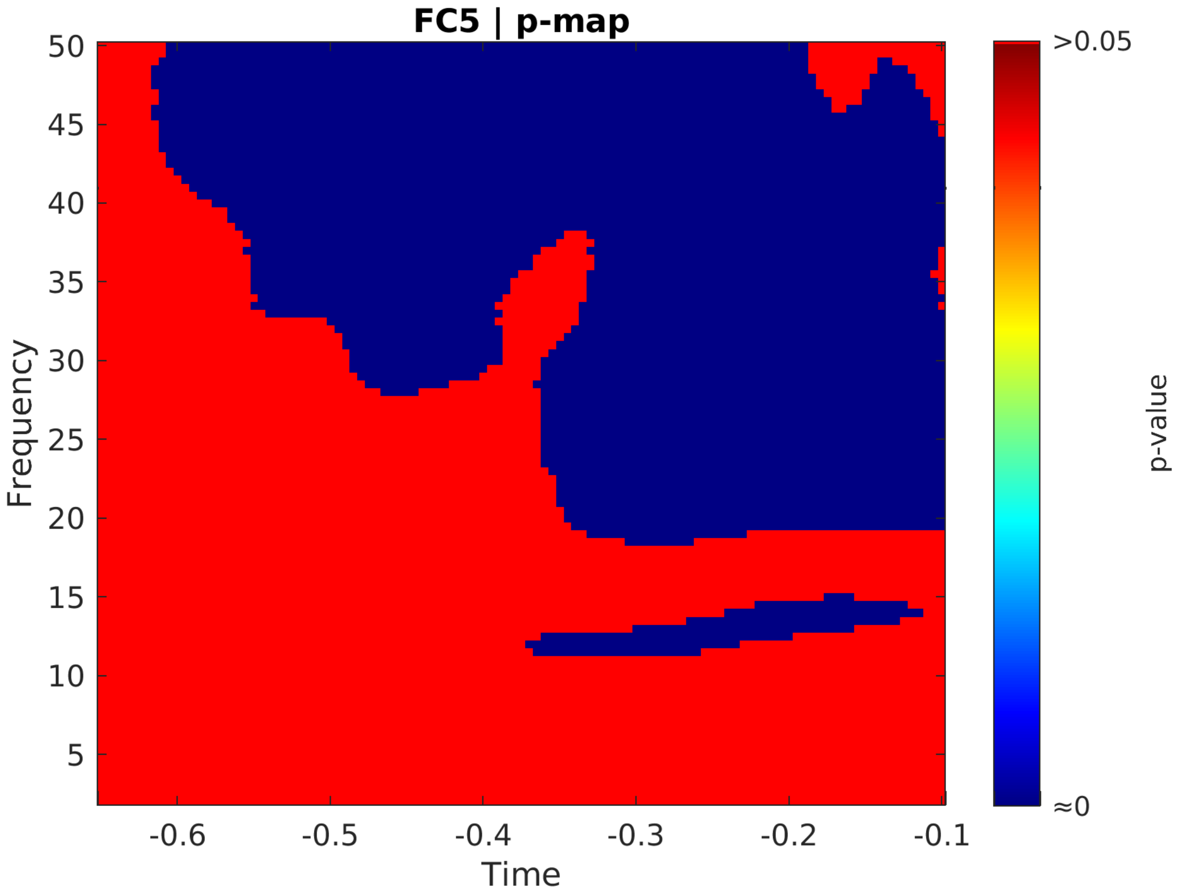
FC5 | p-map
50
45
40
35
30
25
20
15
10
5
Frequency
-0.6
-0.5
-0.4
-0.3
-0.2
-0.1
Time
>0.05
≈0
p-value

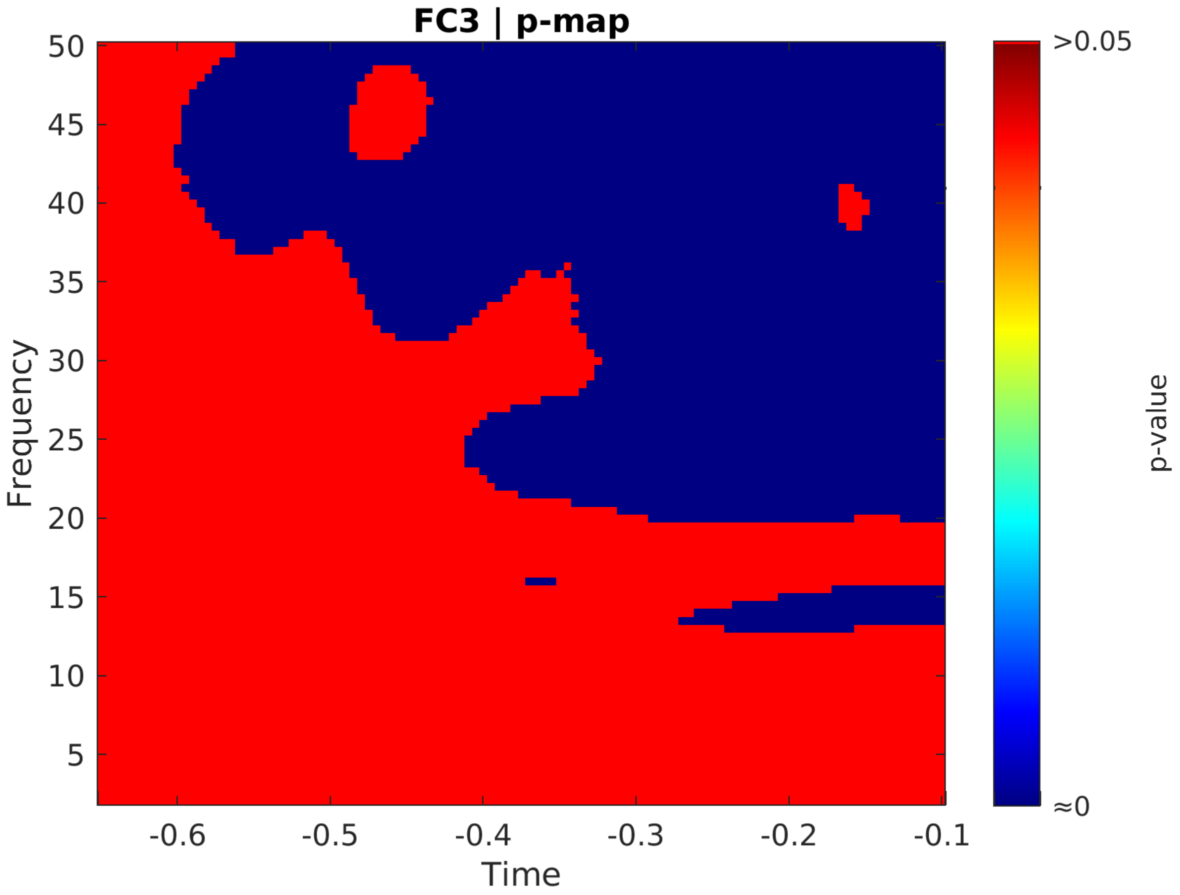

FC3 | p-map
50
45
40
35
30
25
20
15
10
5
Frequency
-0.6
-0.5
-0.4
-0.3
-0.2
-0.1
Time
>0.05
≈0
p-value

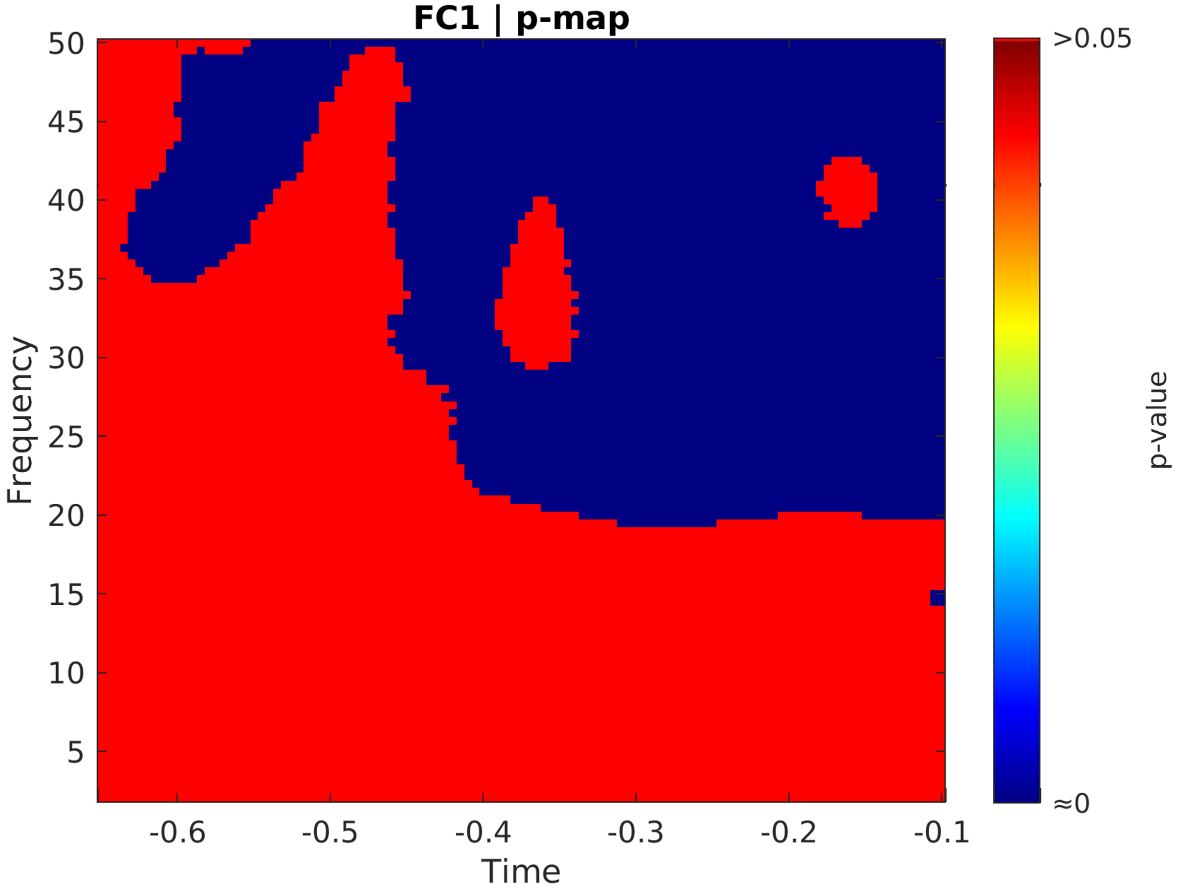
FC1 | p-map
Frequency
Time
p-value
>0.05
≈0
50
45
40
35
30
25
20
15
10
5
-0.6
-0.5
-0.4
-0.3
-0.2
-0.1

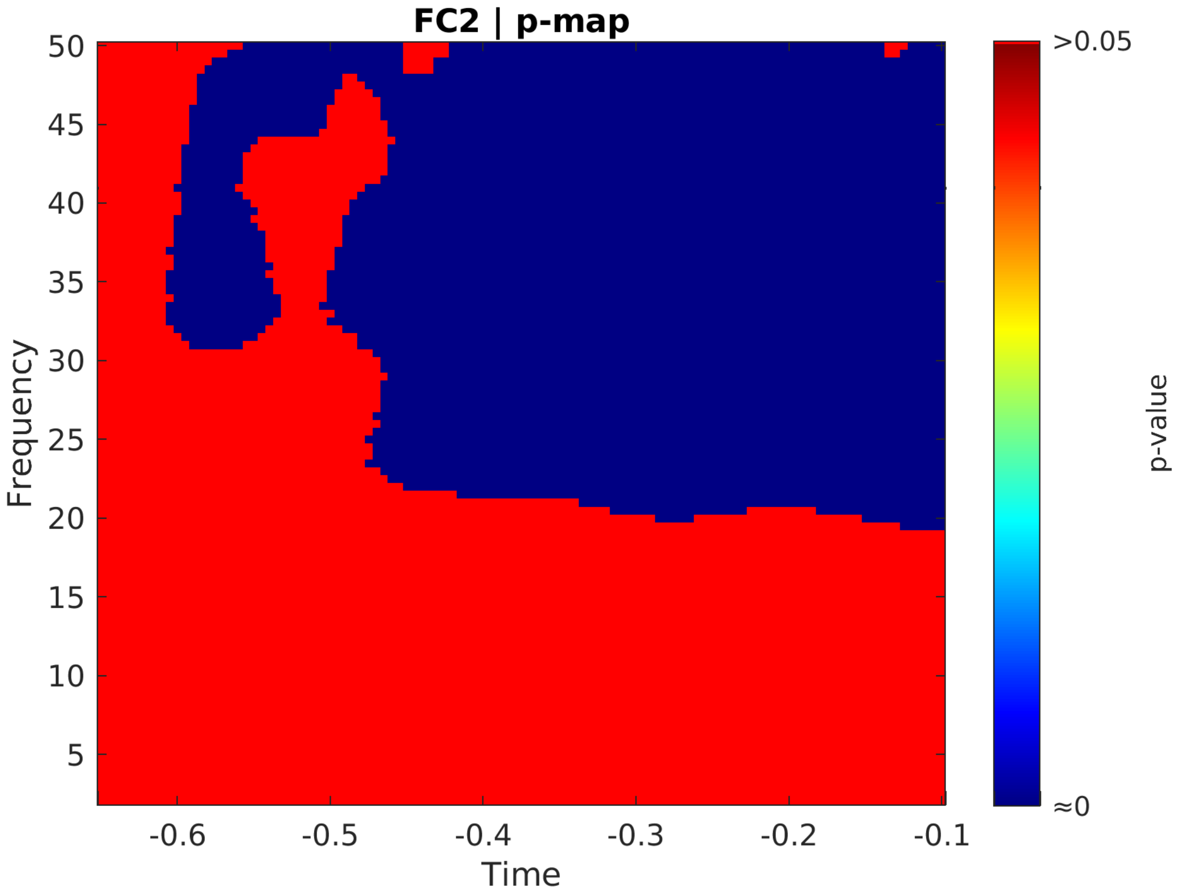
FC2 | p-map
Frequency
Time
p-value
>0.05
≈0
50
45
40
35
30
25
20
15
10
5
-0.6
-0.5
-0.4
-0.3
-0.2
-0.1

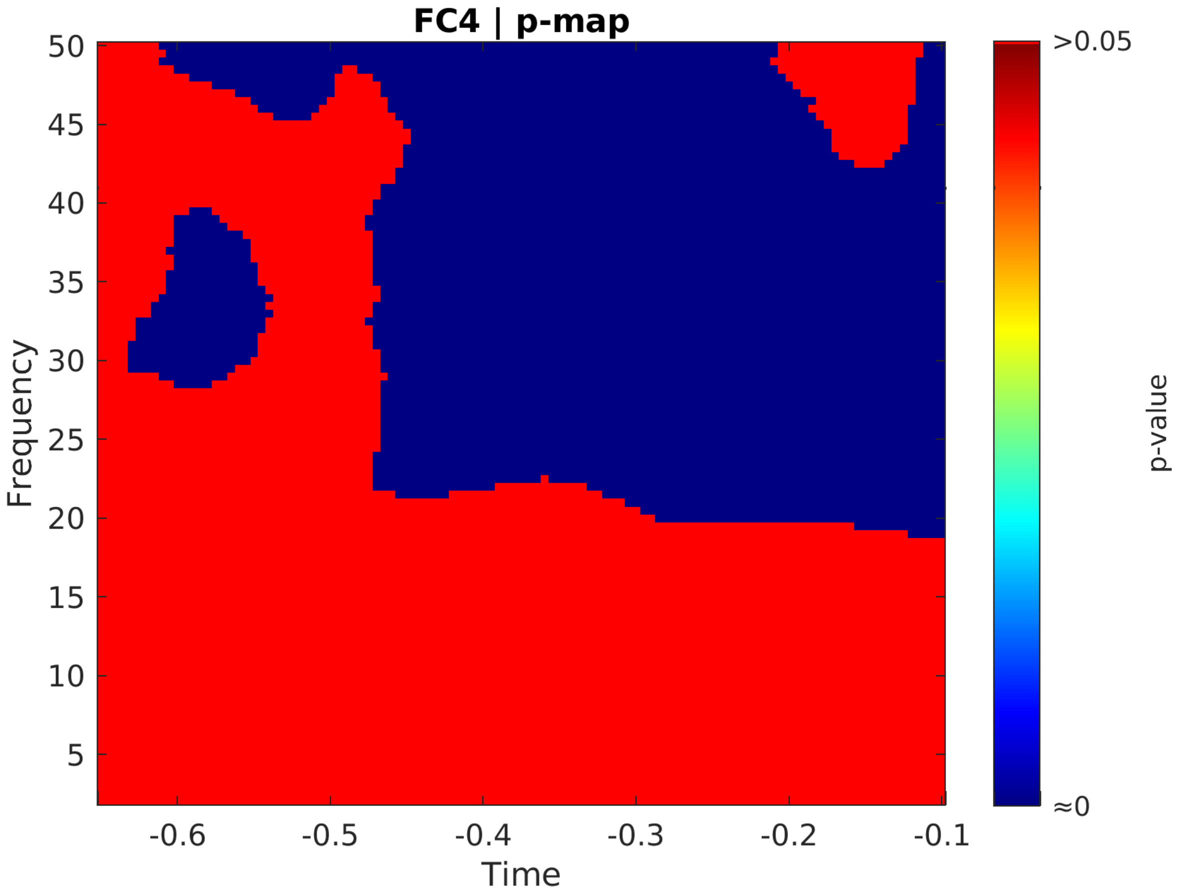
FC4 | p-map
Frequency
Time
p-value
>0.05
≈0
50
45
40
35
30
25
20
15
10
5
-0.6
-0.5
-0.4
-0.3
-0.2
-0.1

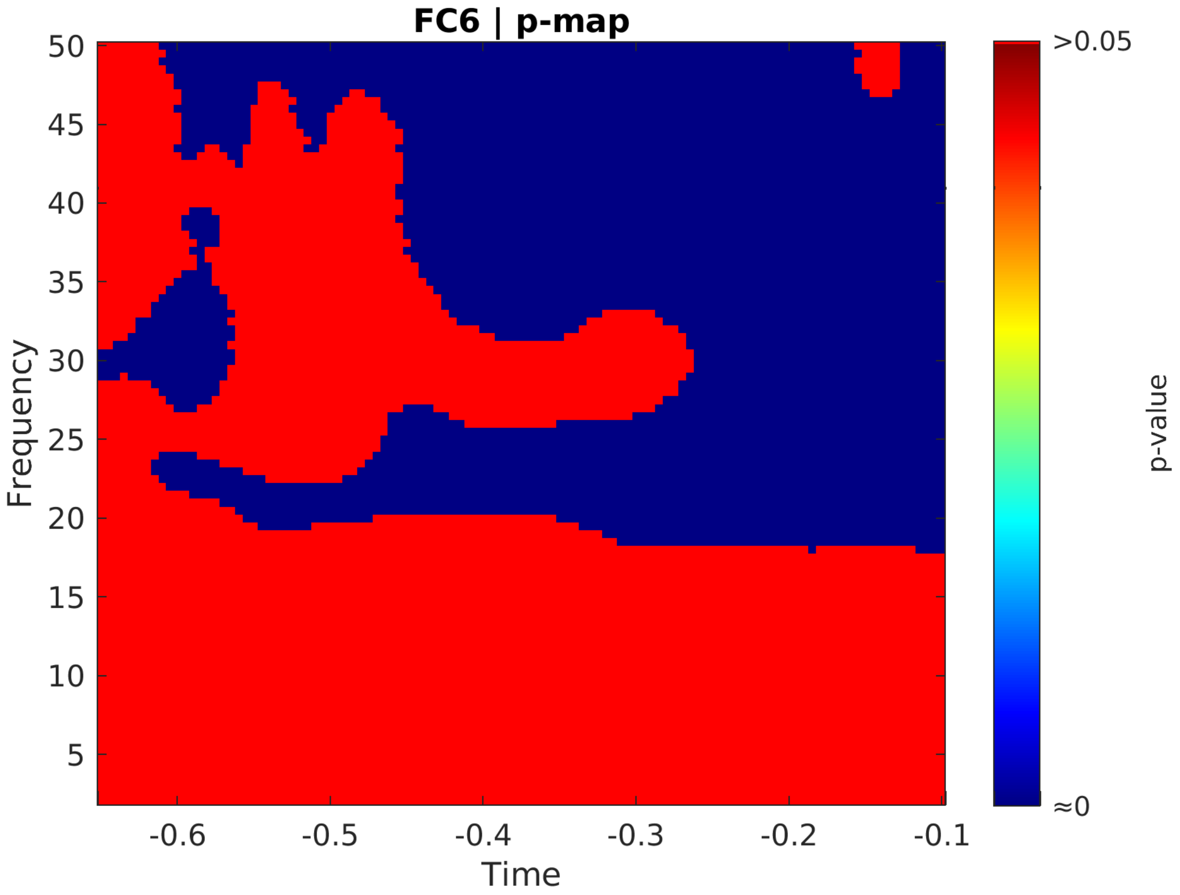
FC6 | p-map
Frequency
Time
50
45
40
35
30
25
20
15
10
5
-0.6
-0.5
-0.4
-0.3
-0.2
-0.1
>0.05
≈0
p-value

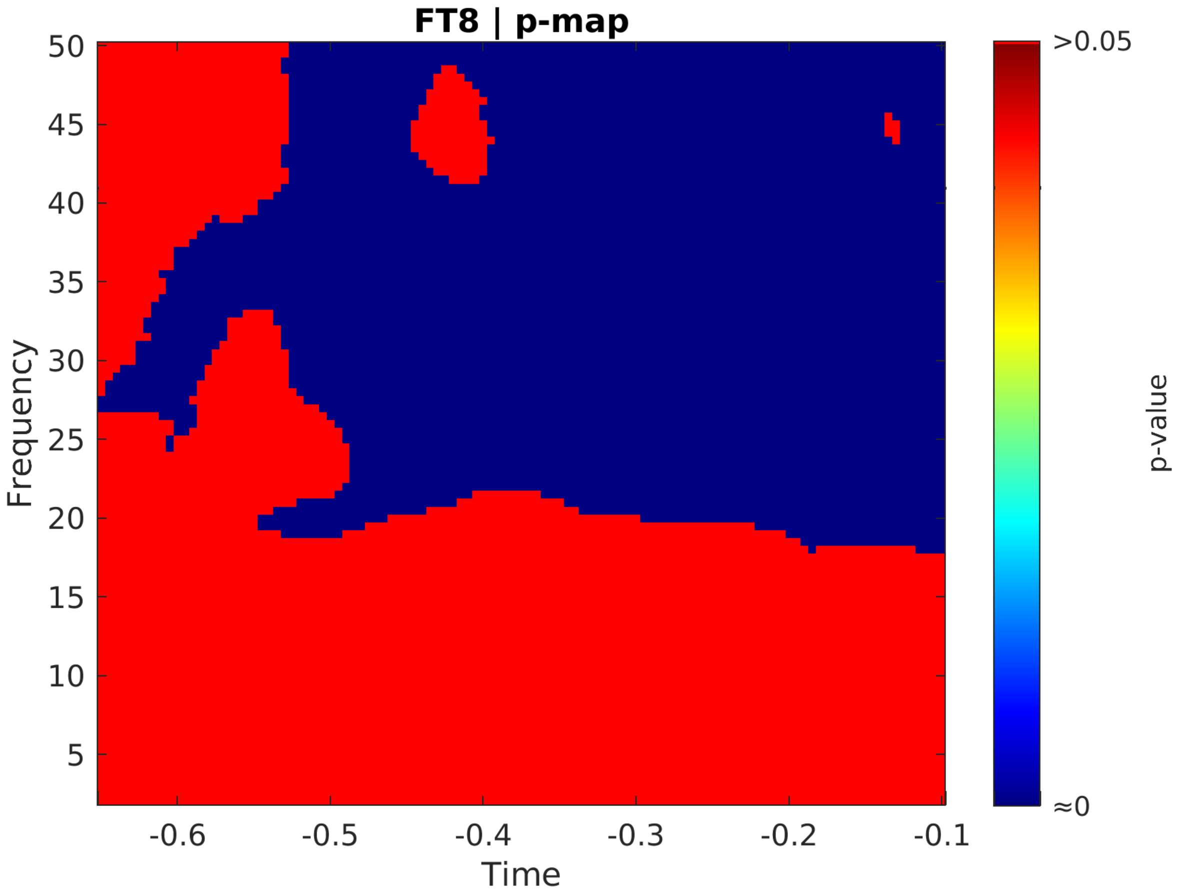
FT8 | p-map
Frequency
50
45
40
35
30
25
20
15
10
5
-0.6
-0.5
-0.4
-0.3
-0.2
-0.1
Time
>0.05
≈0
p-value

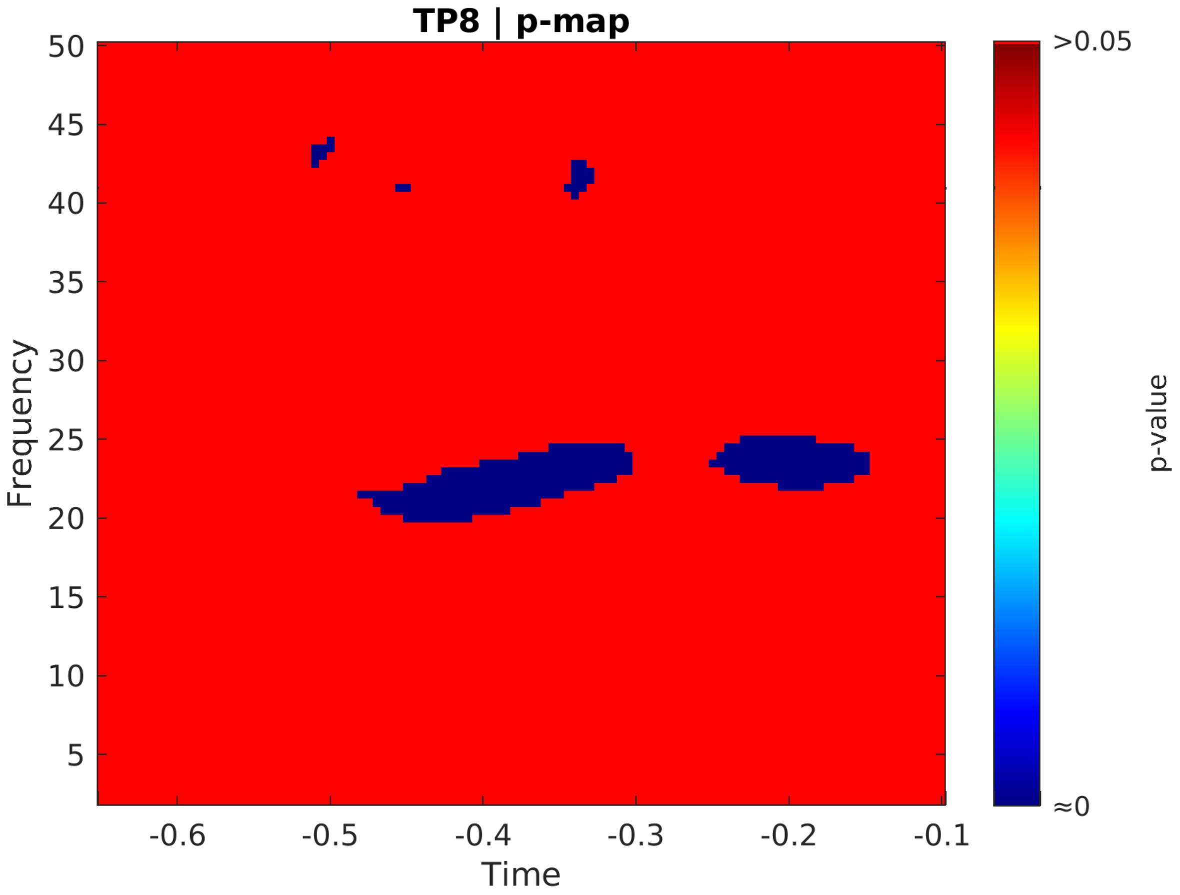
TP8 | p-map
50
45
40
35
30
25
20
15
10
5
Frequency
-0.6
-0.5
-0.4
-0.3
-0.2
-0.1
Time
>0.05
≈0
p-value

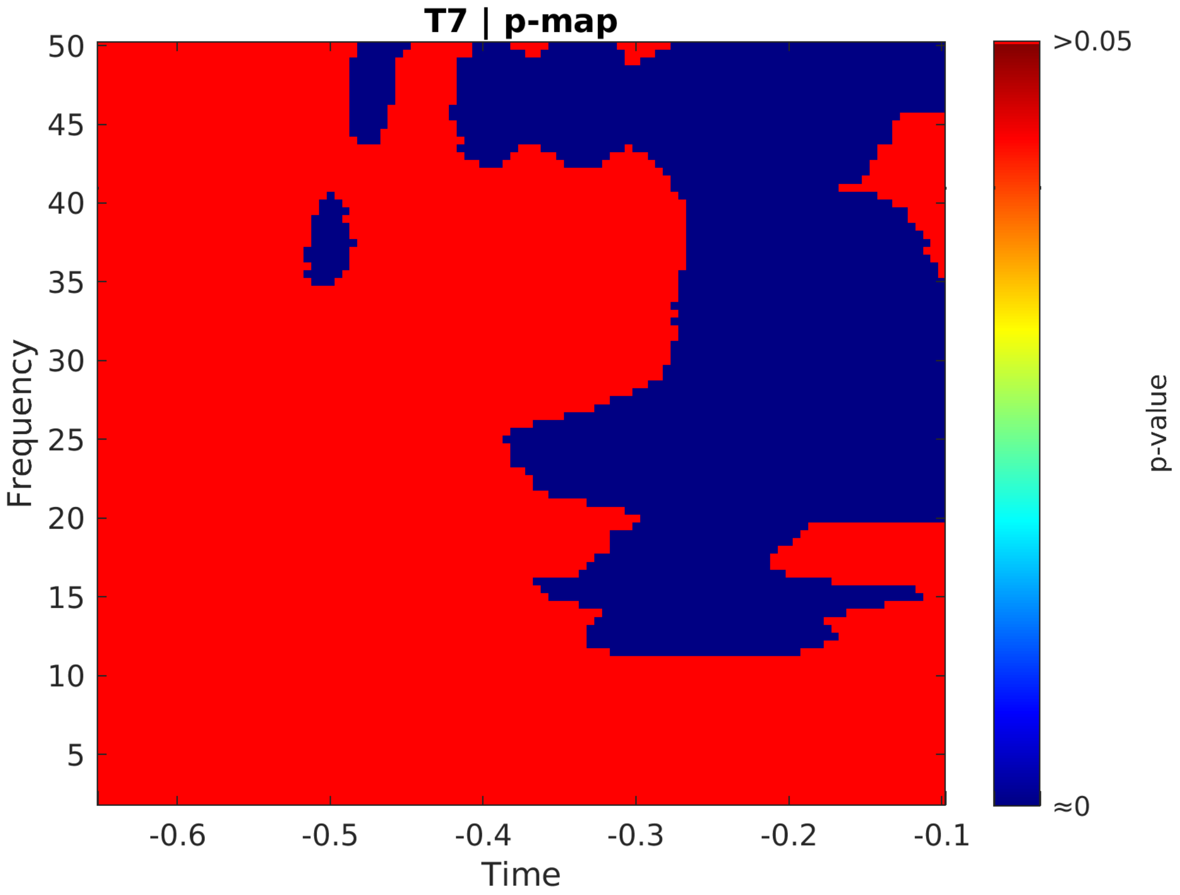
T7 | p-map
Frequency
Time
p-value
>0.05
≈0
50
45
40
35
30
25
20
15
10
5
-0.6
-0.5
-0.4
-0.3
-0.2
-0.1

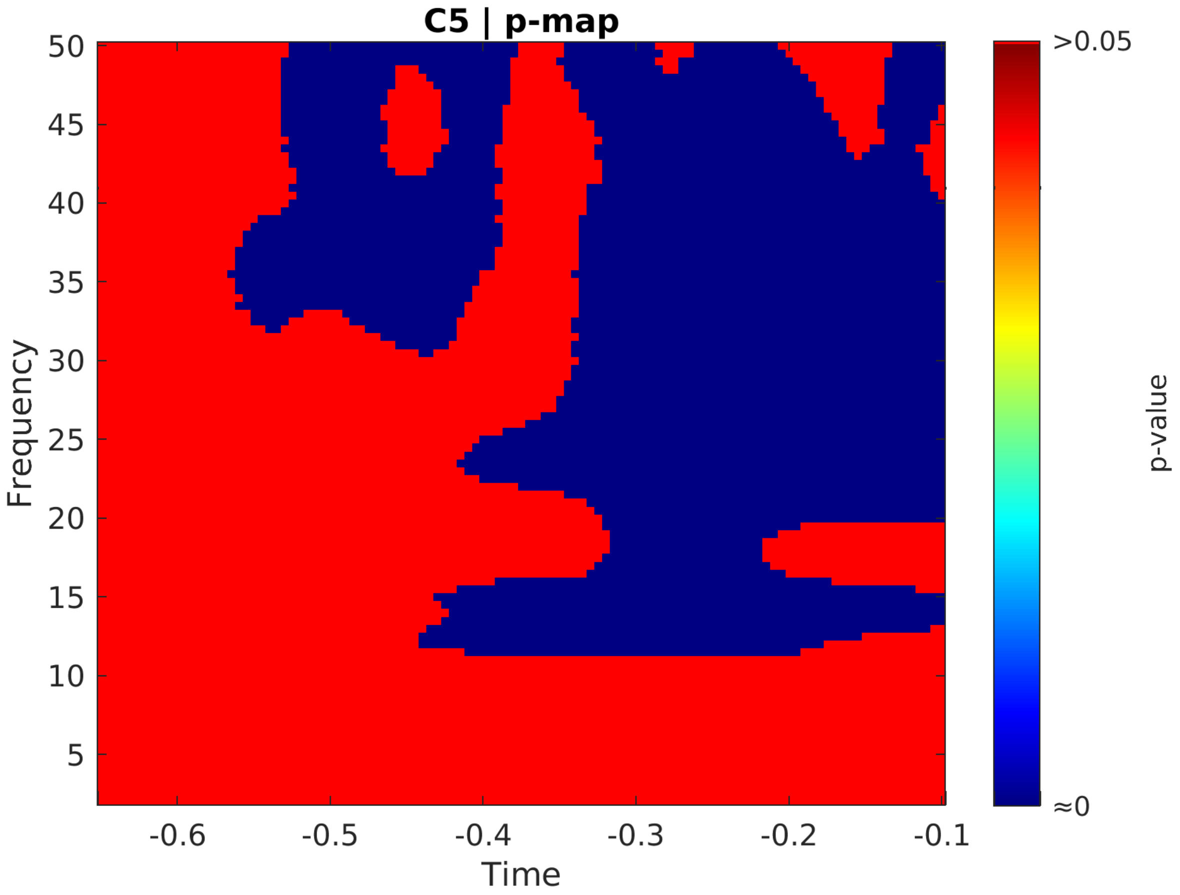
C5 | p-map
Frequency
Time
p-value
>0.05
≈0
50
45
40
35
30
25
20
15
10
5
-0.6
-0.5
-0.4
-0.3
-0.2
-0.1

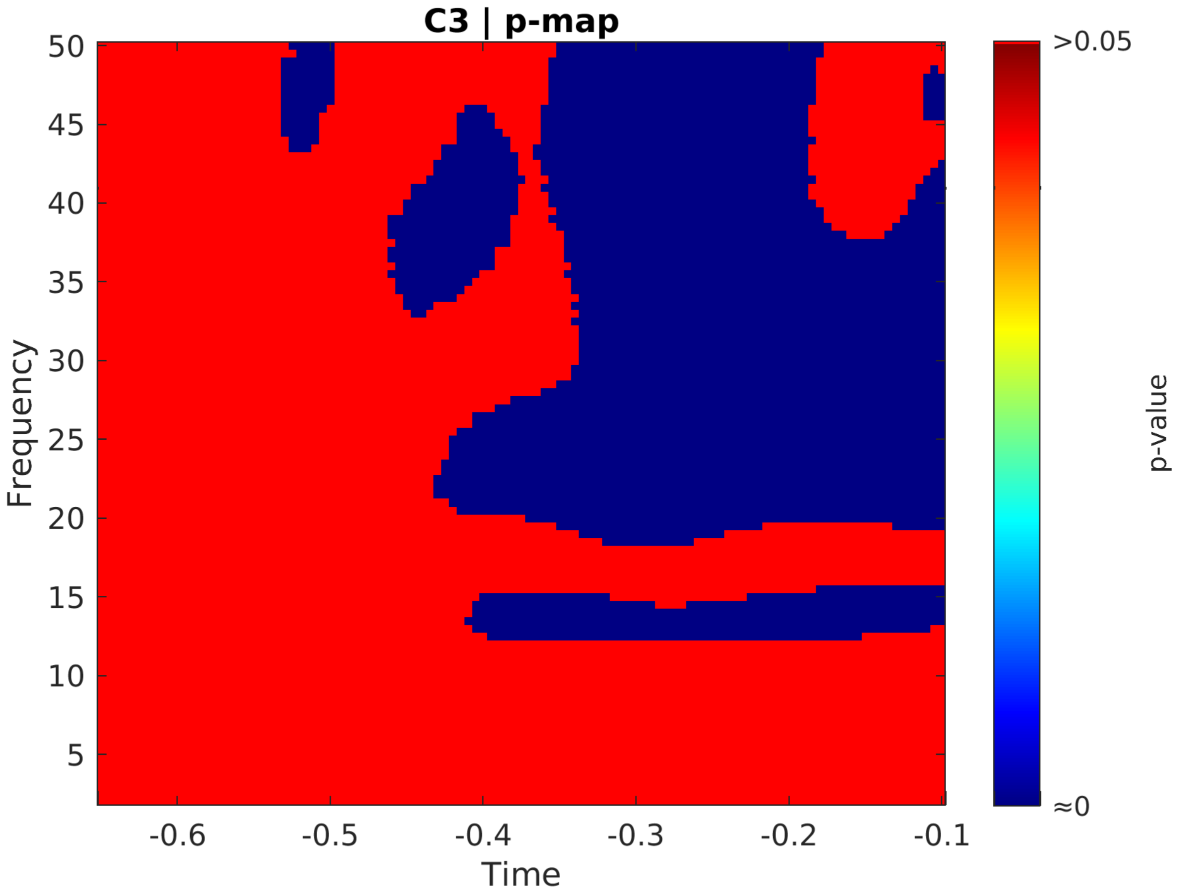

C3 | p-map
50
45
40
35
30
25
20
15
10
5
Frequency
-0.6
-0.5
-0.4
-0.3
-0.2
-0.1
Time
>0.05
≈0
p-value

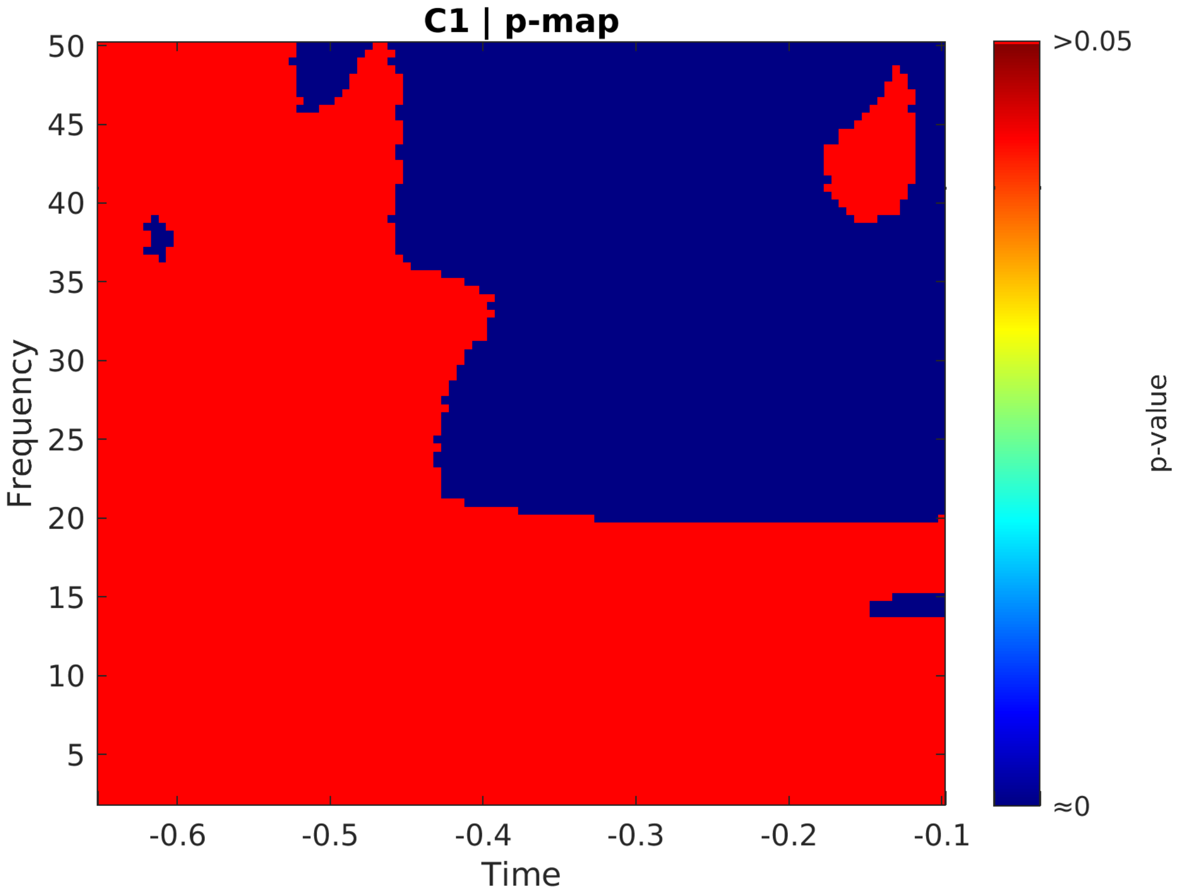
C1 | p-map
Frequency
Time
p-value
>0.05
≈0
50
45
40
35
30
25
20
15
10
5
-0.6
-0.5
-0.4
-0.3
-0.2
-0.1

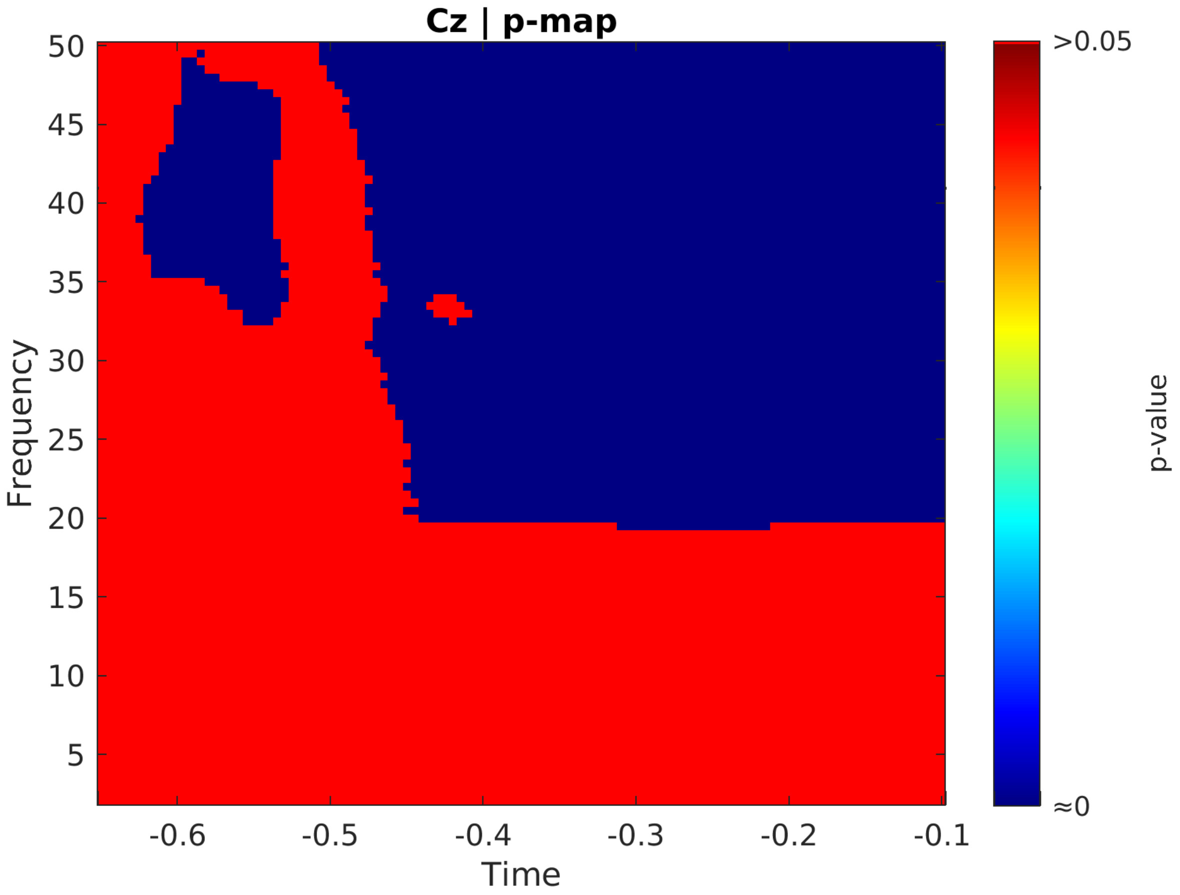
Cz | p-map
Frequency
Time
p-value
>0.05
≈0
50
45
40
35
30
25
20
15
10
5
-0.6
-0.5
-0.4
-0.3
-0.2
-0.1

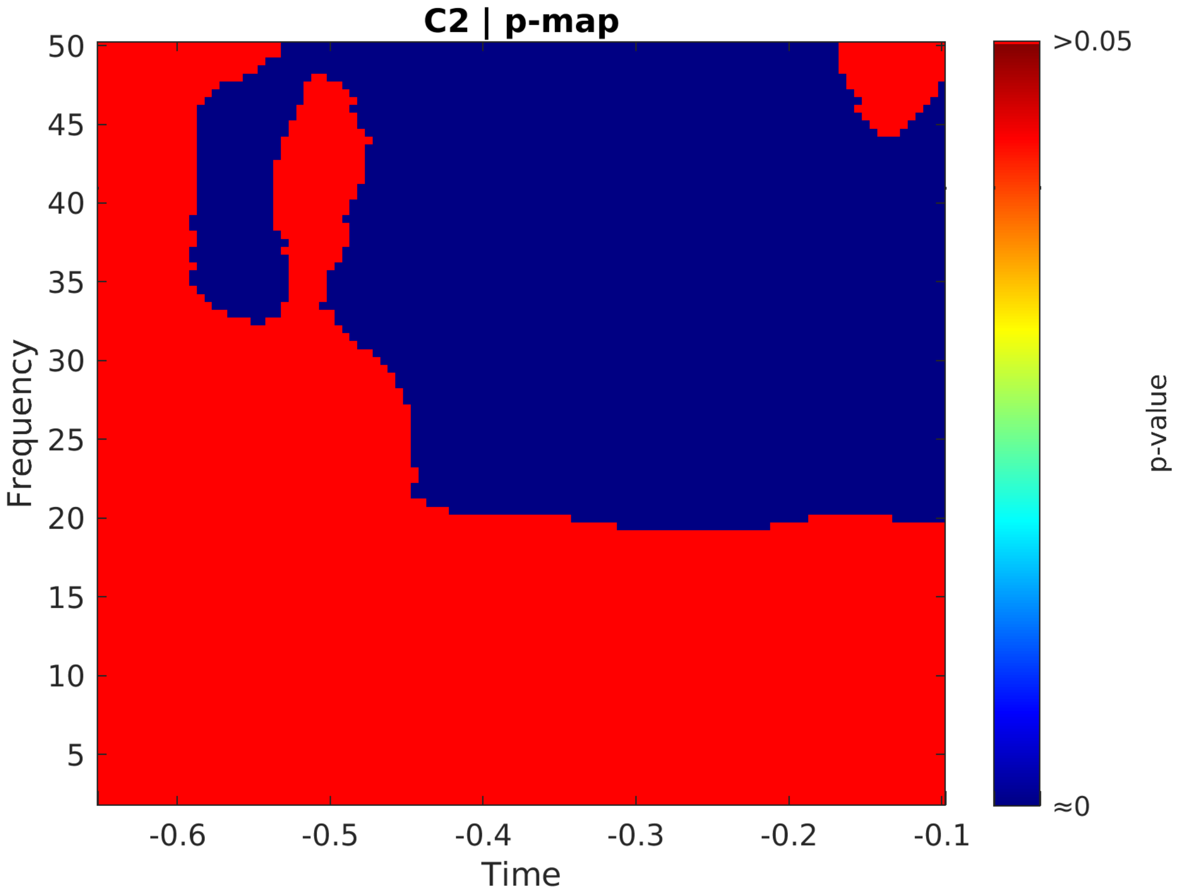
C2 | p-map
Frequency
Time
p-value
>0.05
≈0
50
45
40
35
30
25
20
15
10
5
-0.6
-0.5
-0.4
-0.3
-0.2
-0.1

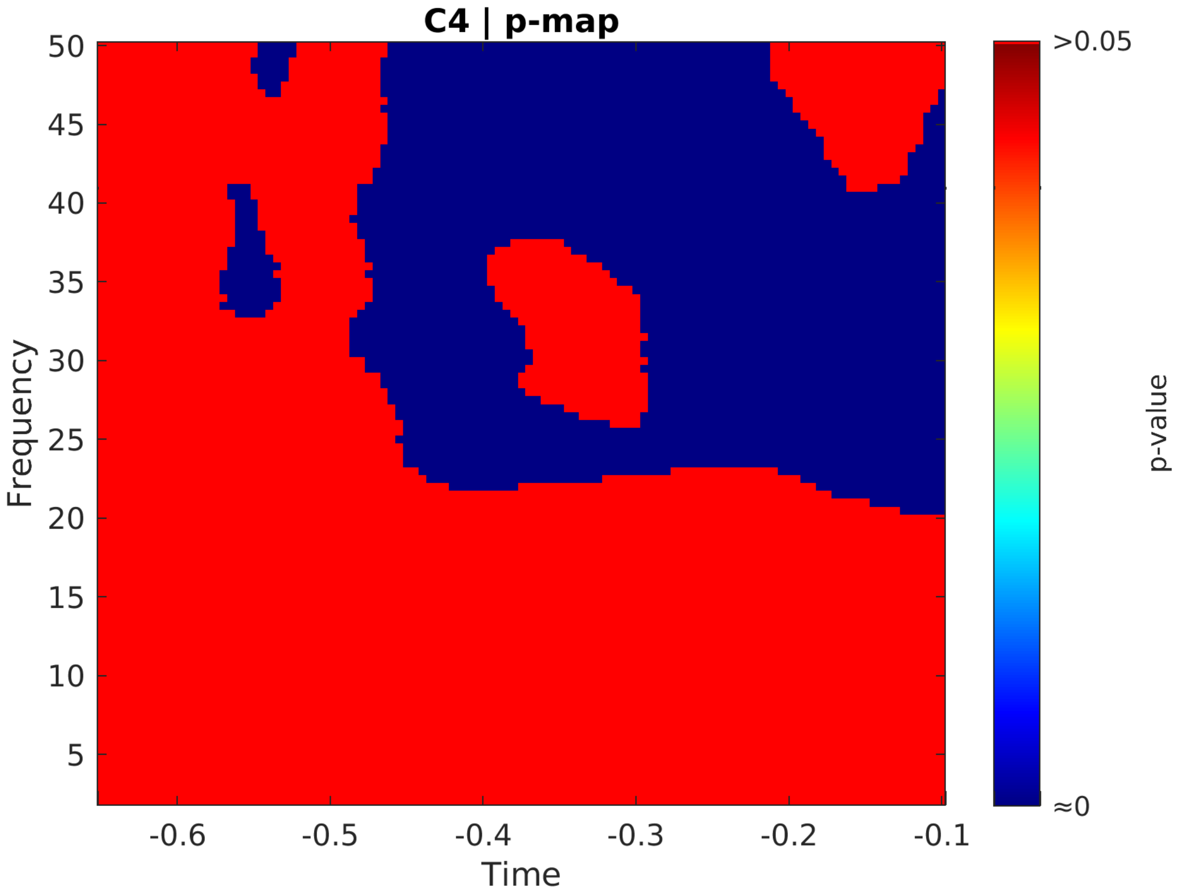

C4 | p-map
Frequency
50
45
40
35
30
25
20
15
10
5
-0.6
-0.5
-0.4
-0.3
-0.2
-0.1
Time
>0.05
≈0
p-value

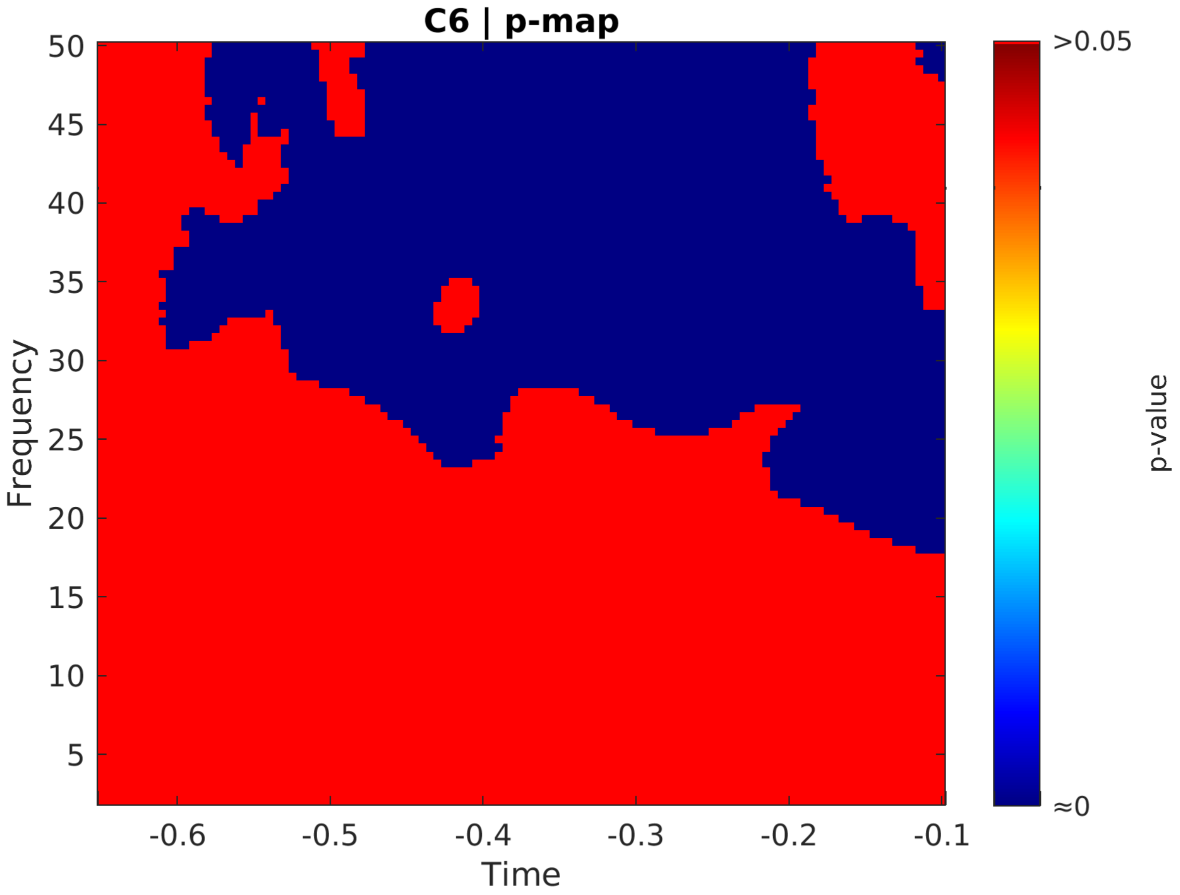
C6 | p-map
Frequency
Time
p-value
>0.05
≈0
50
45
40
35
30
25
20
15
10
5
-0.6
-0.5
-0.4
-0.3
-0.2
-0.1

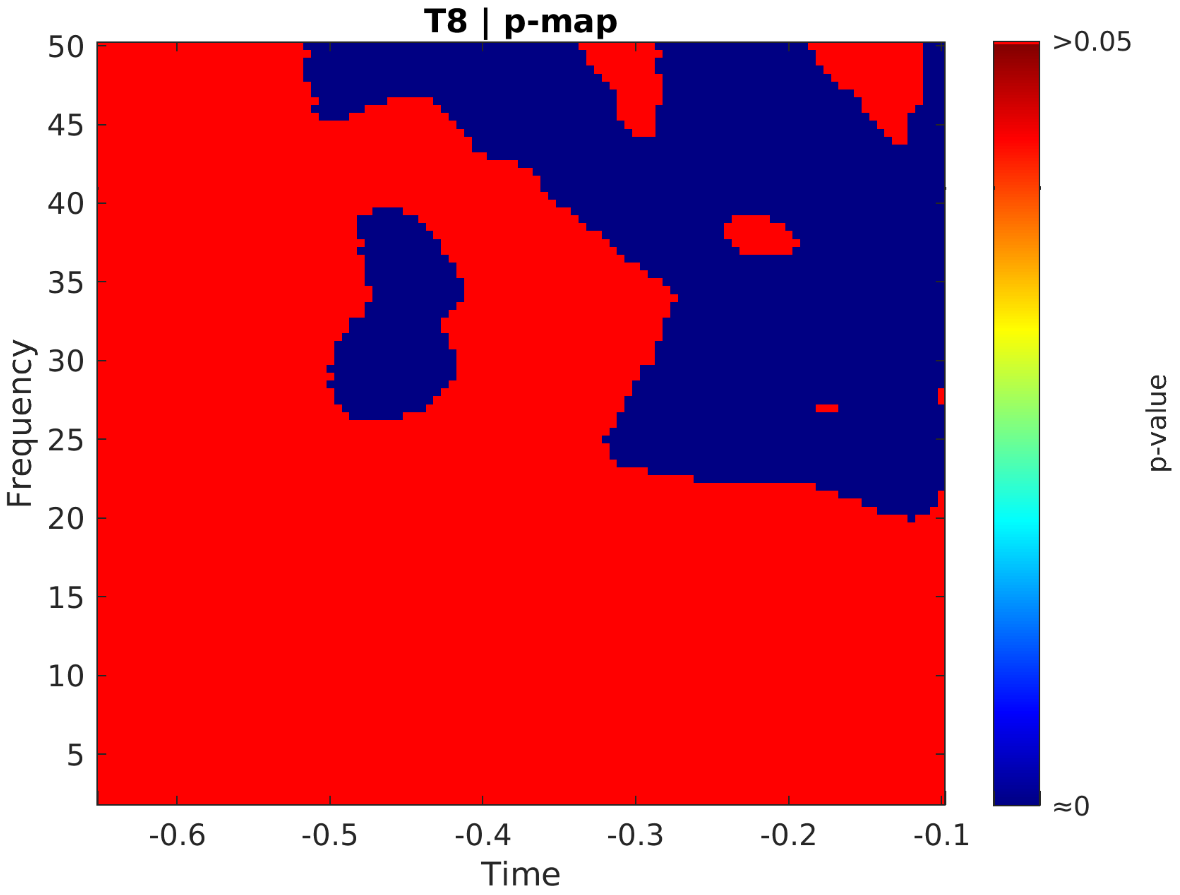

T8 | p-map
50
45
40
35
30
25
20
15
10
5
Frequency
-0.6
-0.5
-0.4
-0.3
-0.2
-0.1
Time
>0.05
≈0
p-value

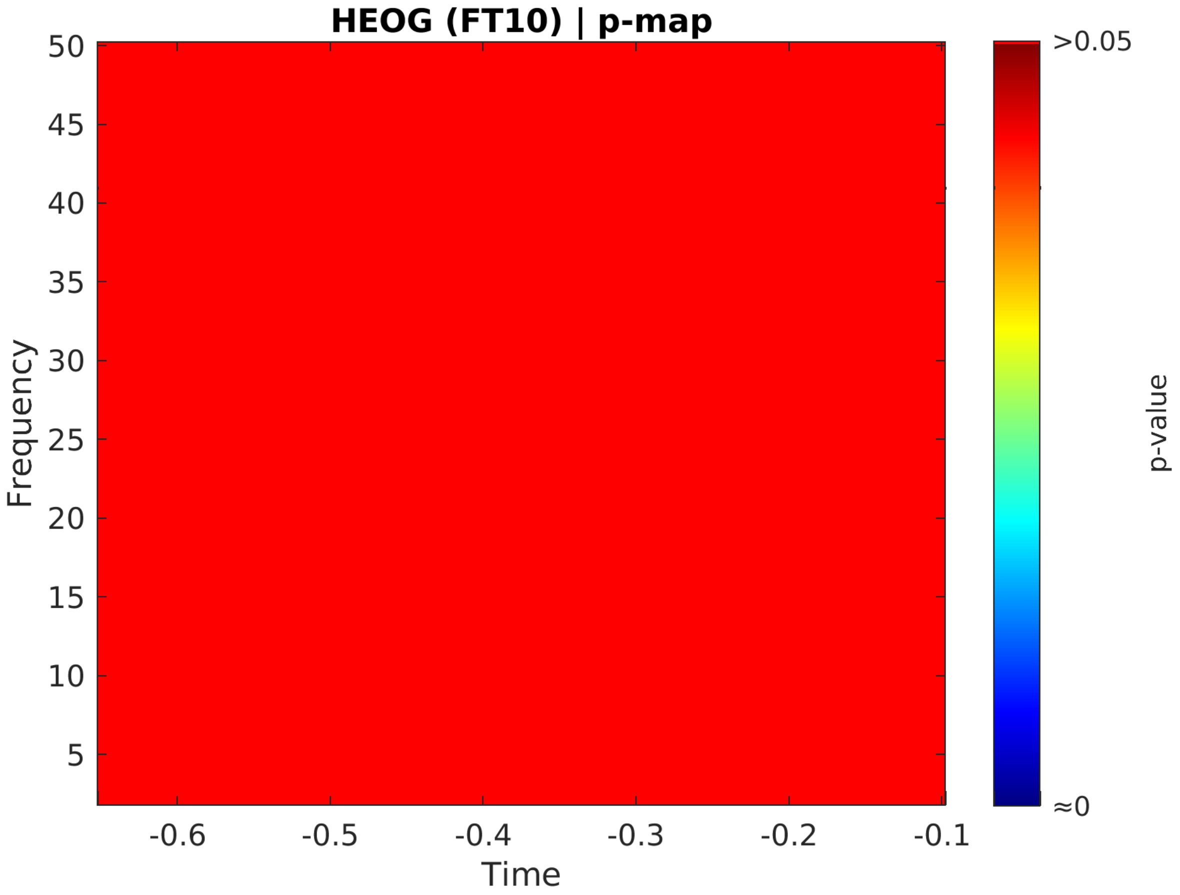

HEOG (FT10) | p-map
Frequency
Time
p-value
>0.05
≈0
50
45
40
35
30
25
20
15
10
5
-0.6
-0.5
-0.4
-0.3
-0.2
-0.1

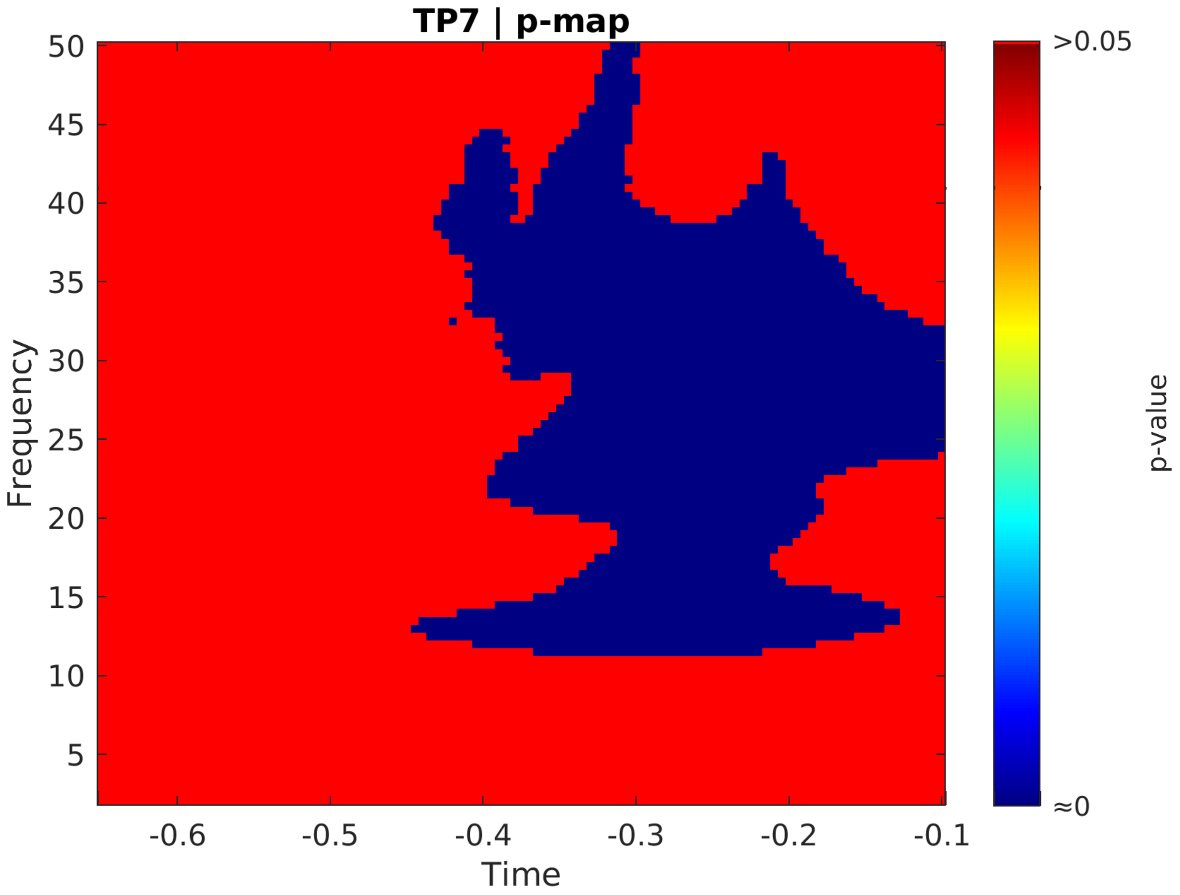
TP7 | p-map
Frequency
Time
50
45
40
35
30
25
20
15
10
5
-0.6
-0.5
-0.4
-0.3
-0.2
-0.1
>0.05
≈0
p-value

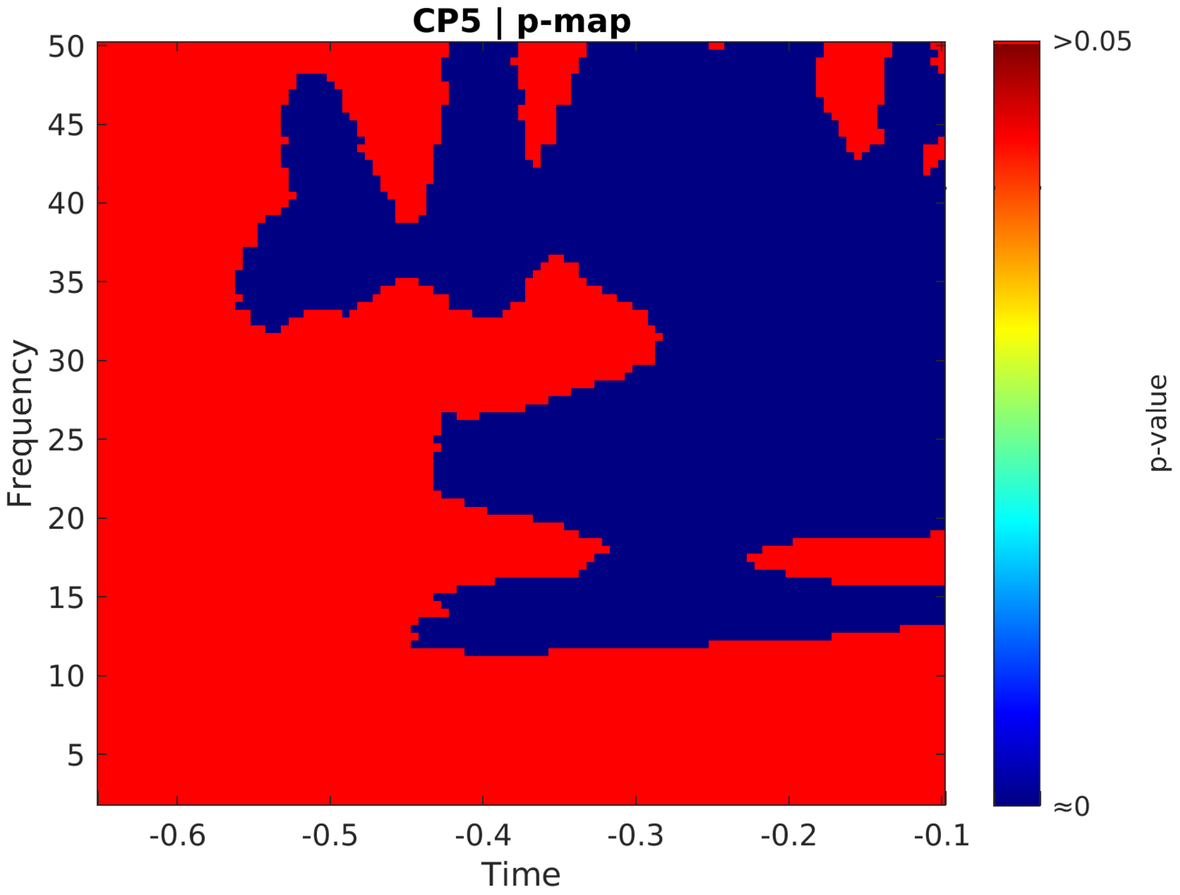
CP5 | p-map
Frequency
Time
p-value
>0.05
≈0
50
45
40
35
30
25
20
15
10
5
-0.6
-0.5
-0.4
-0.3
-0.2
-0.1

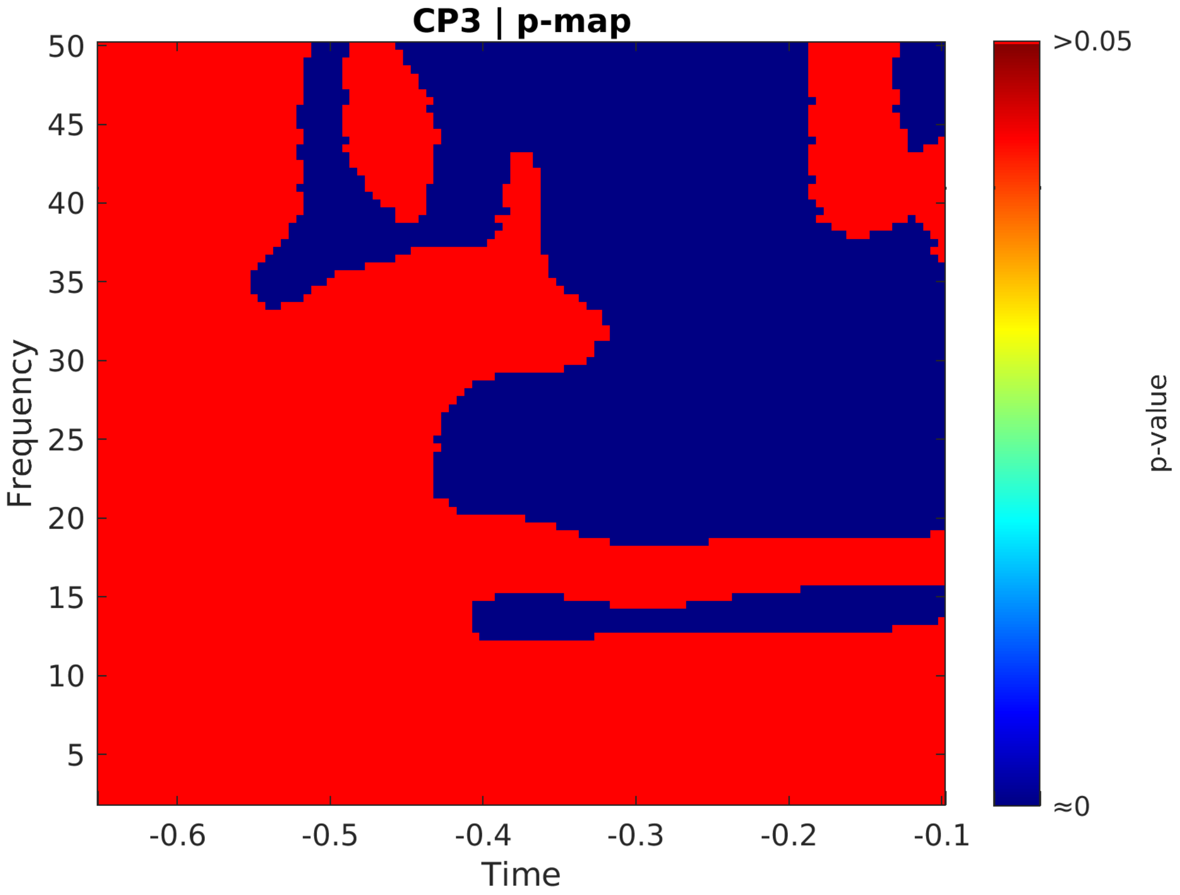
CP3 | p-map
Frequency
50
45
40
35
30
25
20
15
10
5
-0.6
-0.5
-0.4
-0.3
-0.2
-0.1
Time
>0.05
≈0
p-value

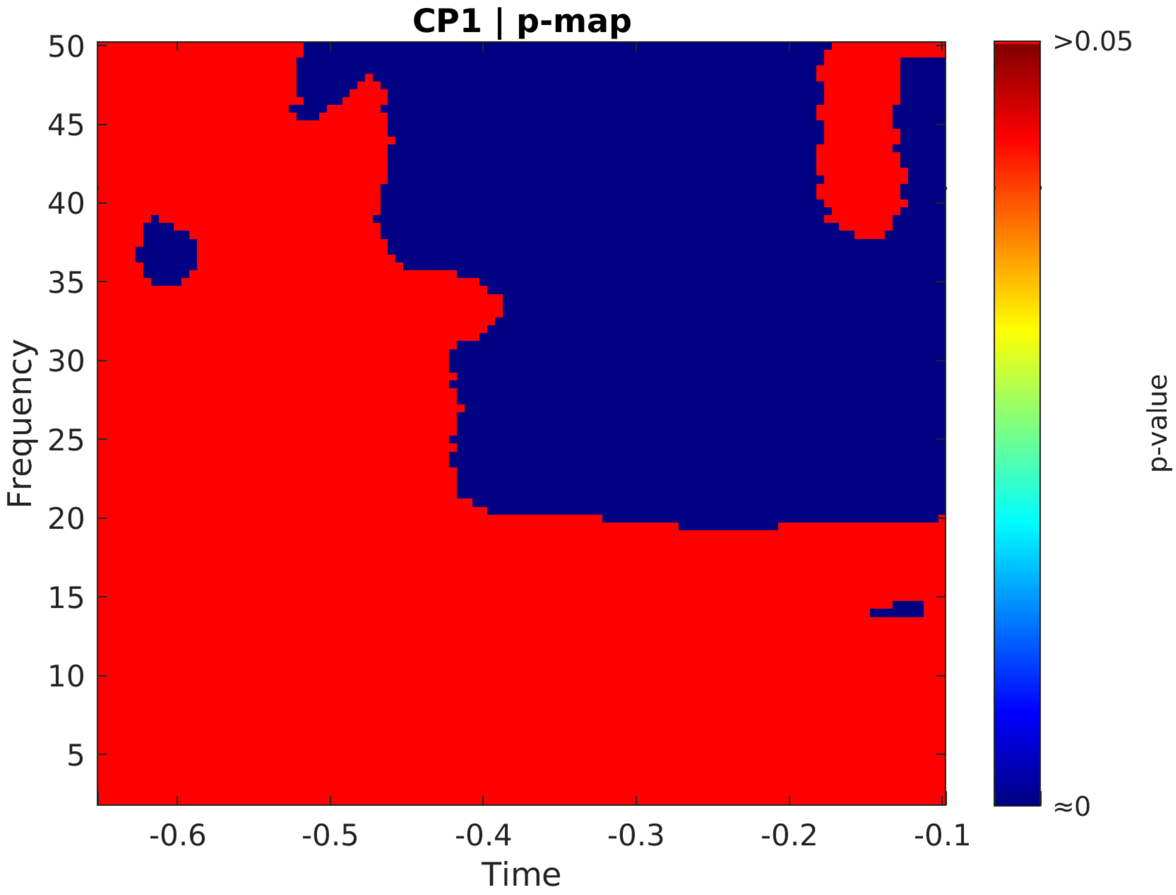

CP1 | p-map
Frequency
Time
p-value
>0.05
≈0
50
45
40
35
30
25
20
15
10
5
-0.6
-0.5
-0.4
-0.3
-0.2
-0.1

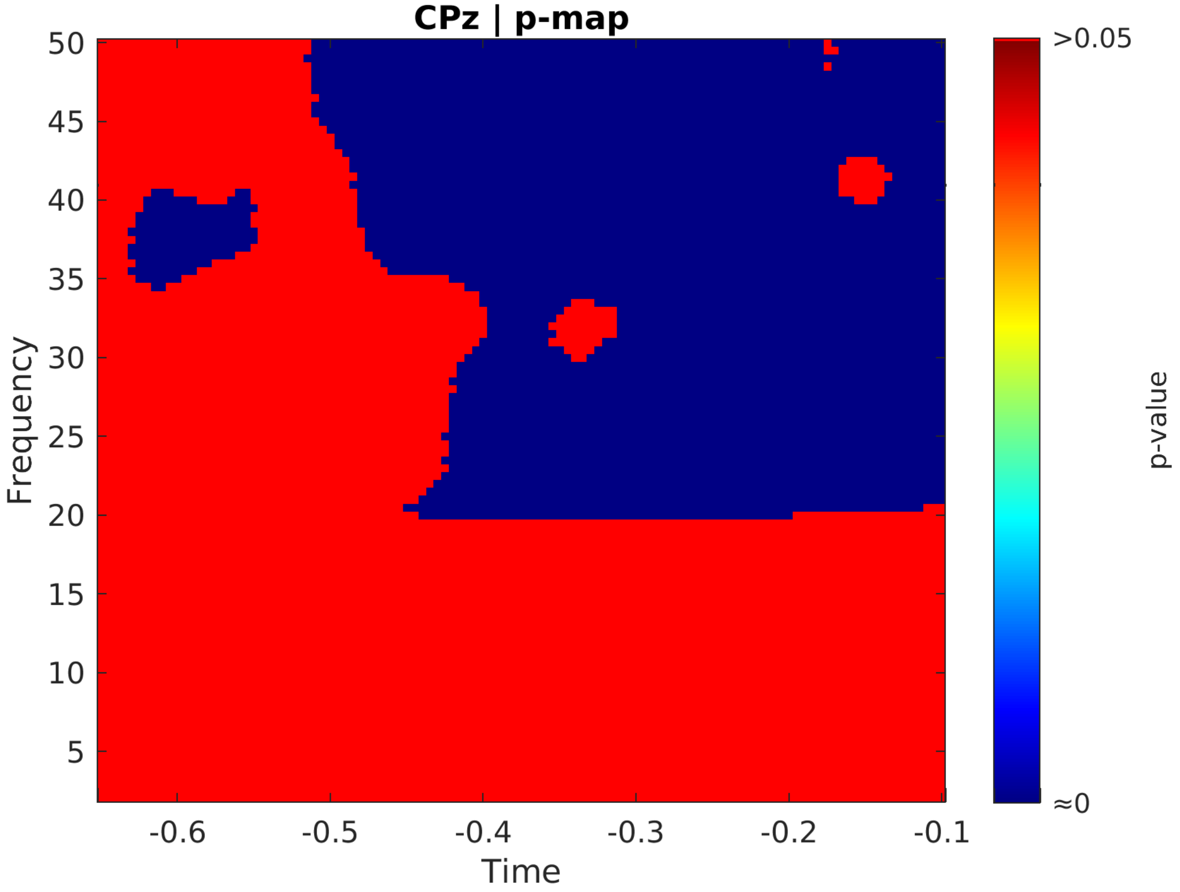
CPz | p-map
Frequency
50
45
40
35
30
25
20
15
10
5
-0.6
-0.5
-0.4
-0.3
-0.2
-0.1
Time
>0.05
≈0
p-value

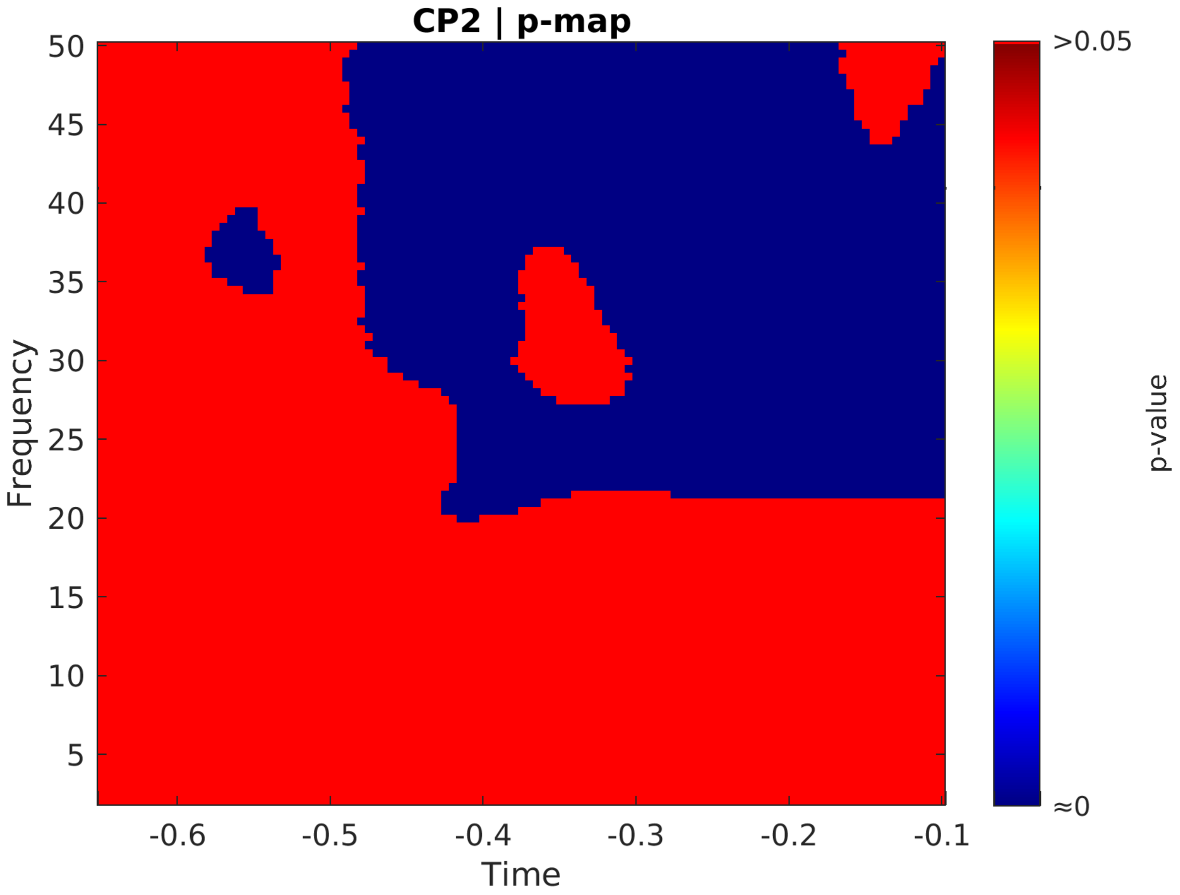

CP2 | p-map
Frequency
Time
p-value
>0.05
≈0
50
45
40
35
30
25
20
15
10
5
-0.6
-0.5
-0.4
-0.3
-0.2
-0.1

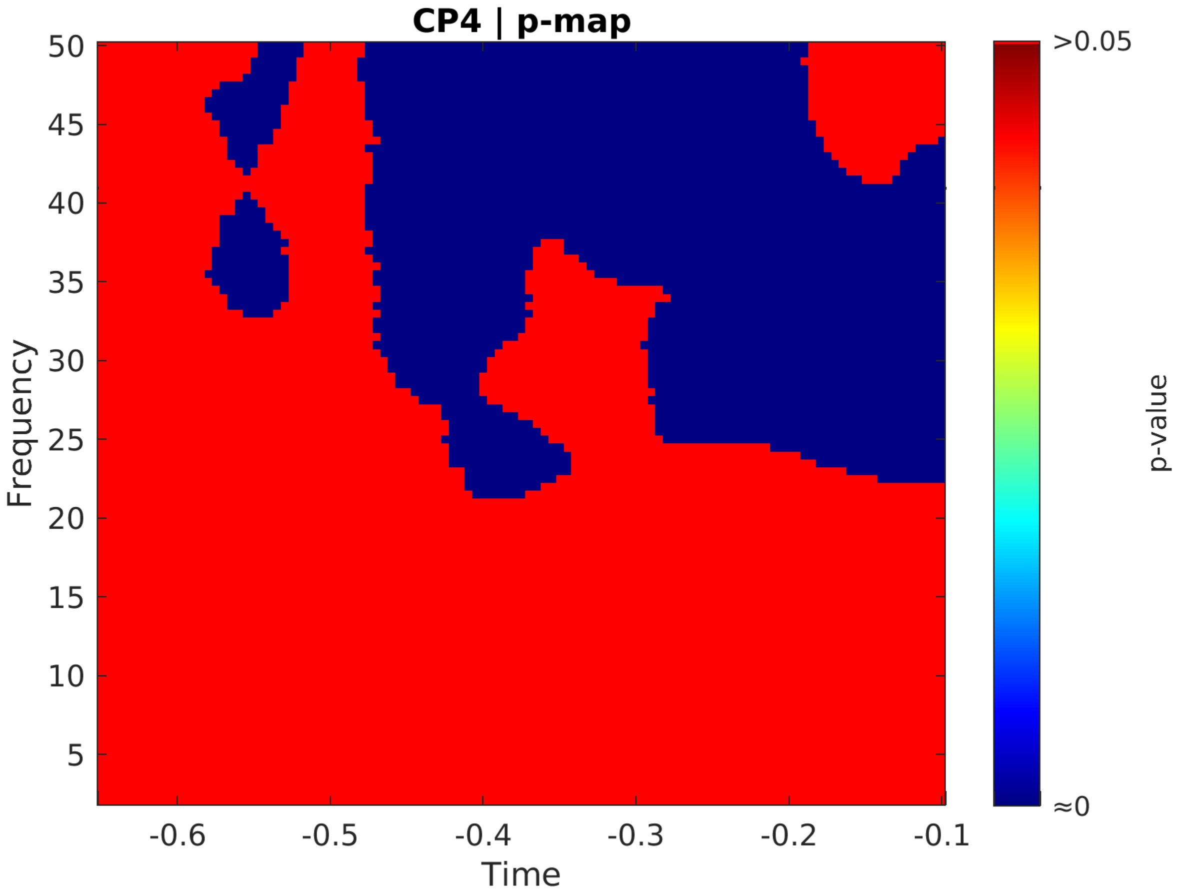

CP4 | p-map
Frequency
Time
p-value
>0.05
≈0
50
45
40
35
30
25
20
15
10
5
-0.6
-0.5
-0.4
-0.3
-0.2
-0.1

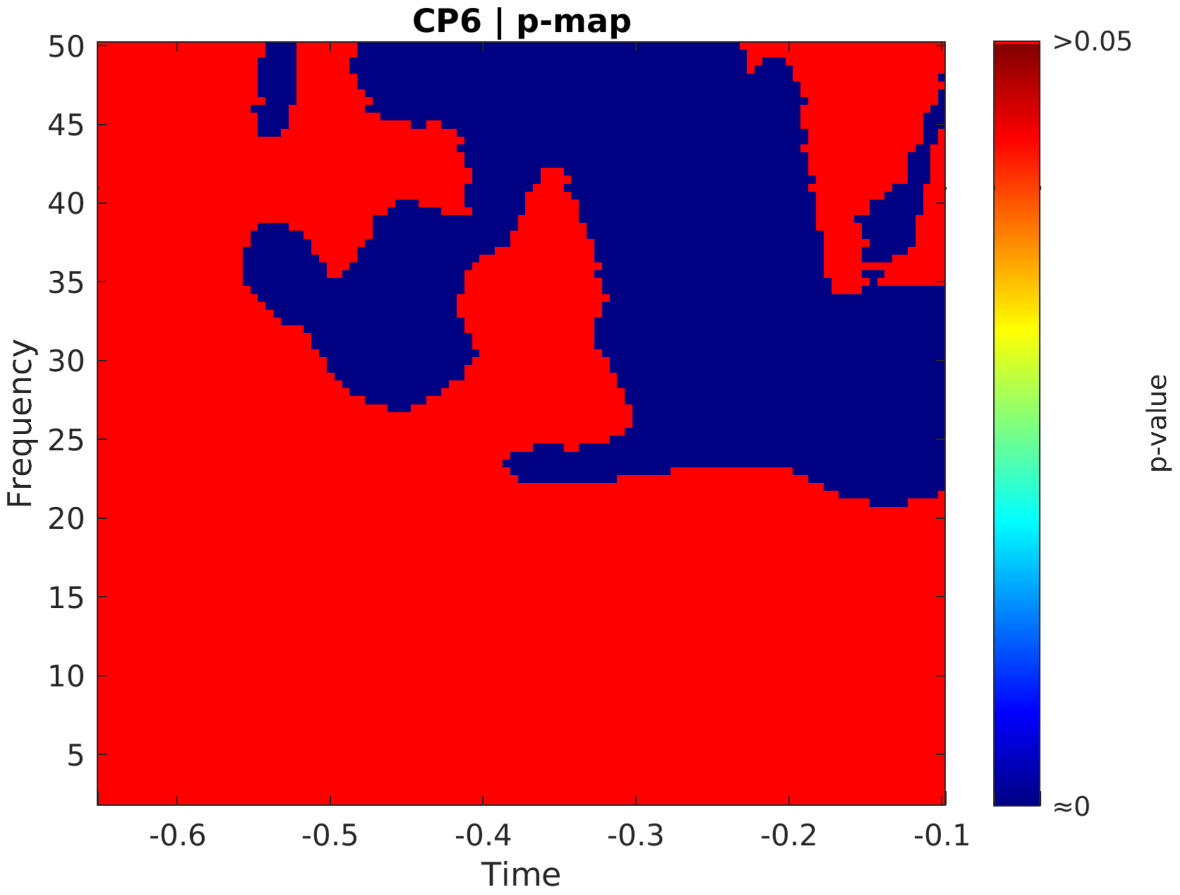

CP6 | p-map
50
45
40
35
30
25
20
15
10
5
Frequency
-0.6
-0.5
-0.4
-0.3
-0.2
-0.1
Time
>0.05
≈0
p-value

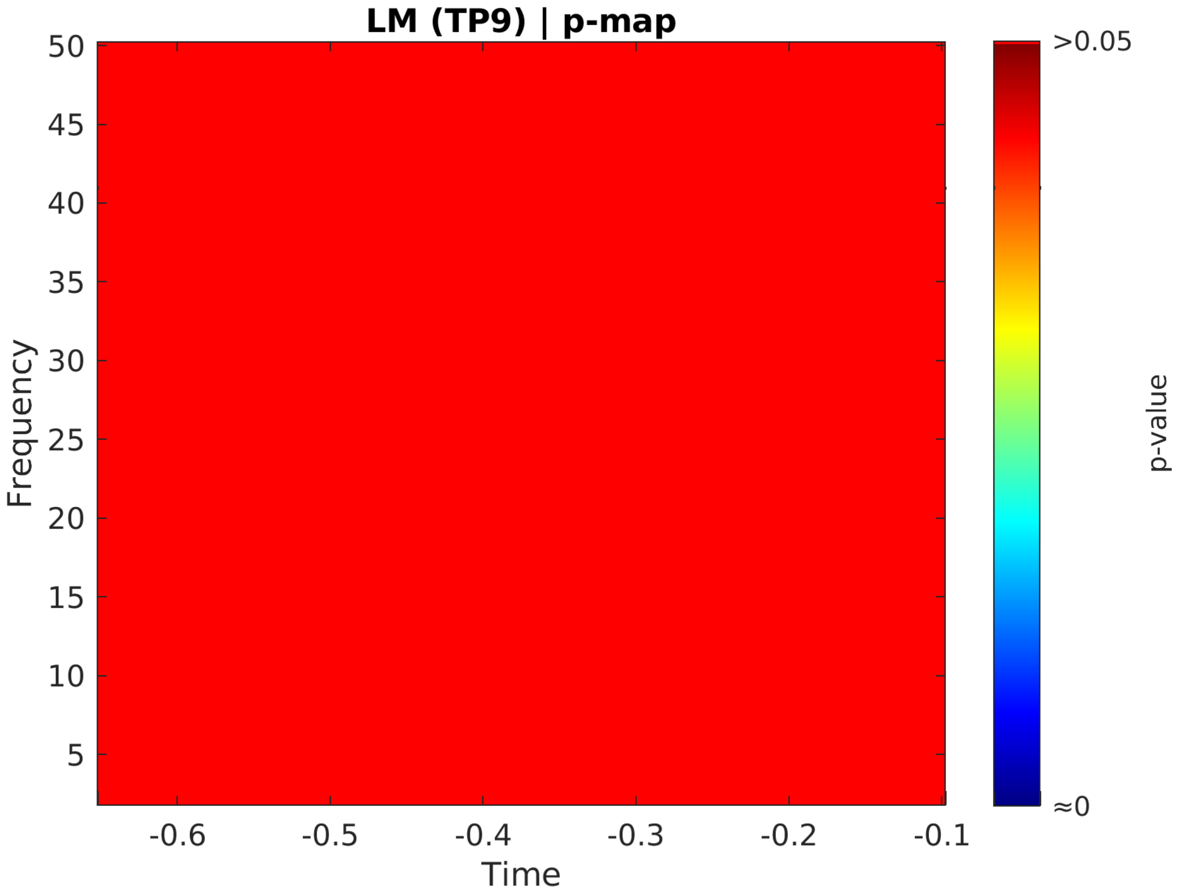
LM (TP9) | p-map
Frequency
50
45
40
35
30
25
20
15
10
5
-0.6
-0.5
-0.4
-0.3
-0.2
-0.1
Time
>0.05
≈0
p-value

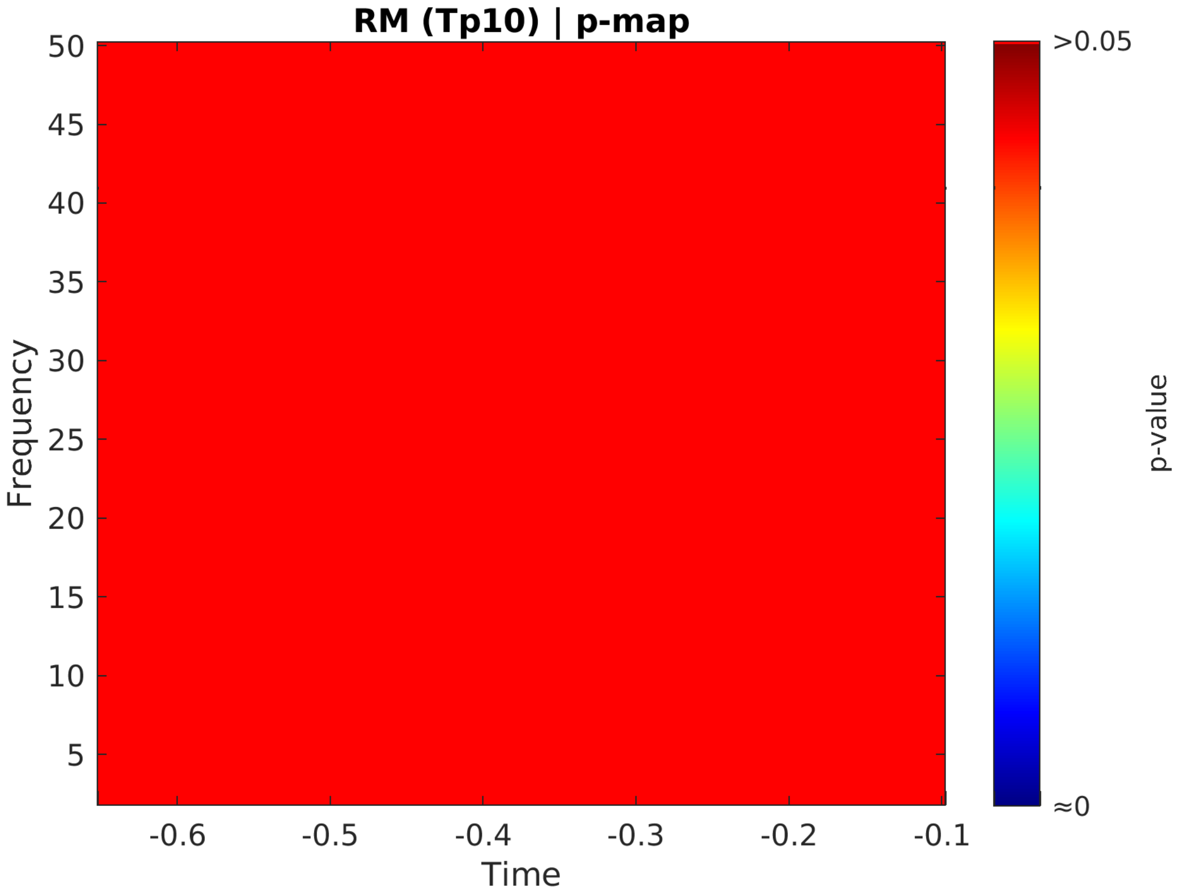
RM (Tp10) | p-map
Frequency
50
45
40
35
30
25
20
15
10
5
-0.6
-0.5
-0.4
-0.3
-0.2
-0.1
Time
>0.05
≈0
p-value

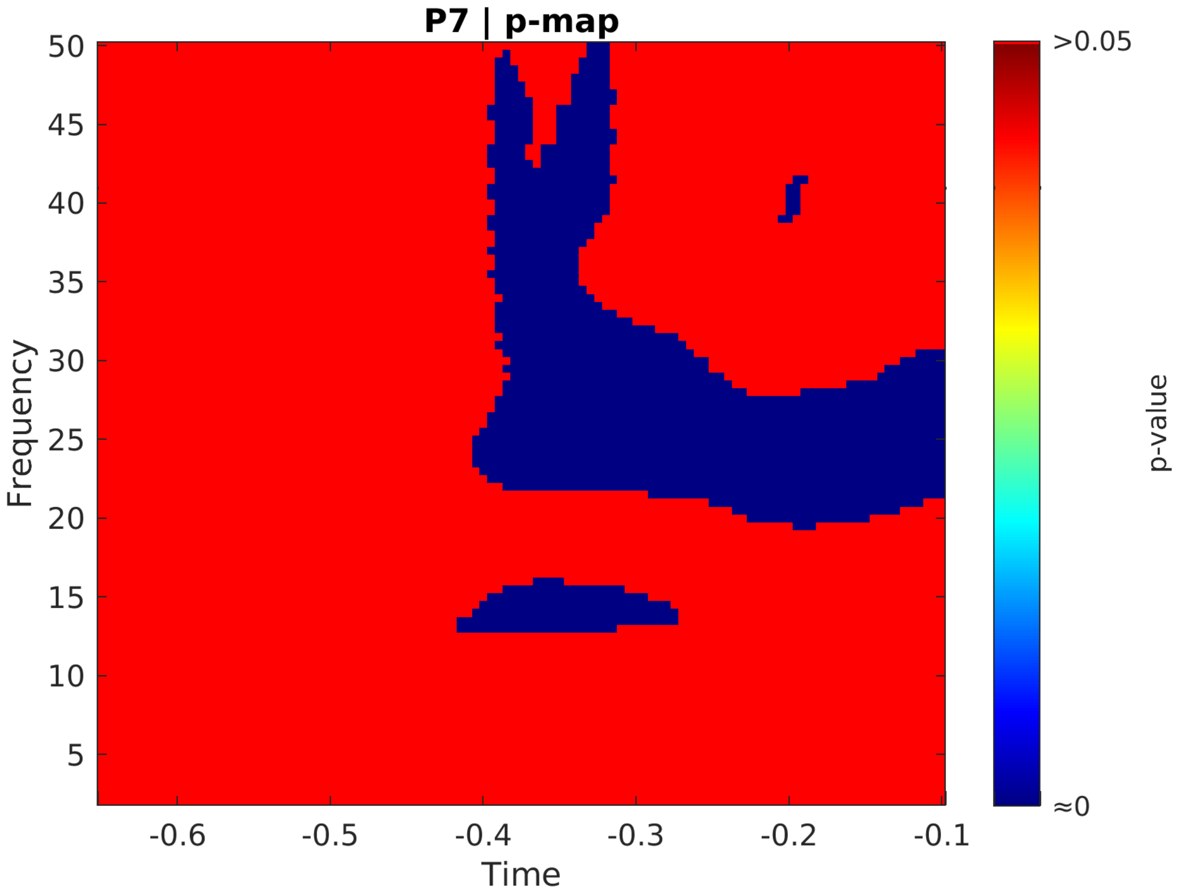
P7 | p-map
Frequency
Time
p-value
>0.05
≈0
50
45
40
35
30
25
20
15
10
5
-0.6
-0.5
-0.4
-0.3
-0.2
-0.1

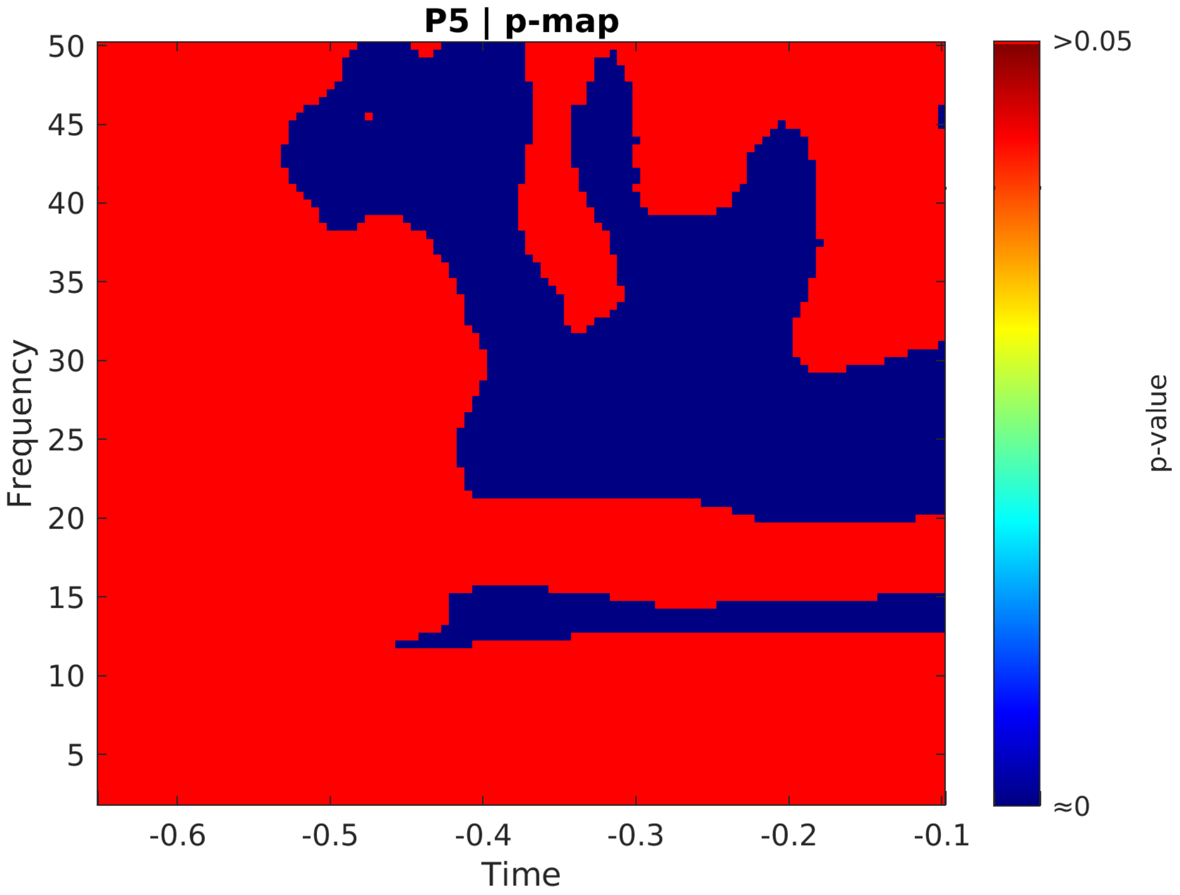
P5 | p-map
Frequency
50
45
40
35
30
25
20
15
10
5
-0.6
-0.5
-0.4
-0.3
-0.2
-0.1
Time
>0.05
≈0
p-value

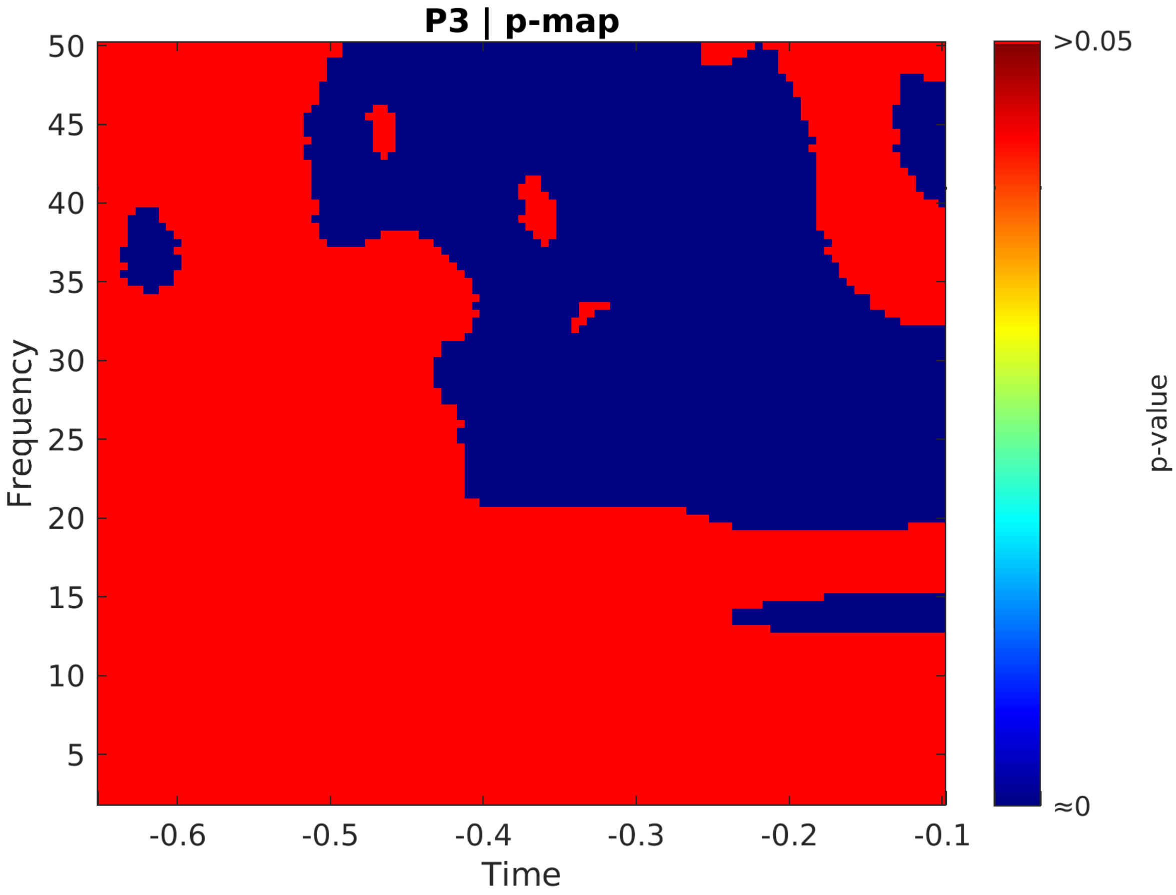
P3 | p-map
50
45
40
35
30
25
20
15
10
5
Frequency
-0.6
-0.5
-0.4
-0.3
-0.2
-0.1
Time
>0.05
≈0
p-value

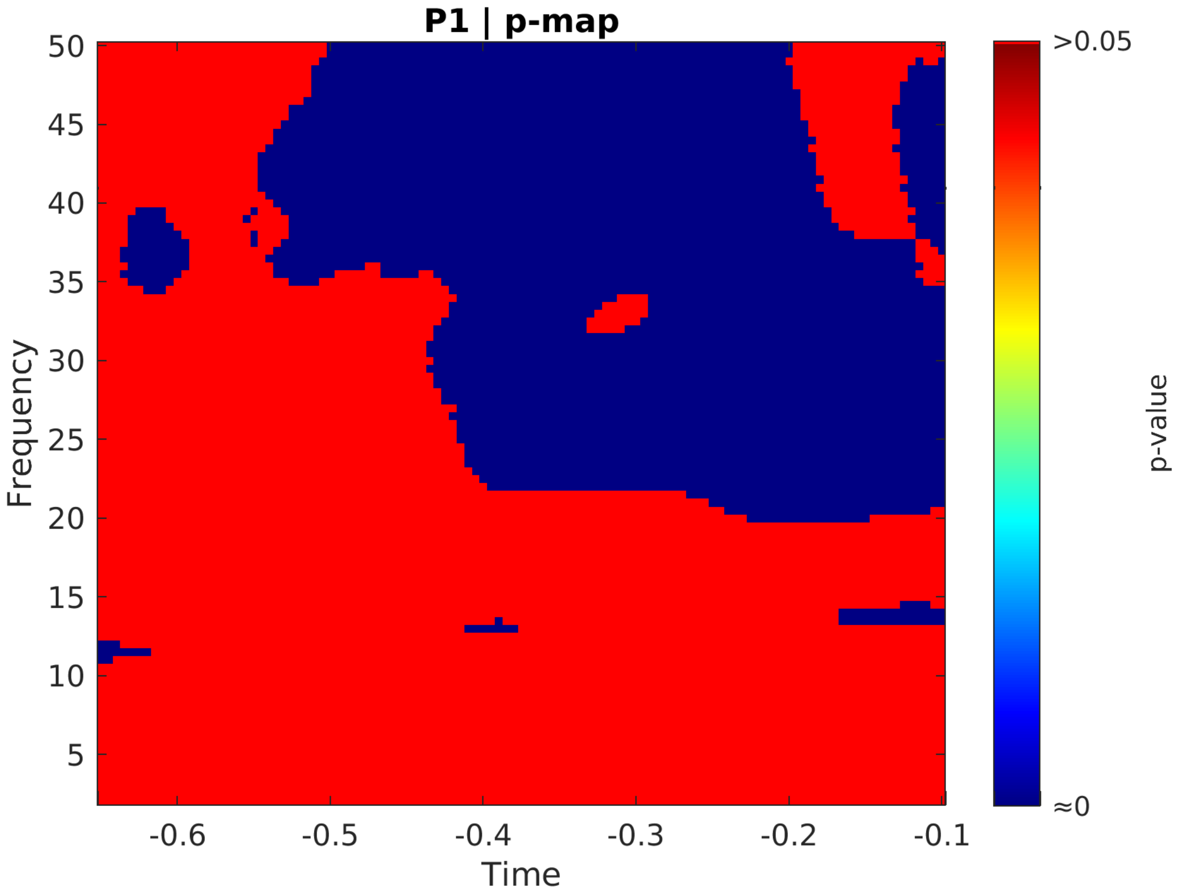
P1 | p-map
Frequency
Time
50
45
40
35
30
25
20
15
10
5
-0.6
-0.5
-0.4
-0.3
-0.2
-0.1
>0.05
≈0
p-value

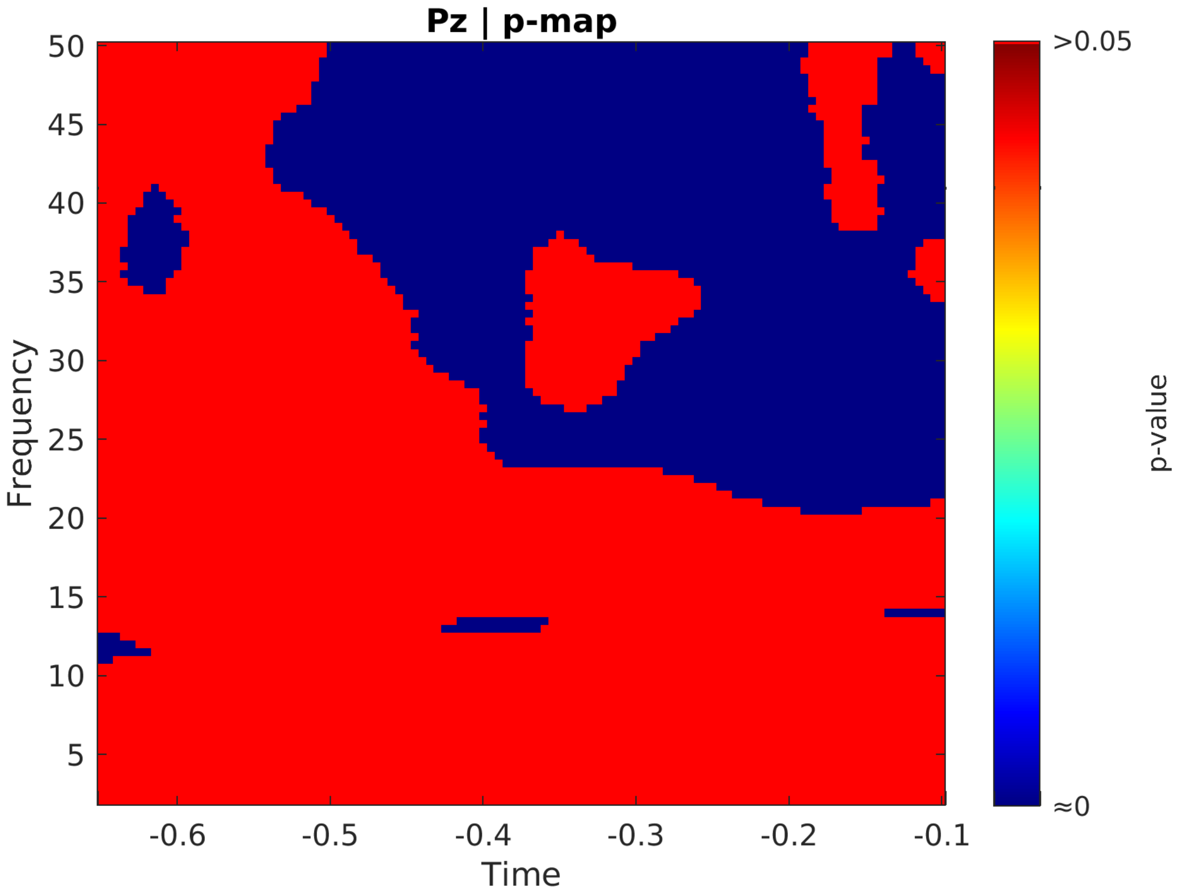
Pz | p-map
Frequency
50
45
40
35
30
25
20
15
10
5
-0.6
-0.5
-0.4
-0.3
-0.2
-0.1
Time
>0.05
≈0
p-value

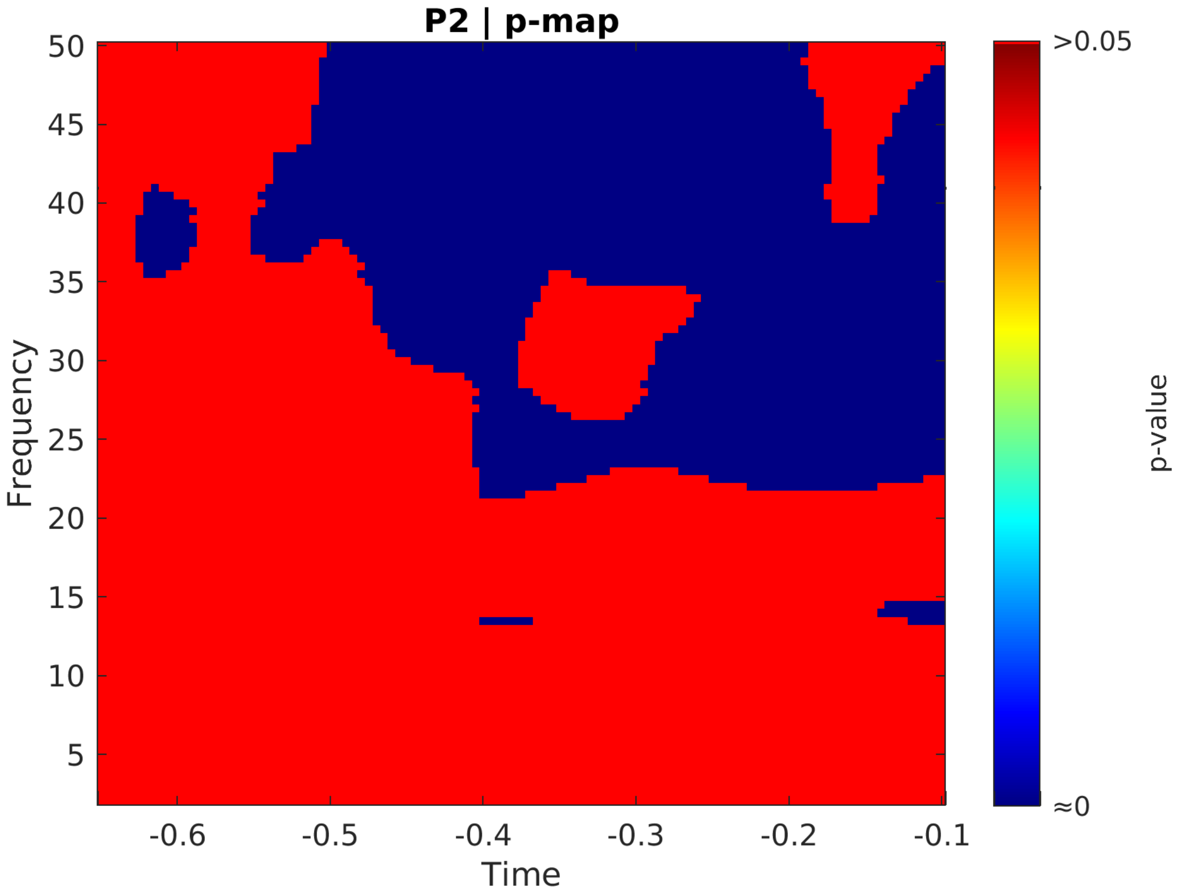
P2 | p-map
Frequency
Time
50
45
40
35
30
25
20
15
10
5
-0.6
-0.5
-0.4
-0.3
-0.2
-0.1
>0.05
≈0
p-value

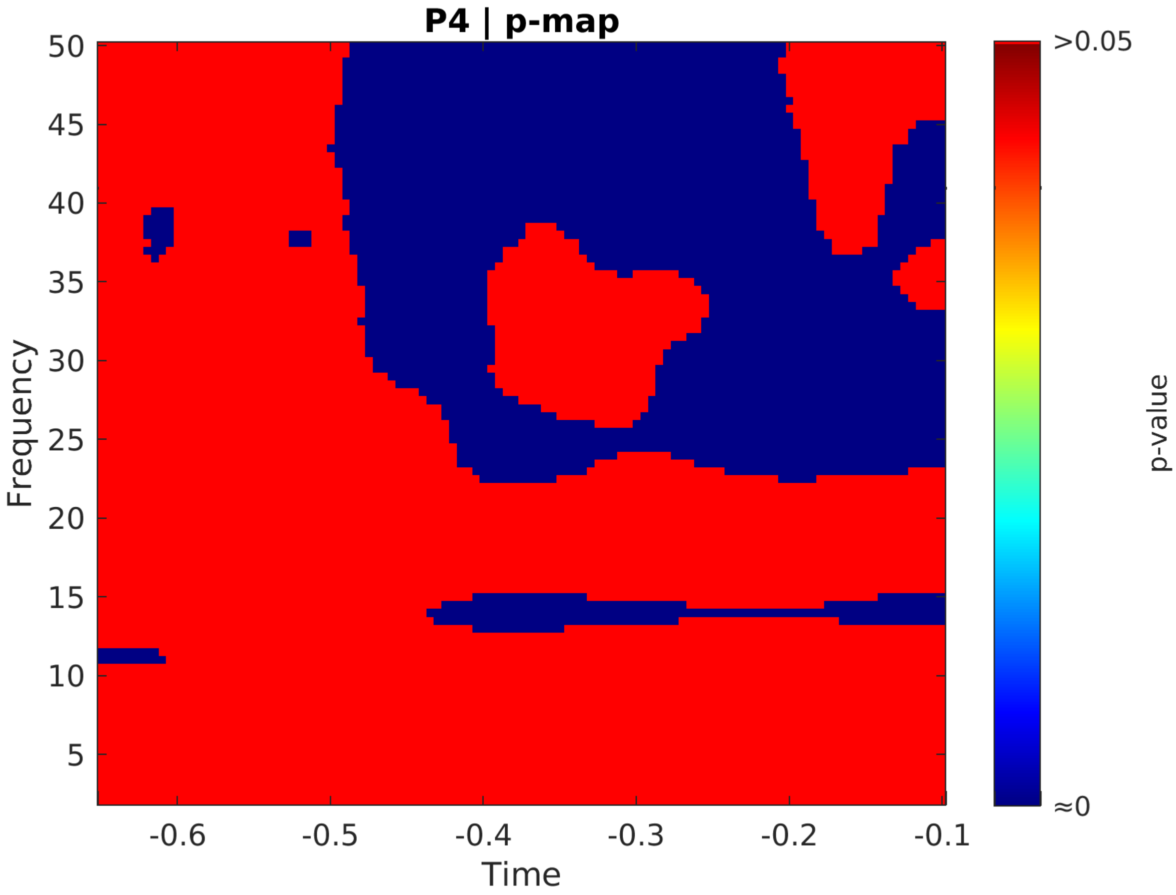
P4 | p-map
Frequency
Time
p-value
>0.05
≈0
50
45
40
35
30
25
20
15
10
5
-0.6
-0.5
-0.4
-0.3
-0.2
-0.1

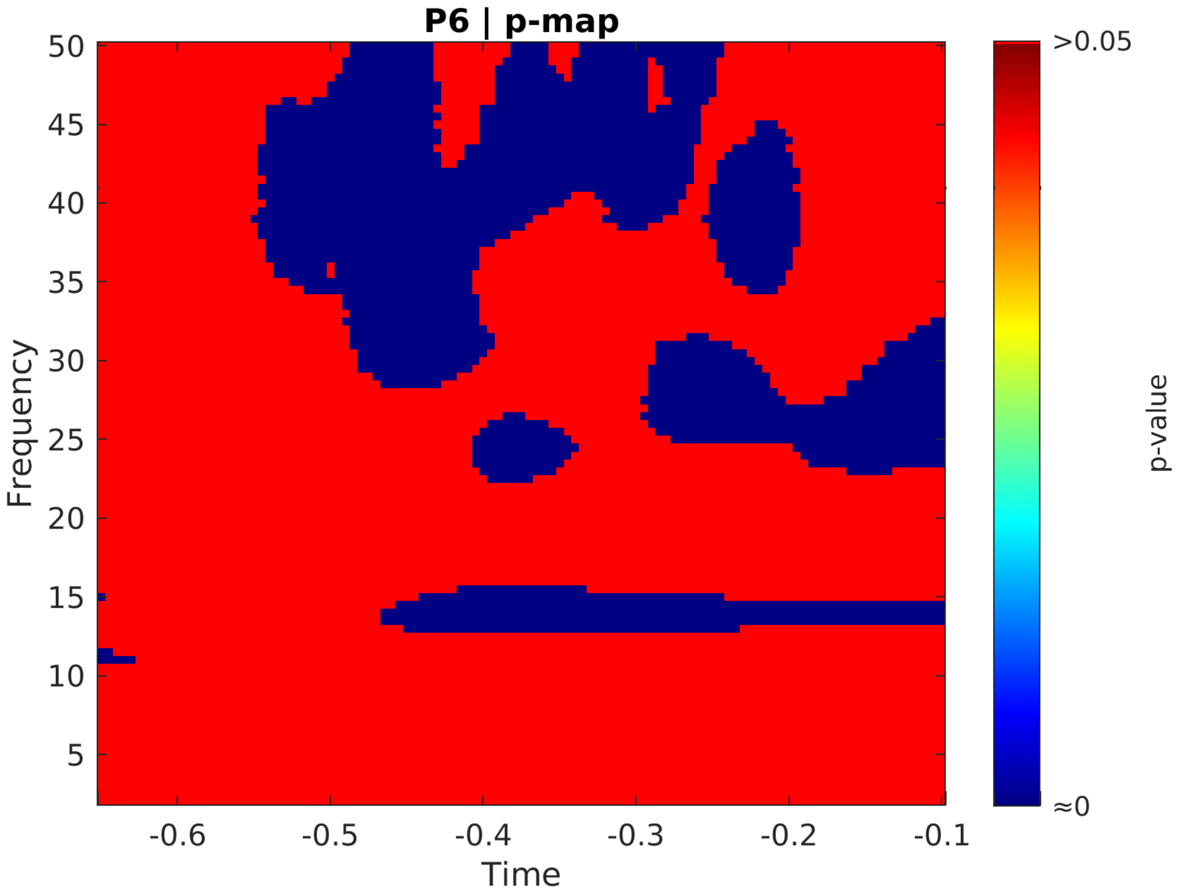
P6 | p-map
Frequency
Time
p-value
>0.05
≈0

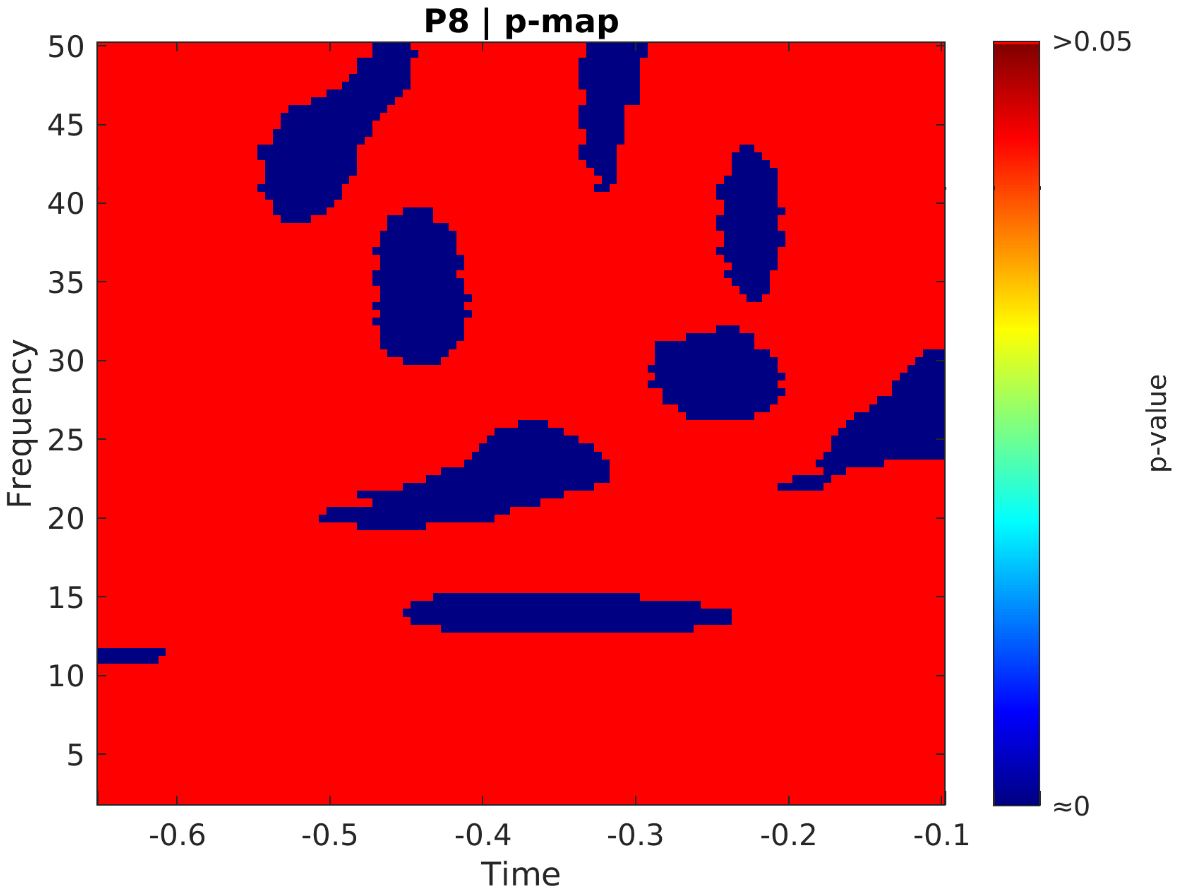

P8 | p-map
Frequency
Time
50
45
40
35
30
25
20
15
10
5
-0.6
-0.5
-0.4
-0.3
-0.2
-0.1
>0.05
≈0
p-value

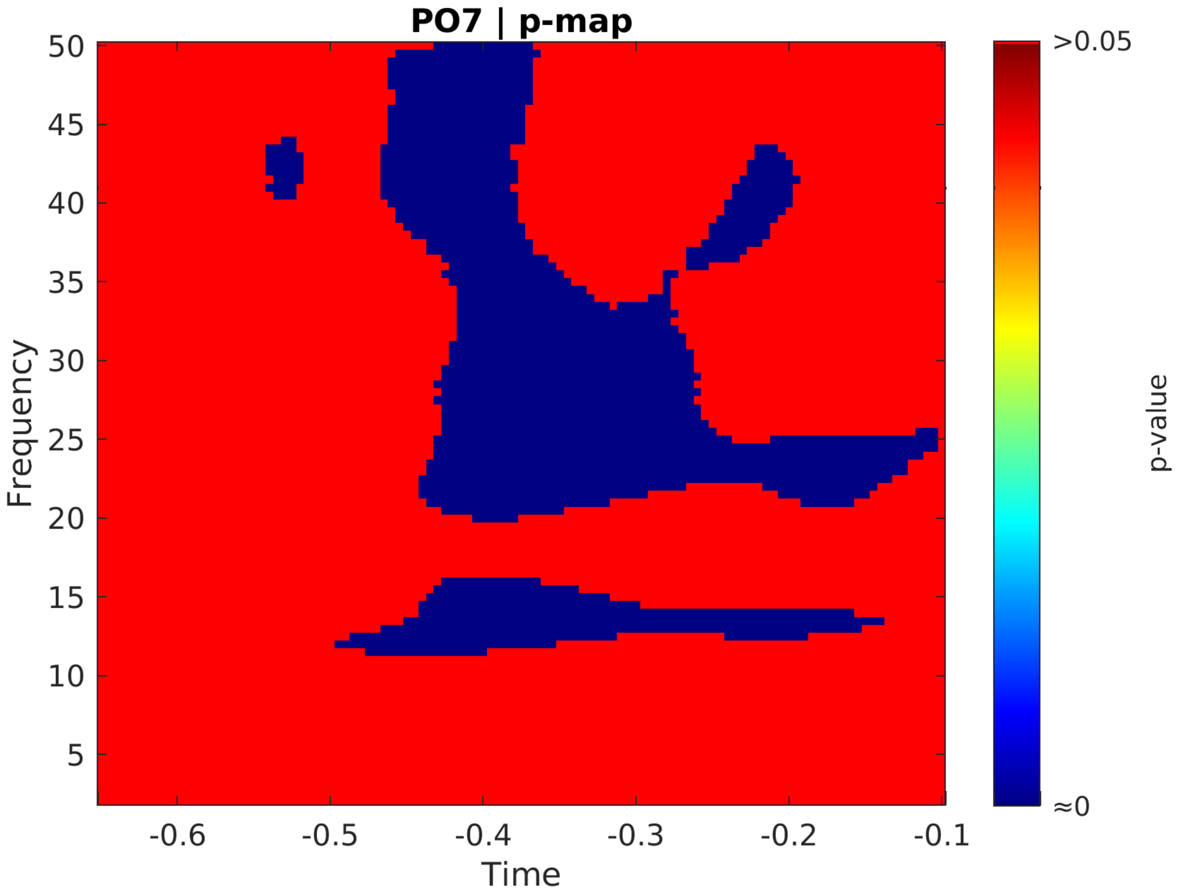
PO7 | p-map
Frequency
Time
50
45
40
35
30
25
20
15
10
5
-0.6
-0.5
-0.4
-0.3
-0.2
-0.1
>0.05
≈0
p-value

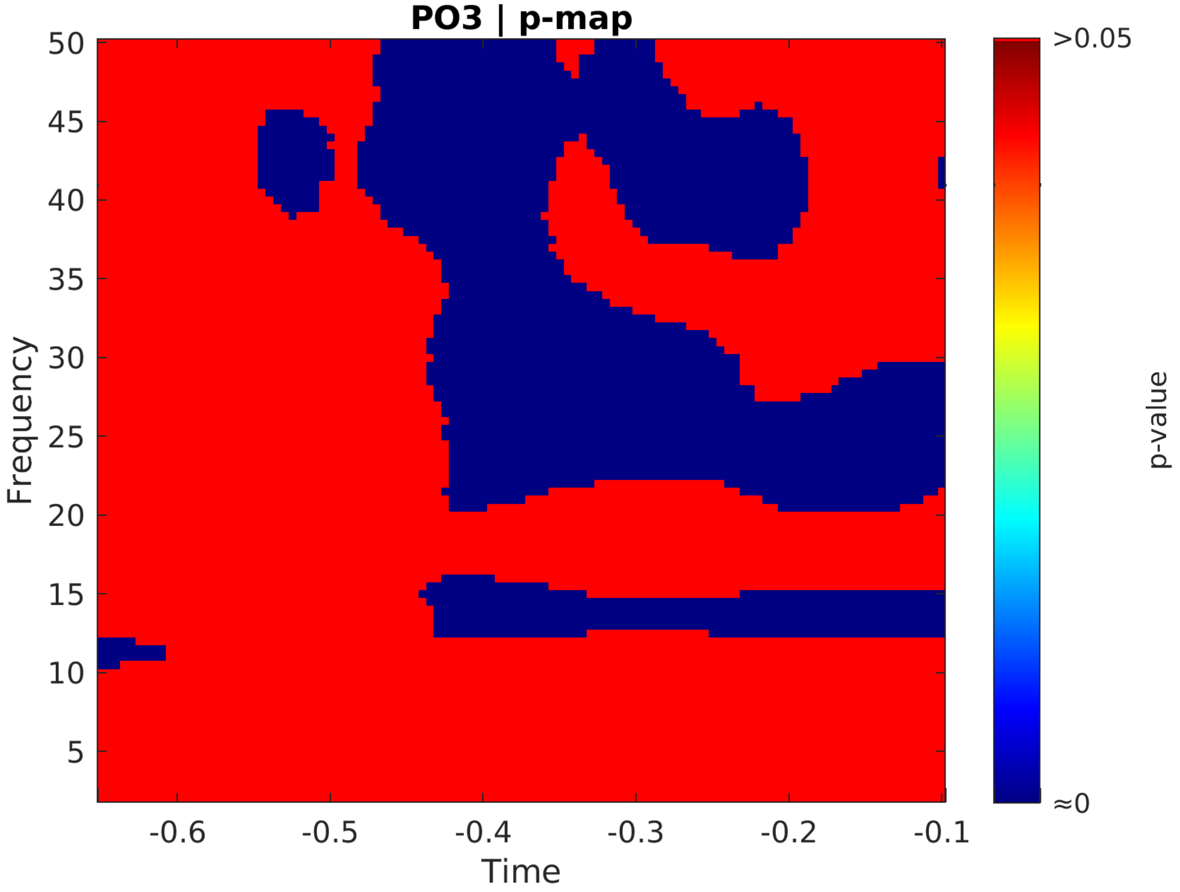
PO3 | p-map
Frequency
Time
p-value
>0.05
≈0
50
45
40
35
30
25
20
15
10
5
-0.6
-0.5
-0.4
-0.3
-0.2
-0.1

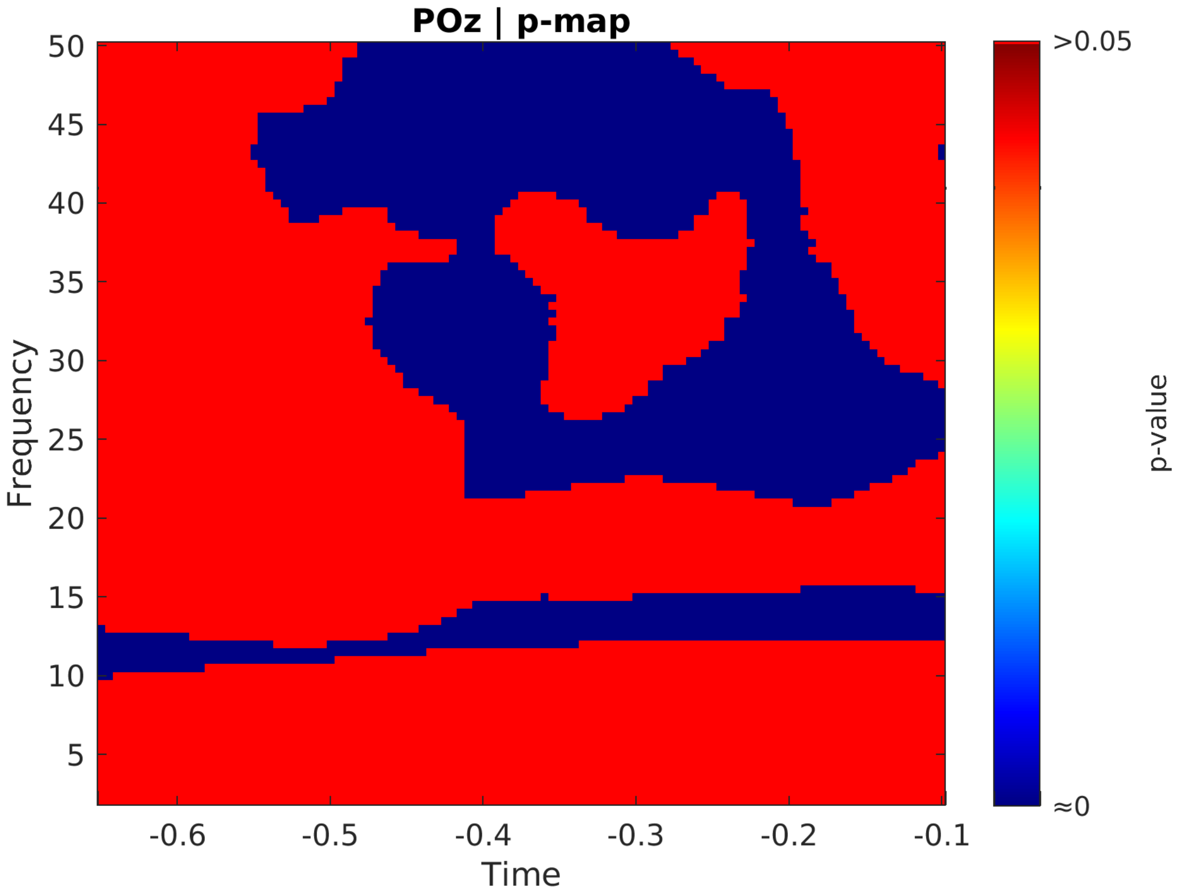
POz | p-map
50
45
40
35
30
25
20
15
10
5
Frequency
-0.6
-0.5
-0.4
-0.3
-0.2
-0.1
Time
>0.05
≈0
p-value

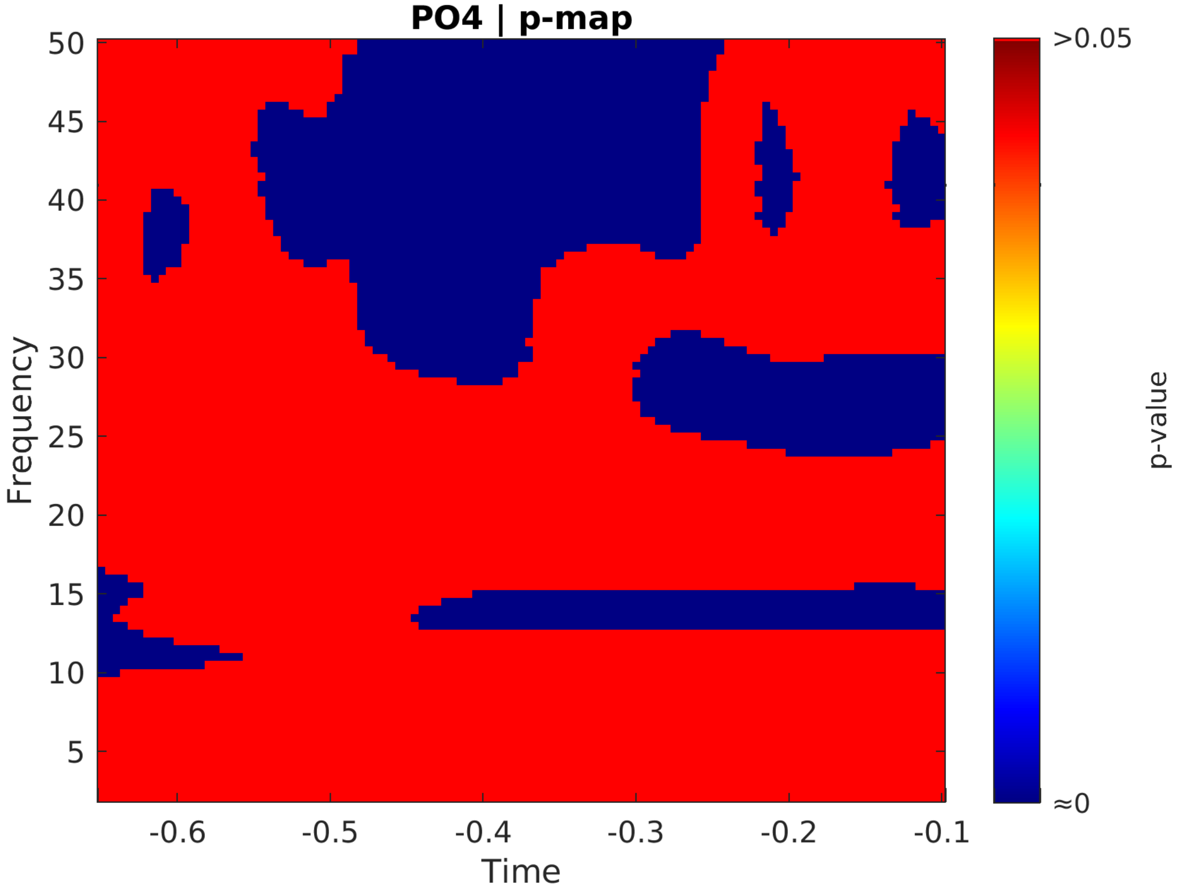
PO4 | p-map
Frequency
Time
p-value
>0.05
≈0
50
45
40
35
30
25
20
15
10
5
-0.6
-0.5
-0.4
-0.3
-0.2
-0.1

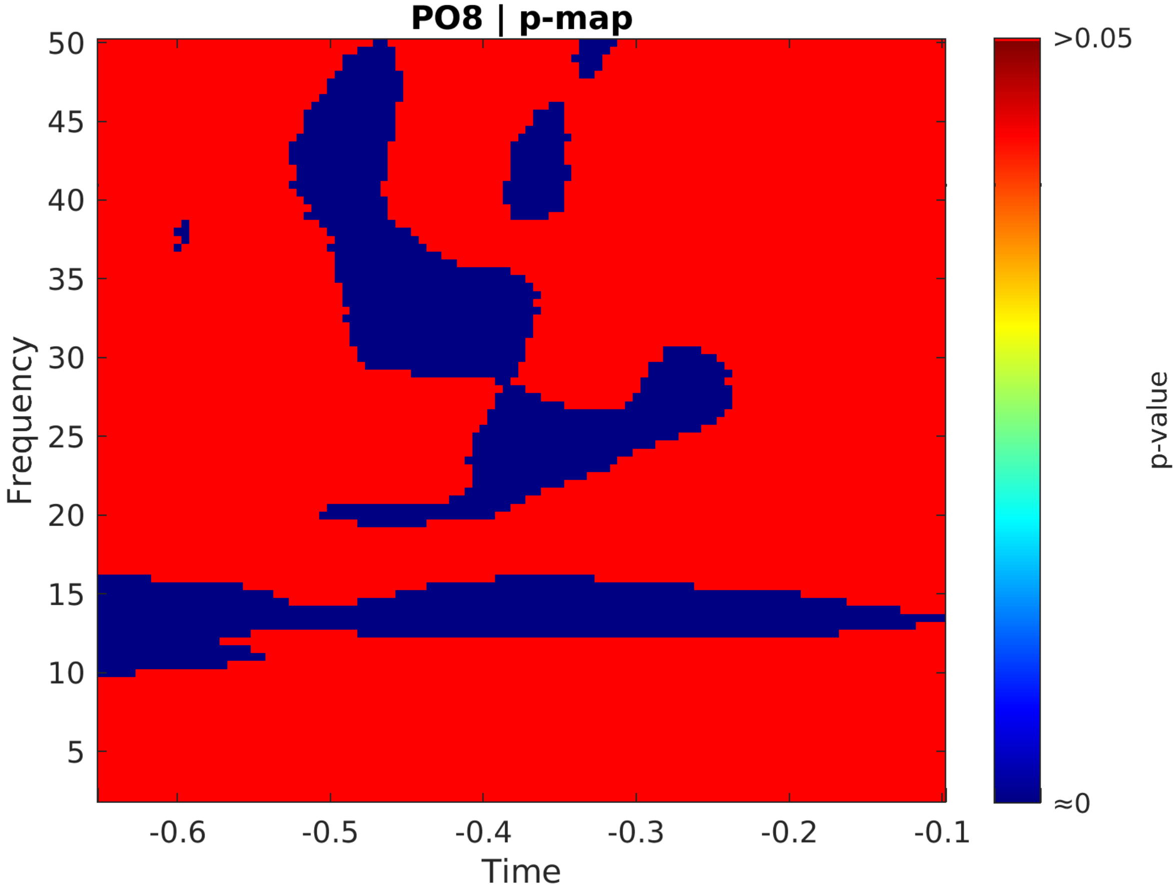

PO8 | p-map
Frequency
50
45
40
35
30
25
20
15
10
5
-0.6
-0.5
-0.4
-0.3
-0.2
-0.1
Time
>0.05
≈0
p-value

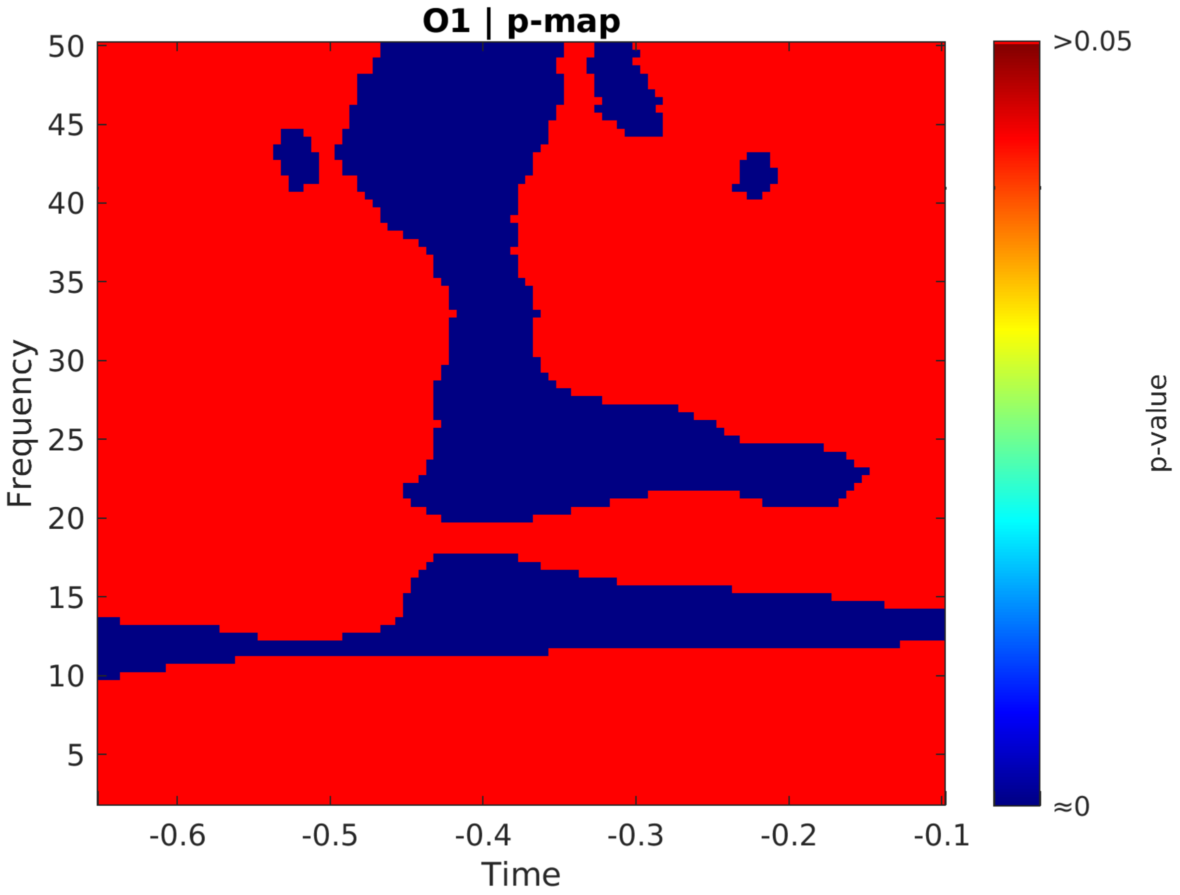

O1 | p-map
Frequency
Time
p-value
>0.05
≈0
50
45
40
35
30
25
20
15
10
5
-0.6
-0.5
-0.4
-0.3
-0.2
-0.1

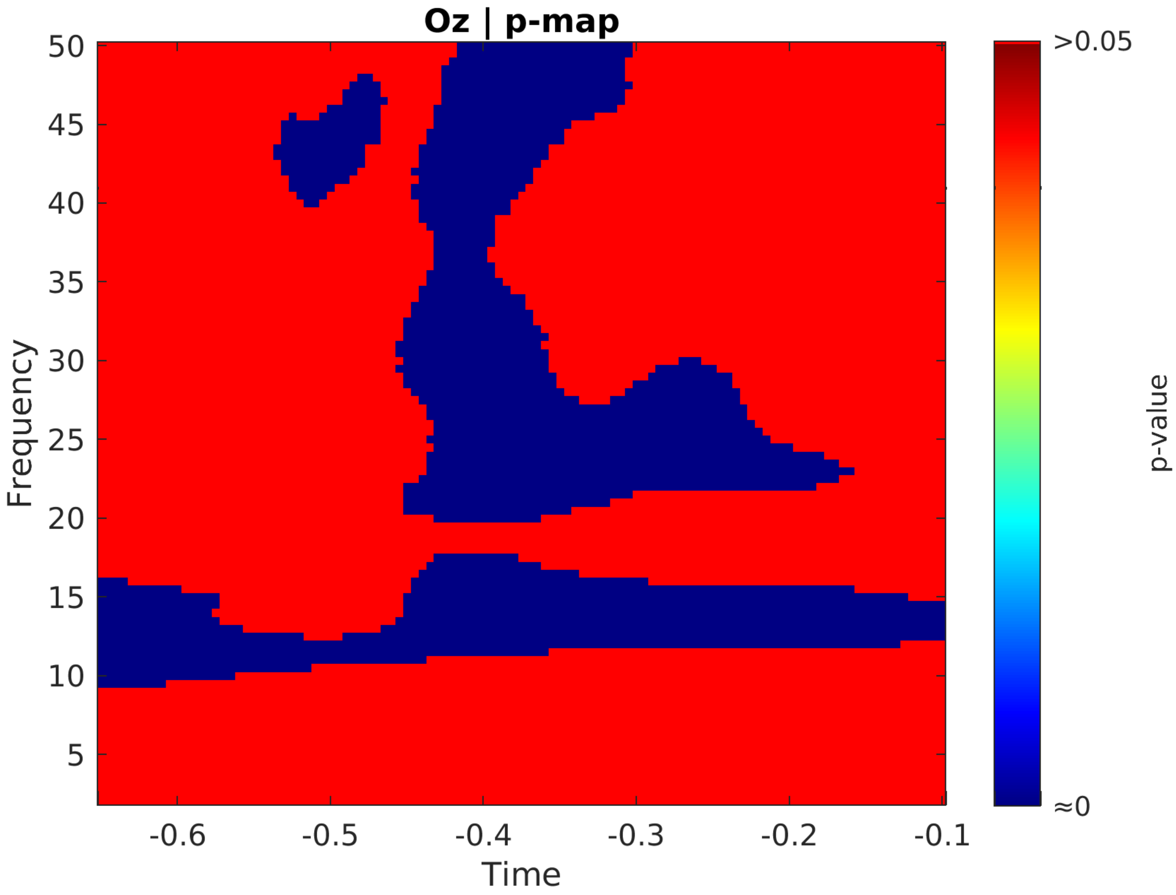
Oz | p-map
50
45
40
35
30
25
20
15
10
5
Frequency
-0.6
-0.5
-0.4
-0.3
-0.2
-0.1
Time
>0.05
≈0
p-value

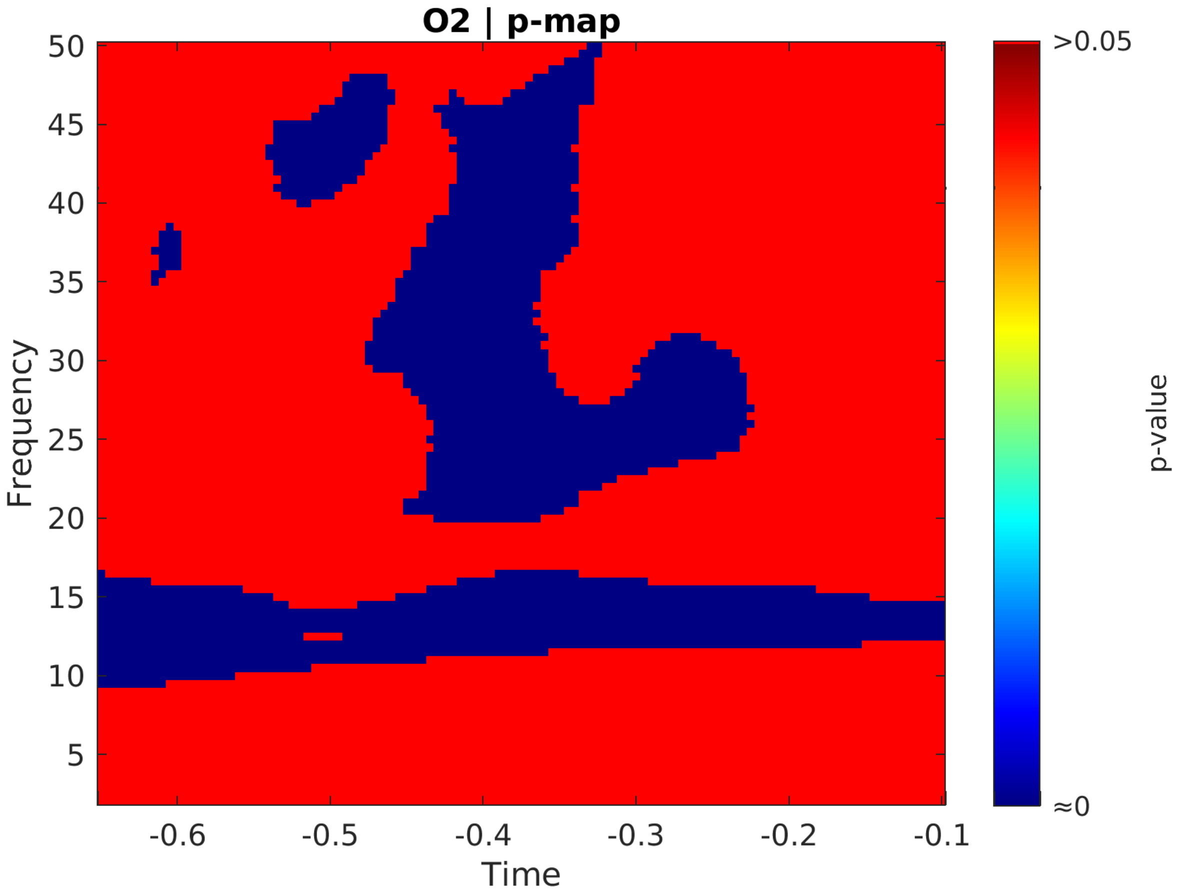
O2 | p-map
50
45
40
35
30
25
20
15
10
5
Frequency
-0.6
-0.5
-0.4
-0.3
-0.2
-0.1
Time
>0.05
≈0
p-value

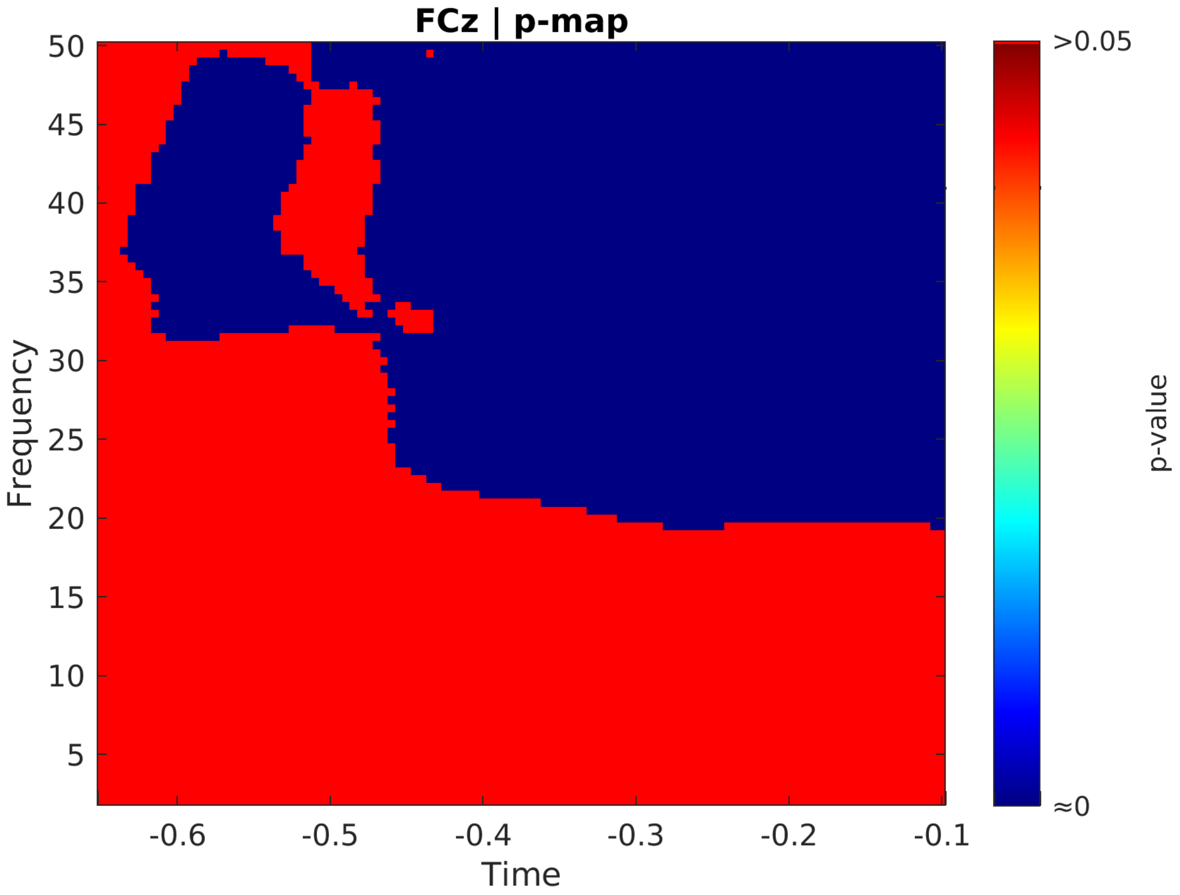

FCz | p-map
Frequency
Time
50
45
40
35
30
25
20
15
10
5
-0.6
-0.5
-0.4
-0.3
-0.2
-0.1
>0.05
≈0
p-value
